# Polymeric assembly of endogenous Tuberous Sclerosis Protein Complex

**DOI:** 10.1101/2021.01.18.426261

**Authors:** David L. Dai, S. M. Naimul Hasan, Geoffrey Woollard, Stephanie A. Bueler, Jean-Philippe Julien, John L Rubinstein, Mohammad T. Mazhab-Jafari

## Abstract

Tuberous Sclerosis protein complex (pTSC) nucleates a proteinaceous signaling hub that integrates information about the internal and external energy status of the cell in regulation of growth and energy consumption. Biochemical and electron cryomicroscopy (cryoEM) studies of recombinant pTSC have revealed the structure and stoichiometry of the pTSC and have hinted at the possibility that the complex form large oligomers. Here, we have partially purified endogenous pTSC from fasted mammalian brains of rat and pig by leveraging a recombinant antigen binding fragment (F_ab_) specific for the TSC2 subunit of pTSC. We demonstrate F_ab_ dependent purification of pTSC from membrane solubilized fractions of the brain homogenates. Negative stain electron microscopy of the samples purified from pig brain demonstrates rod-shaped protein particles with a width of 10 nm, a variable length as small as 40 nm and a high degree of conformational flexibility. Larger filaments are evident with a similar 10 nm width and up to 1 μm in length in linear and web-like organizations prepared from pig brain. These observations suggest polymerization of endogenous pTSC into filamentous super-structures.

## Introduction

pTSC is an intra-cellular stress-sensor that inactivates mTORC1 on lysosomal membranes in response to a multitude of internal and external stimuli (*e*.*g*. energy stress, hypoxia, cytokines, pH variations, and mechanical stress)[1]. The pTSC core complex is a heterotrimer composed of three polypeptides[2]: a 130kDa TSC1 (aka. Hamartin), a 200kDa TSC2 (aka. Tuberin), and a 35kDa accessory protein TBC1D7 that stabilizes the interaction between TSC1 and TSC2. The regulatory role of pTSC on mTORC1 is relayed by the function of the GTPase activating protein (GAP) domain of TSC2 that accelerates GTP hydrolysis by Rheb GTPase[3,4], the primary activator of mTORC1[5]. pTSC is regulated by an extensive network of kinases such as Akt, GSK, and AMPK[1] that phosphorylate the complex. Inactivating mutations in *TSC1* and *TSC2* genes result in TSC disease, manifested by multiple benign tumors in different organ systems, developmental retardation, cysts, lymphangioleiomyomatosis (LAM), and kidney failure[6].

Two recent studies have reported near-complete cryoEM density maps of recombinant human pTSC at near-atomic resolutions[7,8]. Both studies report a rod-shaped architecture that is highly flexible. Each recombinant pTSC assembly is composed of two TSC1, two TSC2, and one TBC1D7 protein. The complex is approximately 10 nm in width and 40 nm in length. To enable high-resolution cryoEM, the authors either stabilized the protein with chemical cross-linking[9] or protected the protein from the air-water interface during cryoEM grid preparation by applying the purified sample on graphene oxide-coated grids[10]. Additionally, both groups utilized focused refinement of the semi-rigid segments of the complex to enable side chain densities to be resolved.

We also observed a peculiar thread-shaped organization of recombinant pTSC (Fig. 1), prompting us to investigate the architecture of the complex from native sources. Signaling protein complexes may form higher order oligomers when expressed under native condition within the host organism as recently observed for fungal TORC1 helical oligomers under stress condition[11]. The higher-order structures of the signaling complexes may require specific post-translational modifications induced via signaling events or interaction with adapter proteins. In mammals, pTSC is naturally abundant in the cerebral cortex[12], we therefore attempted to purify the complex from rat (*Rattus norvegicus*) and pig (*Sus scrofa domesticus*) brain to investigate the overall architecture of the mammalian pTSC by negative stain electron microscopy (NSEM).

**Figure 1.**
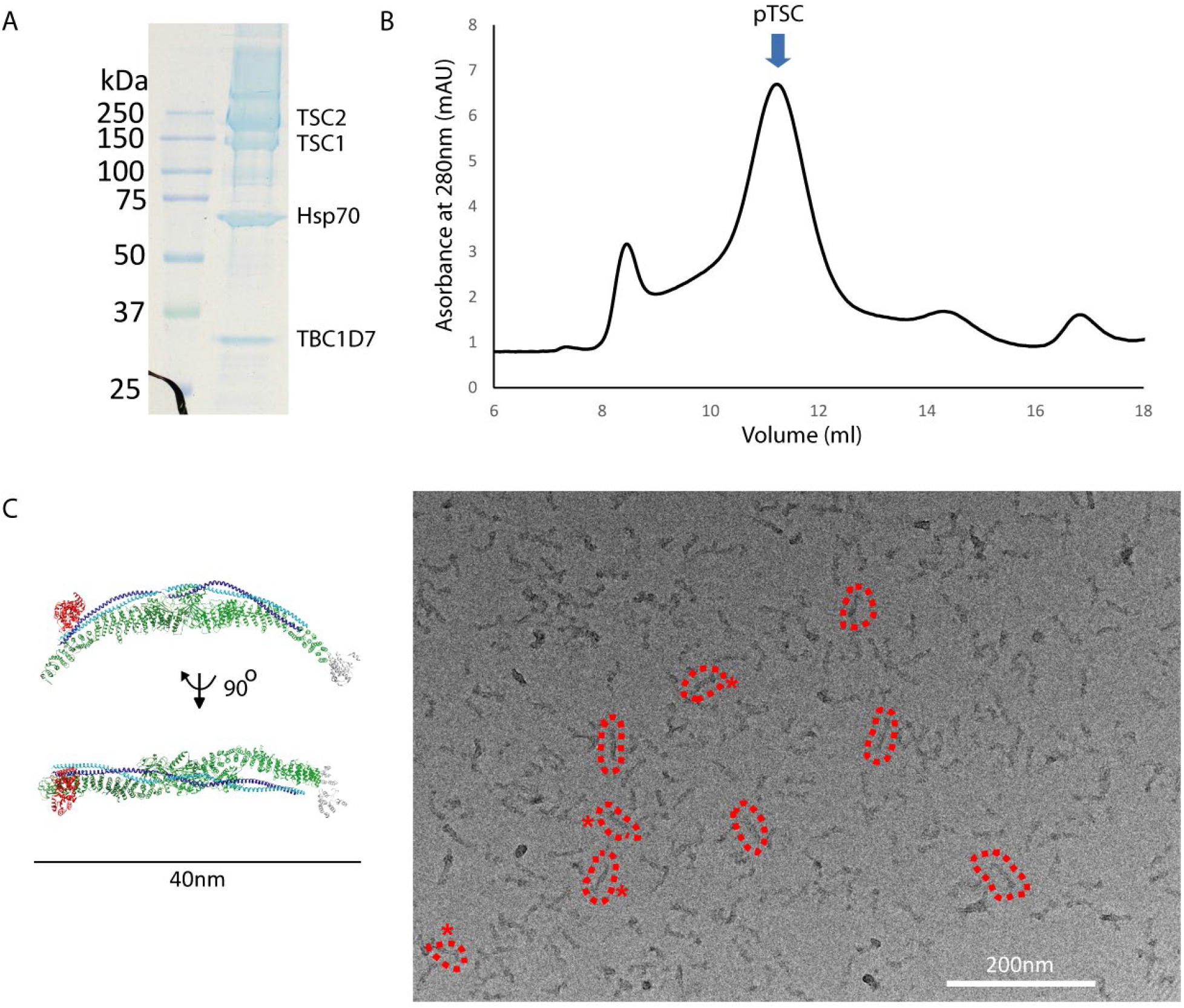
Overall architecture of recombinant human pTSC. **A)** SDS-PAGE analysis of purified protein complex prior to **B)** size exclusion chromatography analysis. **C)** representative NSEM micrograph of recombinant pTSC. Individual pTSC particles highlighted with dashed red circles. The fraction from size exclusion chromatography used for NSEM is highlighted with an arrow in panel B. The atomic model of human pTSC is show to the left (PDB: 7DL1[8]).

## Results

### Recombinant human pTSC forms flexible 40 nm filamentous structures

Consistent with results from two other groups[7,8], we detected ∼10 × 40 nm highly flexible particles in NSEM analysis of human recombinant pTSC overexpressed under the control of CMV promoters in human cells (Fig. 1). A characteristic ‘bump’ was observed in some of the particles at one end, off-axis relative to the long-axis of pTSC, that is at the position of the 35kDa TBC1D7 protein (Fig. 1C). The observation of this asymmetric feature is dependent on the viewing direction of the particles and not all particles show this feature.

We found that recombinant human pTSC co-purifies with heat shock protein 70 (Hsp70), as observed in other studies[7,13]. However, no density could be found for the heat shock protein in the recent cryoEM maps indicating that the heat shock proteins most likely binds misassembled or partially unfolded pTSC.

### Endogenous pTSC from mammalian brain exhibits heterogenous morphology

In order to purify endogenous pTSC, a recombinant F_ab_ construct was designed from an anti-TSC2 monoclonal antibody (mAb) that recognizes the C-terminal region of tuberin and has been shown to immunoprecipitate pTSC from human cells[14]. We tested the ability of the antibody to recognize the unfolded TSC2 from rat and pig brain fractions and confirmed specific recognition of the epitopes by the mAb in both species (Fig. 2A and B). The amino acid sequence of the antibody variable region was then determined through mass spectrometry analysis of the proteolytic fragments. Heavy and light chains encompassing a F_ab_ fragment were cloned into mammalian expression vectors and expressed in HEK293F cells. The F_ab_ harboured a calmodulin binding peptide (CBP) and a 3×FLAG peptide on the C-terminus of the light and heavy chains, respectively. The CBP-tag was used to purify the F_ab_. The purified F_ab_ (Fig. 2C) was immobilized on an anti 3×FLAG column and used to immunopurify pTSC from native sources. pTSC-F_ab_ complex was then competitively eluted via excess 3×FLAG peptide. Rat and pig brains were chosen due to ease of access (for rat brain), high quantity of starting native material (∼70 grams brain weight for pig), and high sequence identity of pTSC components to the human complex (85% to 92% identity for TSC1, TSC2, and TBC1D7 between the largest polypeptide isoforms of the two species to that of the human proteins, see supplementary data 1-6).

**Figure 2.**
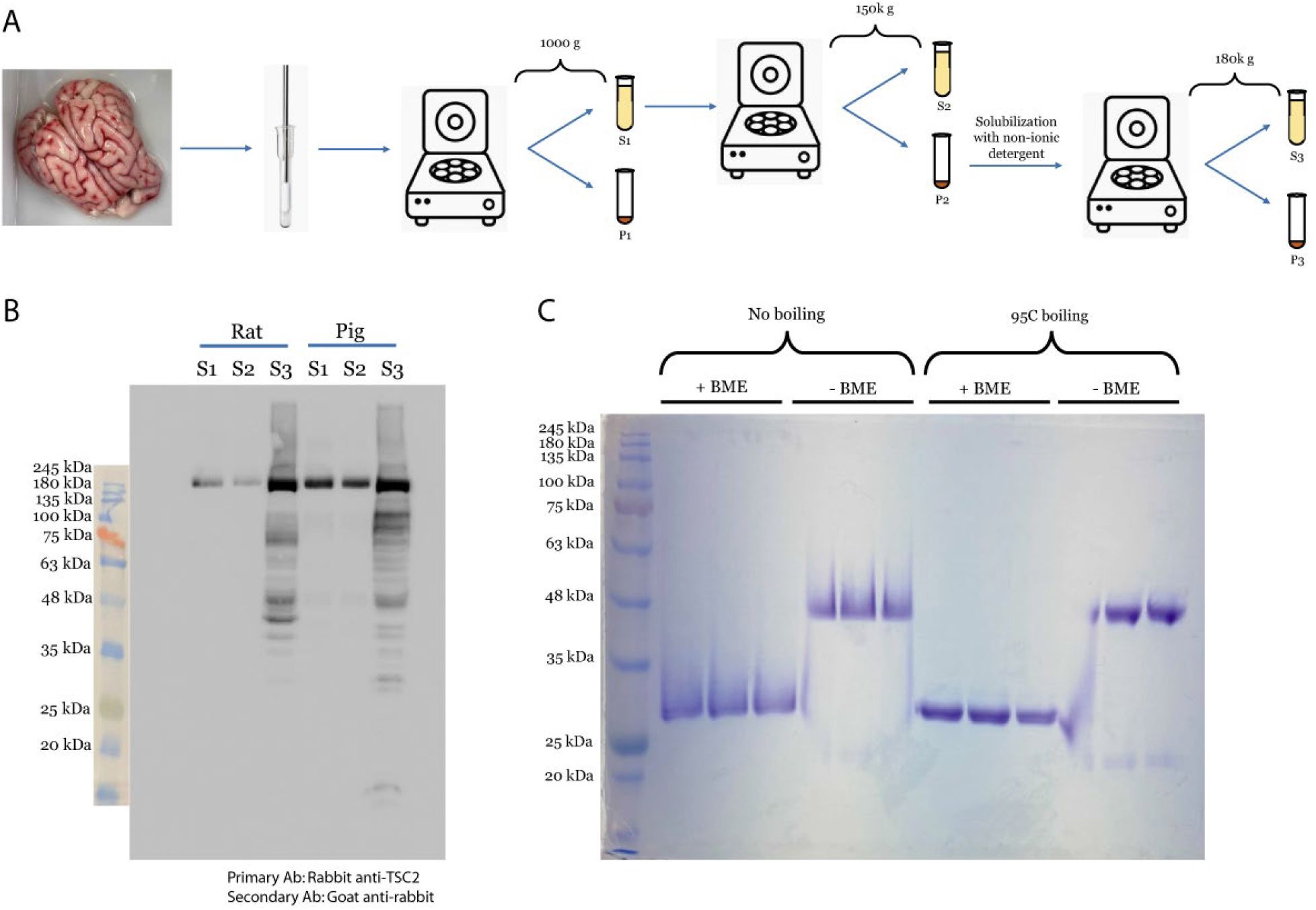
Detection of TSC2 in rat and pig membrane solubilized brain fractions. **A)** Schematic for preparation of membrane solubilized fraction of rat and pig brains. Pig brain is shown as an example. **B)** Immunoblot of the indicated fractions using anti-TSC2 antibody. **C)** SDS-PAGE analysis of the purified recombinant F_ab_ in the presence and absence of reducing agent and sample boiling, demonstrating cysteine-dependent heterodimerization of the light and heavy chains of F_ab_.

Low ATP levels are a well-known stimulatory factor of pTSC[15]. Therefore animals were fasted prior to sacrifice. TSC2 subcellular localization (Fig. 2A) was assessed by western blot on soluble (S2) and membrane fractions (S3) of the brain homogenate. n-Dodecyl β-D-maltoside (DDM) was used to solubilize the rat and pig P2 membrane pellet (Fig. 2A). Both S2 and S3 fractions contained TSC2 (Fig. 2B). However, we proceeded with the membrane fraction (S3) because active pTSC is co-localized with farnesylated Rheb on the outer lysosomal membrane[14,16].

The recombinant F_ab_ enabled purification of TSC2 from membrane fractions of rat and pig brain homogenate as observed via immunoblot analysis using the rabbit monoclonal antibody against tuberin from which the F_ab_ used in purification was derived (Fig. 3A and 4A). TSC2 in both species appeared with a molecular weight of approximately 240kDa. The estimated mass of TSC2 from amino acid sequence in both species is approximately 200kDa, indicating potential post-translational modifications of the endogenous proteins. The 35kDa TBC1D7 was also purified in a F_ab_-dependent manner observed via immunoblotting using a rabbit monoclonal antibody against the protein (Fig. 3C and 4C). Both TSC2 and TBC1D7 were detected with a clear western blot signal in the S3 fraction before and after passage through the F_ab_-column, indicating that the F_ab_ was capturing only a small fraction of the complexes from S3. Presence of the TBC1D7 in the elution fractions indicates the presence of TSC1 because the small 35 kDa accessory protein interacts directly with TSC1 in pTSC and not TSC2 as observed in the cryoEM structure of the human complex[7,8]. This mode of interaction is supported by X-ray crystallographic studies of truncated TSC1 with TBC1D7[13,17]. Direct association of TBC1D7 with TSC1 is also supported by immunoprecipitation studies showing that the 35kDa accessory protein binds TSC1 but not TSC2[8]. TSC1 was detected at approximately 180kDa in the rat elution fractions using a rabbit monoclonal antibody that binds a synthetic peptide from human TSC1 around residue Val640 (Fig. 3B). However, TSC1 was not detected in the elution fractions with and without F_ab_ in the pig preparation using the same antibody and was only weakly detected in the S3 fraction at approximately 180 kDa, as observed in rat (Fig. 4B, left panel). Residues surrounding Val640 in the middle region of human TSC1 are less well conserved in pig than rat species (supplementary data 2 and 5), which could explain the weaker affinity of the primary antibody to the TSC1 peptide in pig. Analysis of the F_ab_ elution fractions from pig with a mouse monoclonal antibody against the N-terminal region of TSC1 showed the presence of a weak 180kDa protein species corresponding to the similar mass found for TSC1 in rat that could be indicative of post-translational modifications (Fig. 4B middle panel). It also detected a signal at approximately 130kDa in the pig elution samples corresponding to the theoretical mass of TSC1 in this species (Fig. 4B middle panel). The strong signal at 25 kDa and 60 kDa are likely breakdown products of TSC1. Higher detergent (*i*.*e*. 0.5% w/v DDM) concentration used in solubilization of pig P2 membrane pellet resulted in stronger relative signal at 130kDa for TSC1 (Fig. 4B, right panel). Overall, these data indicate immuno-purification of minute quantities of pTSC from rat and pig membrane-solubilized brain fractions. To analyse the overall architecture and organization of endogenous pTSC, we proceeded with NSEM imaging of F_ab_ elution fractions (henceforth referred to as ‘plus F_ab_’ preparation). Elution fractions from the purification with affinity column devoid of F_ab_ (henceforth referred to as ‘minus F_ab_’ preparation) were used as negative control in NSEM analysis.

**Figure 3.**
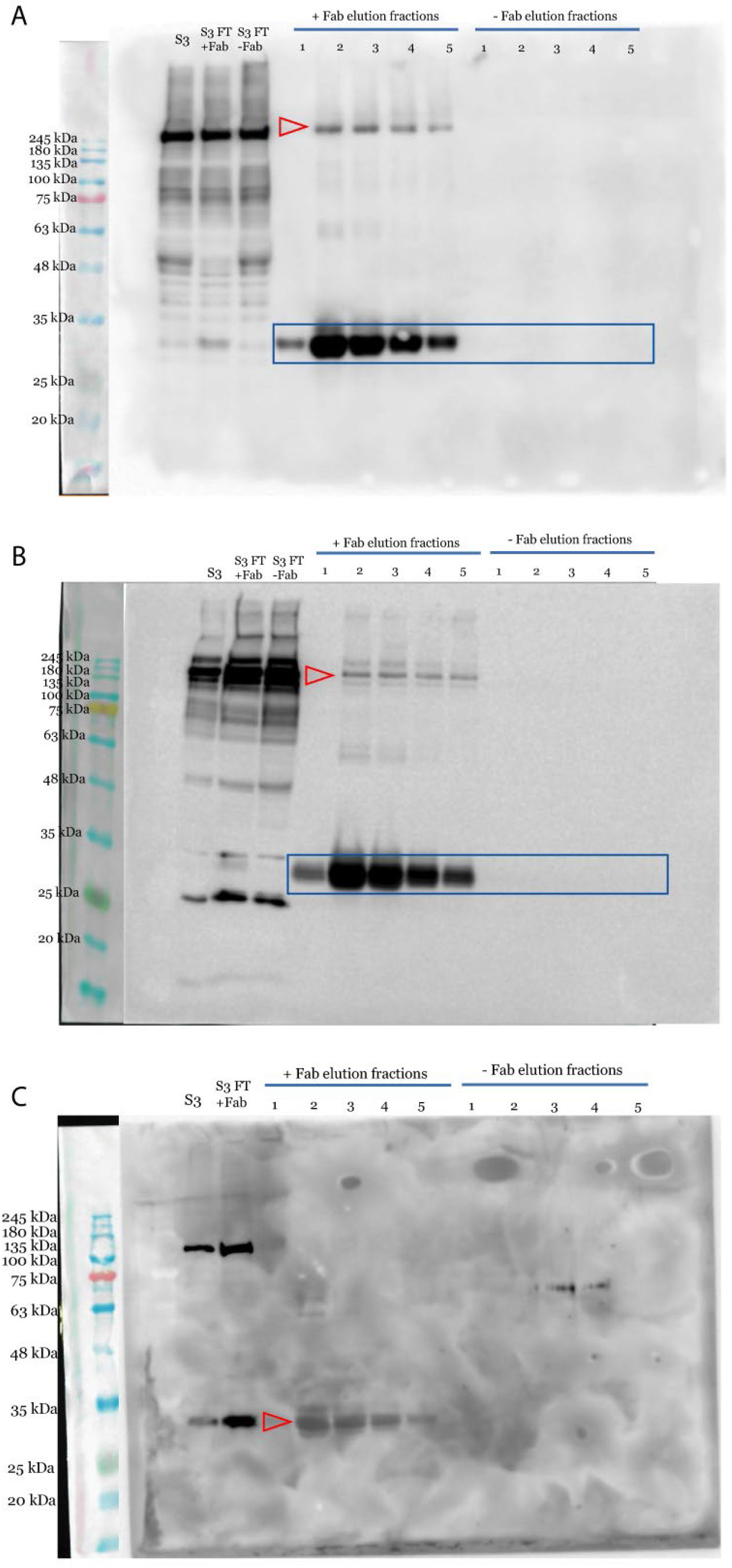
Purification of pTSC from rat membrane solubilized brain fractions. **A)** Western blot analysis of S3 and elution fractions using rabbit mAbs against TSC2 (mAb), **B)** TSC1 (mAb), and **C)** TBC1D7 (mAb). The position of the pTSC subunits are shown with a red arrowhead in each panel. The strong band at 30kDa highlighted by cyan rectangles in panels A and B are a reaction of the goat anti-rabbit secondary antibody with the rabbit-derived F_ab_’s heavy and light chains. A F_c_-specific secondary antibody (mAb) was used in panel C to probe for TBC1D7 at 35kDa.

**Figure 4.**
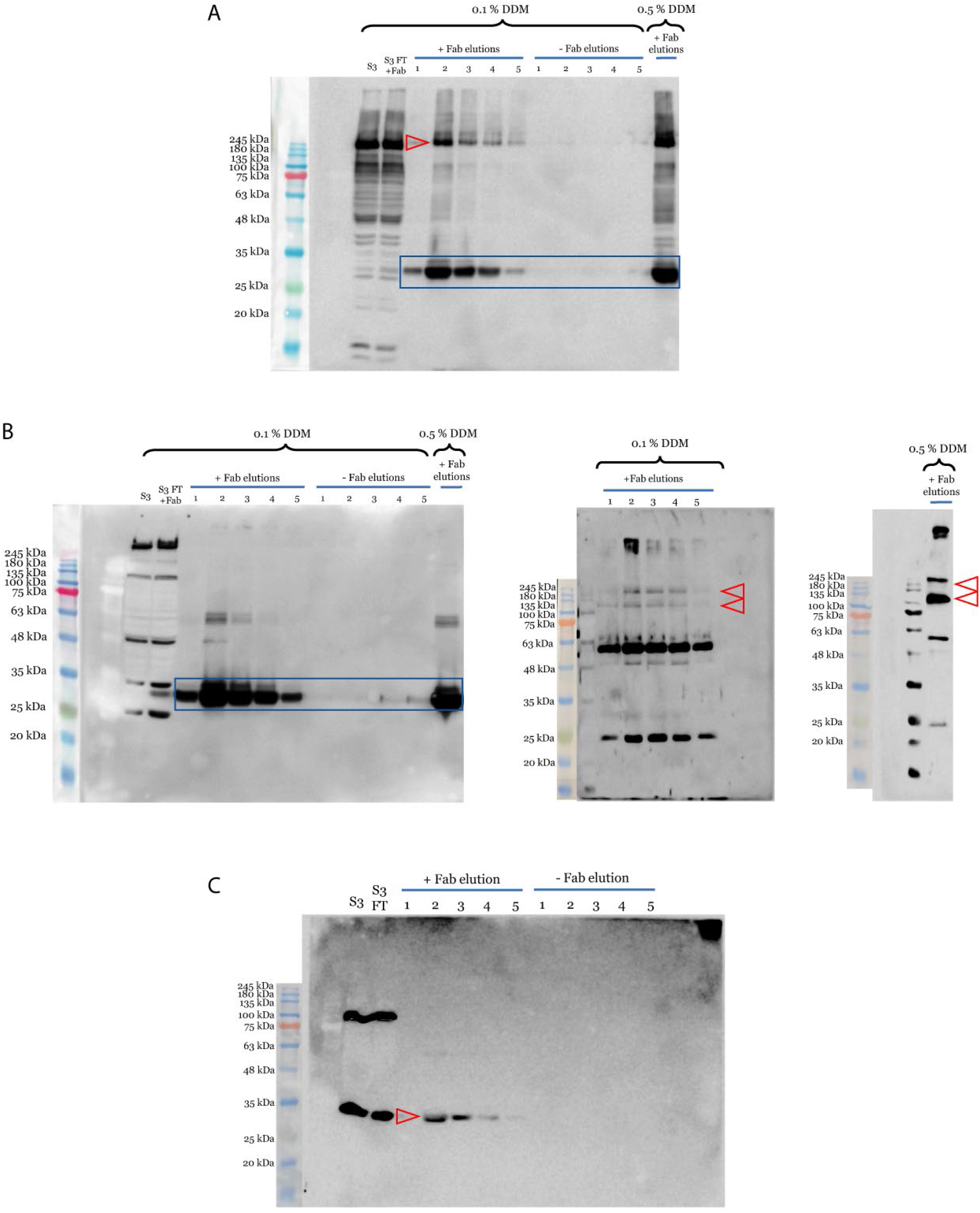
Purification of pTSC from pig membrane solubilized brain fractions. **A)** Western blot analysis of S3 and elution fractions using a rabbit mAb against TSC2 mAb, **B)** TSC1 (rabbit anti-TSC1 mAb left, mouse anti-TSC1 middle and right), and **C)** TBC1D7 (rabbit mAb against TBC1D7 with a secondary antibody specific to the F_c_ region of primary mAb). The red arrowheads and cyan rectangles are as described in figure 3.

NSEM analysis of rat elution fractions demonstrated highly heterogenous particles in both plus and minus F_ab_ samples (supplementary data 7). Observation of protein species in the minus F_ab_ purification is likely due to non-specific binding of the S3 lysate from rat to the anti-FLAG tag matrix. Therefore, it was not possible to distinguish any specific molecular morphology between the positive and negative control samples. Interestingly, however, long and highly flexible structures were observed in elution fractions of pig in the plus F_ab_ preparation that were absent in the minus F_ab_ preparation (Fig. 5A, supplementary data 8). The minus F_ab_ preparation from pig demonstrated significantly fewer total proteins compared to the plus F_ab_ preparation in the NSEM images indicating that most protein complexes observed were directly captured by the anti-TSC2 F_ab_ from pig S3 lysate. pTSC purification from a second pig brain demonstrated similar NSEM images with long and flexible filaments (Fig. 5B, supplementary data 9). These particles represented a filamentous overall shape with 10 nm width and length varying from ∼40 nm up to 1 μm. Small aggregates of the filaments were also observed in a web-like fashion. Other non-filamentous structures were observed as well, which could represent non-pTSC proteins or endogenous pTSC in potentially alternate conformations. Whether these polymeric structures are composed exclusively of pTSC or additional proteins are associated to stabilize or regulate their formation in response to low energy status is still unknown and requires further investigations.

**Figure 5.**
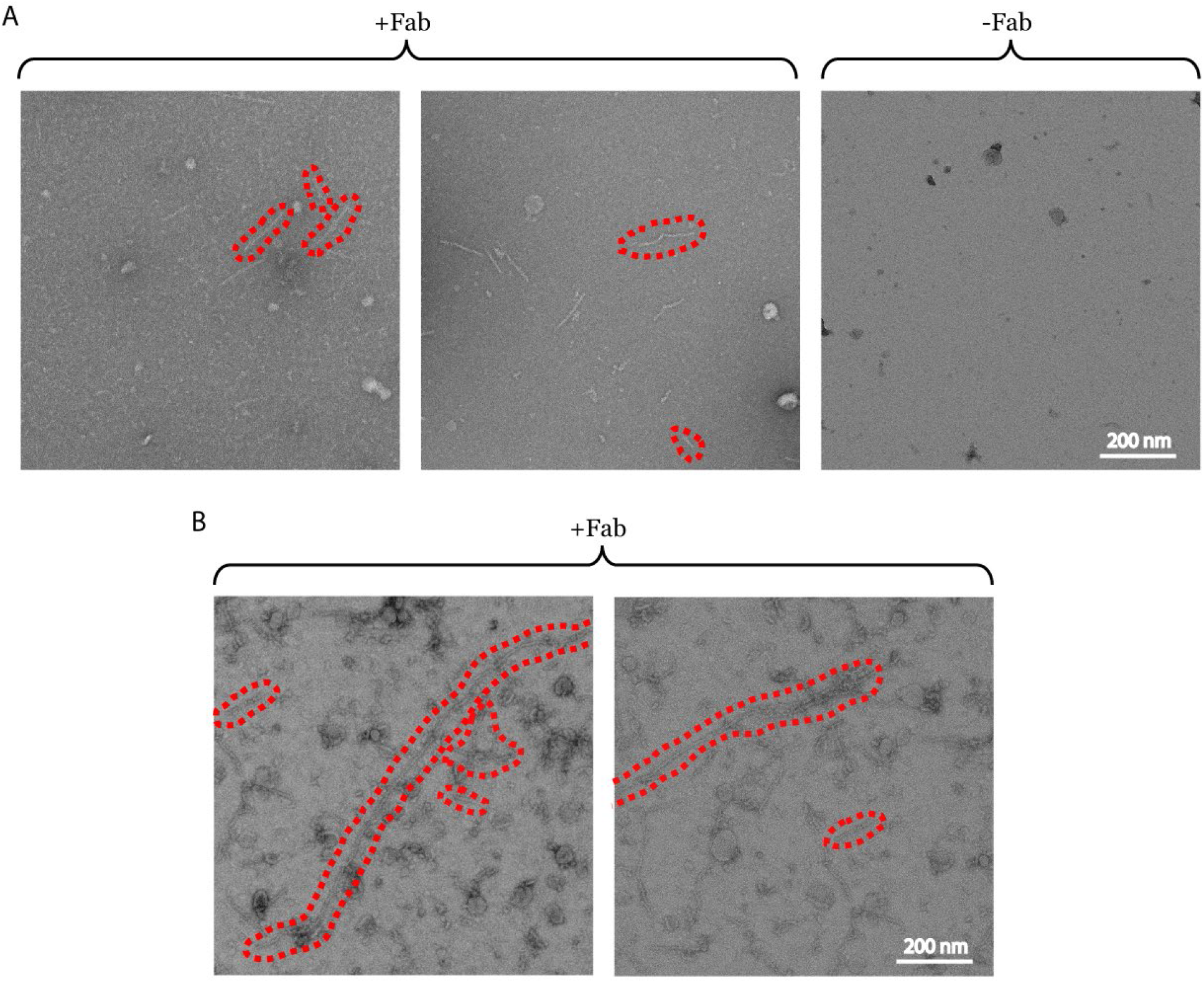
Observation of flexible filaments purified using anti-TSC2 Fab from pig brain. **A)** Two representative micrographs of pTSC purified from pig brain are shown to the left and center. Long and flexible filaments are highlighted with dashed lines. A representative micrograph for minus F_ab_ purification is shown to the right. **B)** Two representative micrographs from pTSC purified from a second pig brain as described in the text. All micrographs are to the same scale.

## Discussion

Hamartin and tuberin are highly regulated proteins in cells with diverse functions. The best studied role of the two proteins is in the context of mTORC1 signaling axis in regulation of mRNA translation and autophagy. However, both proteins are involved in non-mTORC1 (*i*.*e*. non-canonical) signaling and interact with a multitude of partner proteins to regulate diverse functions such as apoptosis, cell fate and differentiation, actin cytoskeletal reorganization, regulation of aggresome, and secretion across cellular membranes[18]. Considering presence of multiple isoforms of both Tuberin and Hamartin, it is not surprizing that immunoprecipitation of endogenous pTSC from human cell lines leads to detection of a multitude of interacting protein partners[2]. Therefore, pTSC may exist in multiple conformational, chemical, and oligomeric states[19] within cells as a ‘versatile signaling complex’.

Our novel purification strategy for pTSC demonstrates the extent of complex morphology of an endogenous mammalian pTSC where protein particles with multiple overall shapes and sizes are seen in NSEM images. Particularly, the long and flexible filamentous structures are suggestive of pTSC polymerization. These assemblies may be highly labile and prone to disassembly upon biochemical purification, which could explain their absence in rat samples. It is also possible that their formation is differentially regulated in rat and pig species upon fasting of the animals. The recent cryoEM models of recombinant human pTSC suggests a head-to-tail oligomerization potential[7]. This hypothesis was supported by a concentration dependent observation of a head-to-tail association of individual pTSC complexes in the cryoEM micrographs and shifting of the elution profile of the recombinant protein toward large molecular weights by increasing concentration[7]. As a signaling complex it may be possible that pTSC requires specific post-translational modifications and/or association with adaptor proteins to form the observed ultra-structural organizations. It may also be possible that inter-subunit interactions between TSC1, TSC2, and TBC1D7 may adopt minor structural rearrangements to accommodate linear and branched (*i*.*e*. web-like) polymerization. A high-resolution structure of the observed polymers from pig purified samples may be the key to answer some of these intriguing questions and may enable structural-based functional interrogation of the supra-molecular organization of pTSC.

### Experimental procedures

#### Recombinant pTSC

TBC1D7 (pcDNA3.1/nMyc-DEST) was purchased from the SPARC BioCentre at the SickKids Research Institute (Toronto). TSC1 (pcDNA2) and TSC2 (pcDNA3.1) were provided by the Vuk Stambolic laboratory at the University Health Network (Toronto). 3×-FLAG was inserted at the N-terminal end of TSC2. Asn1643 of TSC2 was mutated to Glu to abolish the GAP activity of pTSC. The inactive pTSC resulted in a higher protein yield since active pTSC inhibits protein translation and cellular growth.

A 150ml HEK293F culture was transfected with 37.5 μg of each TSC1, 3×-FLAG-TSC2 (N1643Q), and TBC1D7 using FectoPRO transfection agent at a ratio of 1 μL per 1 μg of DNA. Cell density was at 0.8 × 10^6^ cells/mL with 99% viability before transfection. Cells were harvested 48 post-transfection and stored at −80°C until use. Cells were lysed via incubation with lysis buffer (50 mM HEPES, pH 7.5, 100 mM NaCl, 50 mM β-glycerphosphate, 10 mM NaF, 10% v/v glycerol, 0.5 mM PMSF, 1 mM Benzamidine, 1 mM 6-aminocaproic acid, and 0.1% v/v Triton x100) for 10min on ice with occasional gentle agitation. Lysate was cleared at 12,000 g for 30 min and supernatant was filtered and passed through an anti-FLAG M2-affinity column. The column was washed with 50 mM HEPES pH 7.5, 0.5 M NaCl followed by a second wash buffer containing 50 mM HEPES pH 7.5, 100 mM NaCl, 10 mM ATP, and 20 mM MgCl_2_. The ATP wash step was necessary to remove Hsp90 that is routinely observed to co-purify with recombinant pTSC. The protein was then eluted with 150 ng/μL of synthetic 3×-FLAG peptide followed by size exclusion chromatography using a Superose 6 column.

#### Anti-TSC2 F_ab_ expression plasmids

Antibody protein sequencing by mass spectrometry was performed on a rabbit anti-TSC2 antibody (Cell Signaling Technology, Clone #D93F12, Lot #2) by Rapid Novor Inc (www.rapidnovor.com) to obtain the complete amino acid sequence of the antibody variable regions. The sequences were used to synthesize two pcDNA 3.4 expression vectors encoding the anti-TSC2 F_ab_ light and heavy chains (GeneArt (Thermofisher Scientific)). These expression vectors were synthesized to include a Sec-translocase signal peptide at the N-termini of both anti-TSC2 F_ab_ chains, a 3xFLAG tag at the C terminus of the anti-TSC2 F_ab_ heavy chain and a calmodulin binding peptide tag at the C terminus of the anti-TSC2 F_ab_ light chain.

#### Expression and purification of an anti-TSC2 F_ab_ from HEK293F cells

A 25 mL HEK293F starter cell culture in Gibco FreeStyle media was grown in a New Brunswick S41i shaking incubator at 37°C, 8% CO_2_ and 125 RPM until it reached a total cell density of 3.0 × 10^6^ cells/mL and a viability of at least 95%. The starter culture was then used to seed a 22.5 mL transfection culture to a total cell density of × 10^6^ cells/mL. The transfection culture was then grown in New Brunswick S41i shaking incubator at 37°C, 8% CO_2_ and 125 RPM for 5 hours until it reached a total cell density of 1.11 × 10^6^ cells/mL. The transfection culture was then mixed with a filtered 2.5 mL transfection solution in Gibco FreeStyle media composed of 6.25 µg anti-TSC2 F_ab_ light chain DNA, 6.25 µg F_ab_ heavy chain DNA and 12.5 µL FectoPRO (Polyplus Transfection, Ref #166-001, Lot #12W2808FCI). The transfection culture was grown for 96 hours in a New Brunswick S41i shaking incubator at 37°C, 8% CO_2_ and 125 RPM. After 96 hours, the transfection culture was decanted into a 50 mL Sarstedt Falcon tube then centrifuged at 3000 g for 10 minutes at 4°C using a SX4400 Beckman Coulter rotor in a Beckman Coulter Allegra X-30R centrifuge. The transfection culture supernatant was collected and syringe filtered through a Millex 0.45 µm low protein binding Durapore. Anti-TSC2 F_ab_ was purified from this supernatant by gravity filtration using 1 mL calmodulin-agarose beads (Sigma-Aldrich, Product #A6112) packed in a Bio-Rad poly-prep chromatography column pre-equilibrated in binding buffer (50 mM Tris, 100 mM NaCl, 2 mM CaCl_2_, pH 7.4). The column was then washed with 5 column volumes of binding buffer before eluting with five 1 column volume loads of elution buffer (50 mM Tris, 100 mM NaCl, 10 mM EGTA, pH 7.4). The elution fractions were then pooled and concentrated to 1 mL using an Amicon Ultra-15 centrifugal filter unit with 10 kDa cut-off before passing the concentrated eluate through a GE Superose 6 increase 10/300 GL size exclusion column pre-equilibrated with size exclusion buffer (50 mM Tris, 100 mM NaCl, pH 7.4). Size exclusion elution fractions containing highly purified anti-TSC2 F_ab_ were pooled and saved.

#### Validating the purification of anti-TSC2 F_ab_ by +/- β-mercaptoethanol SDS-PAGE assay

Starting with twelve 20 µL aliquots of purified anti-TSC2 F_ab_, one half were mixed with 5 µL 5X Laemmeli buffer with β-mercaptoethanol (200 mM Tris pH 6.8, 45% v/v glycerol, 5% w/v SDS, 0.0125% w/v Bromophenol Blue, 1.8M β-mercaptoethanol) while the other half were mixed with 5 µL 5X Laemmeli buffer without β-mercaptoethanol (200 mM Tris pH 6.8, 45% v/v glycerol, 5% w/v SDS, 0.0125% w/v Bromophenol Blue). The samples were incubated at room temperature for 5 minutes before half of the samples with and without β-mercaptoethanol were boiled for 2 minutes in a Mandel DryBath Plus heat block at 95°C while the other halves were not boiled. All samples were then run on a 10% SDS-gel in running buffer (3.5 mM SDS, 25 mM Tris, 200 mM glycine, pH 8.3) for 60 minutes at 200V. The SDS-PAGE gel was then submerged in Coomassie blue stain (4% v/v methanol, 7% v/v acetic acid, 0.1% w/v Coomassie blue, 88.9% v/v MilliQ H_2_O) and microwaved for 1 minute before incubating in the stain for 1 hour at room temperature on a Bio-Rad Ultra Rocker shaker. The SDS-PAGE gel was then submerged in de-staining solution (45% v/v methanol, 10% v/v acetic acid, 55% v/v MilliQ H_2_O) and microwaved for 1 minute before incubating in the de-stain for 3 hours at room temperature on a Bio-Rad Ultra Rocker shaker. The SDS-gel was then submerged in distilled water and incubated overnight at room temperature on a Bio-Rad Ultra Rocker shaker before imaging.

#### Purification of pTSC from pig brain tissue

One entire male pig brain, provided by the PMCRT animal facility, was harvested from an 11-week old Yorkshire breed blood donor pig that had been fasted for 6 hours prior to its euthanization with an overdose of KCl via intravenous injection. The brain was then placed inside a container of ice-cold Tris-buffer saline (50 mM Tris, 100 mM NaCl, pH 7.4) then processed by slicing off meninges, fatty tissue, connective tissue and any other rubbery non-brain tissue that would not homogenize easily. The processed pig brain was resuspended in 2 mL/g lysis buffer (4 mM HEPES pH 7.4, 320 mM sucrose, 1.0 mM PMSF, 50 mM β-glycerophosphate, 1 mM NaF, 0.3 mM sodium orthovanadate, 5 mM aminocaproic acid, 5 mM benzamide, 5 mM EDTA) supplemented with 0.0028% mammalian protease inhibitor cocktail (Sigma-Aldrich, Product #P8340, MDL #MFCD00677817) and then was grinded inside a 30 mL Thomas glass grinding vessel with Teflon pestle using a Janke & Kunkel RW 20 DZM homogenizer set at 555 RPM. The resulting homogenate was centrifuged at 1000 ×g for 10 minutes at 4°C using a Beckman Coulter SX4400 rotor in a Beckman Coulter Allegra X-30R centrifuge. The pellet fraction (P1) containing nuclei, broken cells and whole cell debris was discarded and the purification proceeded by isolating the supernatant fraction (S1) and ultra-centrifuging it in a Beckman Coulter Type 70 Ti rotor with a Beckman Coulter Optima L-100 XP ultracentrifuge for 45 minutes at 150,000 ×g and 4°C. The resulting supernatant fraction (S2) containing the population of all soluble proteins inside the cell was saved while the purification proceeded with the pellet fraction (P2) containing the population of all membrane proteins in the cell including the active form of pTSC peripherally associated to the outer lysosomal membrane. P2 was resuspended in 30 mL lysis buffer by grinding it inside a 30 mL Thomas glass grinding vessel with Teflon pestle using a Janke & Kunkel RW 20 DZM homogenizer set at 555 RPM. 0.5% or 0.1% N-dodecyl maltoside was then added to the P2 resuspension to solubilize the membranes as the resuspension stirred gently for 30 minutes in 4°C. Finally, the solubilized P2 fraction was ultra-centrifuged in a Beckman Coulter Type 70 Ti rotor with a Beckman Coulter Optima L-100 XP ultracentrifuge for 65 minutes at 180,000 ×g and 4°C to obtain a supernatant fraction (S3) enriched with all solubilized membrane proteins. S3 was then filtered through a Millex 0.45 µm low protein binding Durapore. pTSC was then purified from S3 via gravity filtration by passing S3 through either a +F_ab_ or −F_ab_ Bio-Rad poly-prep chromatography column composed of 250 µL M2 agarose beads (Sigma-Aldrich, Lot #SLCF2052) pre-equilibrated in Tris-buffer saline and that had or had not been pre-bound to purified anti-TSC2 F_ab_ by its 3x-FLAG tag, respectively. Both +F_ab_ and −F_ab_ columns were washed with 5 column volumes of Tris-buffer saline before eluting with five 1 column volume loads of elution buffer (50 mM Tris, 100 mM NaCl, 150 µg/mL 3x-FLAG peptide) to obtain +F_ab_ elution fractions containing purified pig pTSC or −F_ab_ elution fractions not containing purified pig pTSC.

#### Purification of pTSC from rat brain tissue

An entire male rat brain, provided by the PMCRT animal facility, was harvested from a 6-week old Sprague-Dawley strain rat that had been fasted for 16 hours prior to its euthanization by CO_2_. The brain was then submerged in 18 mL/g lysis buffer (4 mM HEPES pH 7.4, 320 mM sucrose, 1.0 mM PMSF, 50 mM β-glycerophosphate, 1 mM NaF, 0.3 mM sodium orthovanadate, 5 mM aminocaproic acid, 5 mM benzamide, 5 mM EDTA) supplemented with 0.0028% mammalian protease inhibitor cocktail (Sigma-Aldrich, Product #P8340, MDL #MFCD00677817) and was then grinded inside a 30 mL Thomas glass grinding vessel with Teflon pestle using a Janke & Kunkel RW 20 DZM homogenizer set at 555 RPM. The resulting homogenate was centrifuged at 1000 ×g for 10 minutes at 4°C using a Beckman Coulter SX4400 rotor in a Beckman Coulter Allegra X-30R centrifuge. The pellet fraction (P1) containing nuclei, broken cells and whole cell debris was discarded and the purification proceeded by isolating the supernatant fraction (S1) and ultra-centrifuging it in a Beckman Coulter Type 70 Ti rotor in a Beckman Coulter Optima L-100 XP ultracentrifuge for 45 minutes at 150,000 ×g and 4°C. The resulting supernatant fraction (S2) containing the population of all soluble proteins inside the cell was saved while the purification proceeded with the pellet fraction (P2) containing the population of all membrane proteins in the cell including the active form of pTSC peripherally associated to the outer lysosomal membrane. P2 was resuspended in 30 mL lysis buffer by grinding it inside a 30 mL Thomas glass grinding vessel with Teflon pestle using a Janke & Kunkel RW 20 DZM homogenizer set at 555 RPM. 0.5% N-dodecyl maltoside was then added to the P2 resuspension to solubilize the membranes as the resuspension stirred gently for 30 minutes in 4°C. Finally, the solubilized P2 fraction was ultra-centrifuged in a Beckman Coulter Type 70 Ti rotor with a Beckman Coulter Optima L-100 XP ultracentrifuge for 65 minutes at 180,000 ×g and 4°C to obtain a supernatant fraction (S3) enriched with all solubilized membrane proteins. S3 wan then filtered through a Millex 0.45 µm low protein binding Durapore. pTSC was then purified from S3 via gravity filtration by passing S3 through either a +F_ab_ or −F_ab_ Bio-Rad poly-prep chromatography column composed of 250 µL M2 agarose beads (Sigma-Aldrich, Lot #SLCF2052) pre-equilibrated in Tris-buffer saline (50 mM Tris, 100 mM NaCl, pH 7.4) and that had or had not been pre-bound to purified anti-TSC2 F_ab_ by its 3x-FLAG tag respectively. Both +F_ab_ and −F_ab_ columns were washed with 5 column volumes of Tris-buffer saline before eluting with five 1 column volume loads of elution buffer (50 mM Tris, 100 mM NaCl, 150 µg/mL 3x-FLAG peptide) to obtain +F_ab_ elution fractions containing purified rat pTSC or −F_ab_ elution fractions not containing purified rat pTSC.

#### Tracking pTSC throughout rat and pig brain subcellular fractions by western blot

1 mL aliquots of subcellular fractions S1, S2 and S3 were saved from both rat and pig brain pTSC purifications. Their protein concentrations were normalized using the ThermoFisher Pierce BCA Protein Assay Kit, leading to fractions S1 and S2 from both rat and pig brains being diluted 1:40 and 1:20 in Tris-buffer saline (50 mM Tris, 100 mM NaCl, pH 7.4) respectively. 20 µL of each normalized fraction were mixed with 5 µL 5X Laemmeli buffer (200 mM Tris pH 6.8, 45% v/v glycerol, 5% w/v SDS, 0.0125% w/v Bromophenol Blue, 1.8M β-mercaptoethanol) then incubated at room temperature for 5 minutes before boiling them in a Mandel DryBath Plus heat block for 2 minutes at 95°C. All samples were then run on a 10% SDS-gel in running buffer (3.5 mM SDS, 25 mM Tris, 200 mM glycine, pH 8.3) for 65 minutes at 200V. The SDS-gel along with an Amersham Biosciences nitrocellulose membrane were then equilibrated in transfer buffer (50 mM Tris, 40 mM Glycine, 1 mM SDS, 10% v/v methanol, pH 8.3) for 5 minutes before being placed in a transfer sandwich submerged in transfer buffer. The proteins on the gel were transferred onto the membrane for 120 minutes at 100V on ice. The membrane was then washed in TBST buffer (50 mM Tris, 100 mM NaCl, 1% v/v Tween 20) for 5 minutes three times before incubating in blocking buffer (50 mM Tris, 100 mM NaCl, 1% v/v Tween 20, 5% w/v skim milk powder) for 1 hour at room temperature. The membrane was washed again in TBST buffer for 5 minutes three times before incubating overnight for 17 hours at 4°C in 10 mL primary antibody solution [50 mM Tris, 100 mM NaCl, 1% v/v Tween 20, 5% w/v skim milk powder, 1:5000 v/v rabbit anti-TSC2 antibody (Cell Signaling Technology, Clone #D93F12, Lot #6)]. The membrane was washed again in TBST buffer for 5 minutes three times then incubated for 1 hour at room temperature in 10 mL secondary antibody solution [50 mM Tris, 100 mM NaCl, 1% v/v Tween 20, 5% w/v skim milk powder, 1:5000 v/v goat anti-rabbit secondary antibody (Cell Signaling Technology, Product #7074P2, Lot #28)]. The membrane was washed a final time in TBST buffer for 5 minutes three times then exposed with 1 mL Super Signal West Femto (ThermoFisher, Ref #34095, Lot #UG286791) for 3 minutes before capturing a chemiluminescent image with an Azure 600 Western Blot Imager. A bright-field image of the membrane was then captured using the same imager and camera settings to obtain a size-accurate protein ladder that was aligned precisely to the western blot membrane.

#### Validating the purification of pig pTSC by western blot

20 µL aliquots of S3, S3 flow-through and each +F_ab_ and −F_ab_ elution fraction from the pig pTSC purification were mixed with 5 µL 5X Laemmeli buffer (200 mM Tris pH 6.8, 45% v/v glycerol, 5% w/v SDS, 0.0125% w/v Bromophenol Blue, 1.8M β-mercaptoethanol) then incubated at room temperature for 5 minutes before boiling them in a Mandel DryBath Plus heat block for 2 minutes at 95°C. A set of all these samples were then run on four 10% SDS-gels in running buffer (3.5 mM SDS, 25 mM Tris, 200 mM glycine, pH 8.3) for 65 minutes at 200V. The SDS-gels along with four Amersham Biosciences nitrocellulose membranes were then equilibrated in transfer buffer (50 mM Tris, 40 mM Glycine, 1 mM SDS, 10% v/v methanol, pH 8.3) for 5 minutes before being placed into transfer sandwiches submerged in transfer buffer. The proteins on the gel were transferred onto the nitrocellulose membranes for 120 minutes at 100V on ice. The membranes were then washed in TBST buffer (50 mM Tris, 100 mM NaCl, 1% v/v Tween 20) for 5 minutes three times before incubating in blocking buffer (50 mM Tris, 100 mM NaCl, 1% v/v Tween 20, 5% w/v skim milk powder) for 1 hour at room temperature. The membranes were washed again in TBST buffer for 5 minutes three times before incubating overnight for 17 hours in their respective primary antibody solutions: the first membrane was incubated in 10 mL rabbit anti-TSC1 primary antibody solution [50 mM Tris, 100 mM NaCl, 1% v/v Tween 20, 5% w/v skim milk powder, 1:1000 v/v rabbit anti-TSC1 antibody (Cell Signaling Technology, Clone #D43E2, Lot #3)], the second membrane was incubated in 10 mL mouse anti-TSC1 primary antibody solution [50 mM Tris, 100 mM NaCl, 1% v/v Tween 20, 5% w/v skim milk powder, 1:1000 v/v mouse anti-TSC2 antibody (Invitrogen, Ref #37-0400, Lot #UD288308)], the third membrane was incubated in 10 mL rabbit anti-TSC2 primary antibody solution [50 mM Tris, 100 mM NaCl, 1% v/v Tween 20, 5% w/v skim milk powder, 1:5000 v/v rabbit anti-TSC2 antibody (Cell Signaling Technology, Clone #D93F12, Lot #6)] and the fourth membrane was incubated in 10 mL rabbit anti-TBC1D7 primary antibody solution [50 mM Tris, 100 mM NaCl, 1% v/v Tween 20, 5% w/v skim milk powder, 1:1000 v/v rabbit anti-TBC1D7 antibody (Cell Signaling Technology, Clone #D8K1Y, Lot #3)]. The membranes were then washed in TBST buffer for 5 minutes three times then incubated for 1 hour at room temperature in their respective secondary antibody solutions: the membranes blotted with rabbit anti-TSC1 and rabbit anti-TSC2 primary antibodies were incubated in 10 mL goat anti-rabbit secondary antibody solution [50 mM Tris, 100 mM NaCl, 1% v/v Tween 20, 5% w/v skim milk powder, 1:5000 v/v goat anti-rabbit secondary antibody (Cell Signaling Technology, Product #7074P2, Lot #28)], the membrane blotted with mouse anti-TSC1 primary antibody was incubated in 10 mL goat anti-mouse secondary antibody solution [50 mM Tris, 100 mM NaCl, 1% v/v Tween 20, 5% w/v skim milk powder, 1:5000 v/v goat anti-mouse secondary antibody (Cell Signaling Technology, Product #7076S, Lot #33)] and the membrane blotted with anti-TBC1D7 primary antibody was incubated in 10 mL goat anti-rabbit F_c_ secondary antibody solution [50 mM Tris, 100 mM NaCl, 1% v/v Tween 20, 5% w/v skim milk powder, 1:5000 v/v goat anti-rabbit F_c_ secondary antibody (Abcam, Lot #GR3347004-5)]. The membranes were washed a final time in TBST buffer for 5 minutes three times then exposed with 1 mL Super Signal West Femto (ThermoFisher, Ref #34095, Lot #UG286791) for 3 minutes before capturing a chemiluminescent image with an Azure 600 Western Blot Imager. A bright-field image of the membranes were then captured using the same imager and camera settings to obtain size-accurate protein ladders that were aligned precisely to their respective western blot membranes.

#### Validating the purification of rat pTSC by western blot

20 uL aliquots of S3, S3 flow-through and each +F_ab_ and −F_ab_ elution fraction from the rat pTSC purification were mixed with 5 µL 5X Laemmeli buffer (200 mM Tris pH 6.8, 45% v/v glycerol, 5% w/v SDS, 0.0125% w/v Bromophenol Blue, 1.8M β-mercaptoethanol) then incubated at room temperature for 5 minutes before boiling them in a Mandel DryBath Plus heat block for 2 minutes at 95°C. A set of all these samples were then run on three 10% SDS-gels in running buffer (3.5 mM SDS, 25 mM Tris, 200 mM glycine, pH 8.3) for 65 minutes at 200V. The SDS-gels along with three Amersham Biosciences nitrocellulose membranes were then equilibrated in transfer buffer (50 mM Tris, 40 mM Glycine, 1 mM SDS, 10% v/v methanol, pH 8.3) for 5 minutes before being placed into transfer sandwiches submerged in transfer buffer. The proteins on the gel were transferred onto the nitrocellulose membranes for 120 minutes at 100V on ice. The membranes were then washed in TBST buffer (50 mM Tris, 100 mM NaCl, 1% v/v Tween 20) for 5 minutes three times before incubating in blocking buffer (50 mM Tris, 100 mM NaCl, 1% v/v Tween 20, 5% w/v skim milk powder) for 1 hour at room temperature. The membranes were washed again in TBST buffer for 5 minutes three times before incubating overnight for 17 hours in their respective primary antibody solutions: the first membrane was incubated in 10 mL rabbit anti-TSC1 primary antibody solution [50 mM Tris, 100 mM NaCl, 1% v/v Tween 20, 5% w/v skim milk powder, 1:1000 v/v rabbit anti-TSC1 antibody (Cell Signaling Technology, Clone #D43E2, Lot #3)], the second membrane was incubated in 10 mL rabbit anti-TSC2 primary antibody solution [50 mM Tris, 100 mM NaCl, 1% v/v Tween 20, 5% w/v skim milk powder, 1:5000 v/v rabbit anti-TSC2 antibody (Cell Signaling Technology, Clone #D93F12, Lot #6)] and the last membrane was incubated in 10 mL rabbit anti-TBC1D7 primary antibody solution [50 mM Tris, 100 mM NaCl, 1% v/v Tween 20, 5% w/v skim milk powder, 1:1000 v/v rabbit anti-TBC1D7 antibody (Cell Signaling Technology, Clone #D8K1Y, Lot #3)]. The membranes were then washed in TBST buffer for 5 minutes three times then incubated for 1 hour at room temperature in their respective secondary antibody solutions: the membranes blotted with rabbit anti-TSC1 and rabbit anti-TSC2 primary antibodies were incubated in 10 mL goat anti-rabbit secondary antibody solution [50 mM Tris, 100 mM NaCl, 1% v/v Tween 20, 5% w/v skim milk powder, 1:5000 v/v goat anti-rabbit secondary antibody (Cell Signaling Technology, Product #7076S, Lot #33)] and the membrane blotted with rabbit anti-TBC1D7 primary antibody was incubated in 10 mL goat anti-rabbit F_c_ secondary antibody solution [50 mM Tris, 100 mM NaCl, 1% v/v Tween 20, 5% w/v skim milk powder, 1:5000 v/v goat anti-rabbit F_c_ secondary antibody (Abcam, Lot #GR3347004-5)]. The membranes were washed a final time in TBST buffer for 5 minutes three times then exposed with 1 mL Super Signal West Femto (ThermoFisher, Ref #34095, Lot #UG286791) for 3 minutes before capturing a chemiluminescent imagewith an Azure 600 Western Blot Imager. A bright-field image of the membranes were then captured using the same imager and camera settings to obtain size-accurate protein ladders that were aligned precisely to their respective western blot membranes.

#### Negative stain electron microscopy of purified pig pTSC

400-mesh copper-rhodium grids (Electron Microscopy Sciences, Lot #191202) coated with collodion and 10 nm carbon (in a Leica EM ACE200 Vacuum Coater were glow discharged in air with a Pelco EasiGlow glow discharger at settings of: 10 second process, 10 second hold and 15 mA current. 5 µL +F_ab_ and −F_ab_ elution fractions 2 from the pig and rat pTSC purifications were then pipetted onto the surface of the carbon side of the grid and incubated there for 3 minutes at room temperature or overnight in a humid Petri dish kept at 4 °C. The grids were then blotted with filter paper (Whatman grade 1, Cat #1001-090, Lot #17031753) before washing them for 5 seconds in three 50 µL droplets of MilliQ water whilst blotting on filter paper in between. The grids were then immediately stained for 10 seconds in two 50 µL droplets of 2% w/v uranyl acetate, blotting with filter paper between droplets. Imaging was done with a Talos L120C electron microscope (Thermo Fisher Scientific) operating at 120kV using a lanthanum hexaboride (LaB_6_) emitter and images were recorded with a Ceta camera. Defocus was set to −1 to −2 μm range. Nominal magnification was at 57,000x with a total electron exposure of 50 e/Å^2^. Calibrated pixel size was at 2.45Å. Micrographs were converted to PNG format using EMAN2 software[20].

## Data and Materials Availability

Data and reagents used in this study are available upon request.

## Acknowledgements

This study was supported by Princess Margaret Cancer Foundation (Toronto, Canada), Canadian Institute of Health Research (grant# MOP-419240), and Natural sciences and engineering research council of Canada (grant# RGPIN-2018-06070). Talos L120C was funded by Canada Foundation for Innovation (CFI) and Ontario Research Fund-Research Innovation (ORF-RI). We thank Vuk Stambolic for providing the pTSC expression vectors and critical review of our manuscript.

## Author contributions

**DLD** and **MTMJ**: conceptualize the study. **DLD**: expressed and purified F_ab_, prepared the brain homogenates, purified the pTSC, performed the western blot analysis, and collected the EM images. **SMNH**: performed the expression and purification optimizations on the F_ab_. **DLD** and **GW**: processed EM images. **SAB**: cloned the 3×FLAG TSC2 expression vector. **MTMJ**: expressed, purified, and performed negative stain EM analysis of the recombinant pTSC. **DLD, JPJ, JLR, MTMJ**: wrote the manuscripts and prepared the figures.

