## Supplementary material for "Polymeric assembly of endogenous Tuberous Sclerosis Protein Complex": README.txt

Supplemantary_data_7 - file identifiers

12E_1 - 12E_21: elution fraction 2, +Fab TSC2, overnight incubation on the grid, 0.5% DDM
13B_1 - 13B_2: elution fraction 2, -Fab TSC2, overnight incubation on the grid, 0.5% DDM
