## Supplementary material for "Polymeric assembly of endogenous Tuberous Sclerosis Protein Complex": README.txt

Supplemantary_data_8 - file identifiers. 

16A_0000 - 16A_0016: elution fraction 2, +Fab pTSC, 3 minutes incubation on the grid, 0.1% DDM
16C_0000 - 16C_0009: elution fraction 2, +Fab pTSC, 3 minutes incubation on the grid, 0.5% DDM
16E_0000 - 16E_0007: elution fraction 2, -Fab pTSC, 3 minutes incubation on the grid, 0.1% DDM
