## Supplementary figures and images for "Polymeric assembly of endogenous Tuberous Sclerosis Protein Complex"

### 12E_1_1.pdf

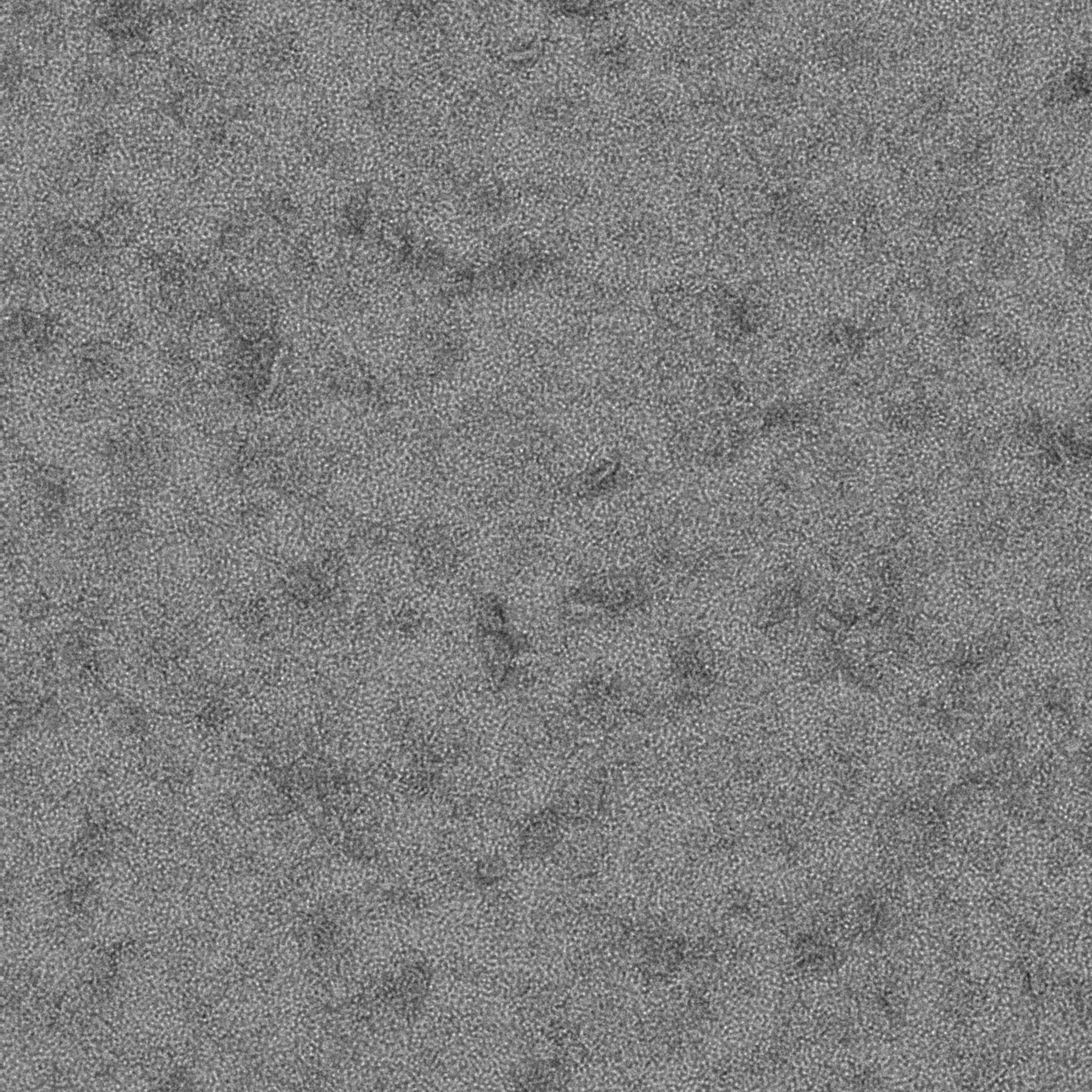

### 12E_2_1.pdf

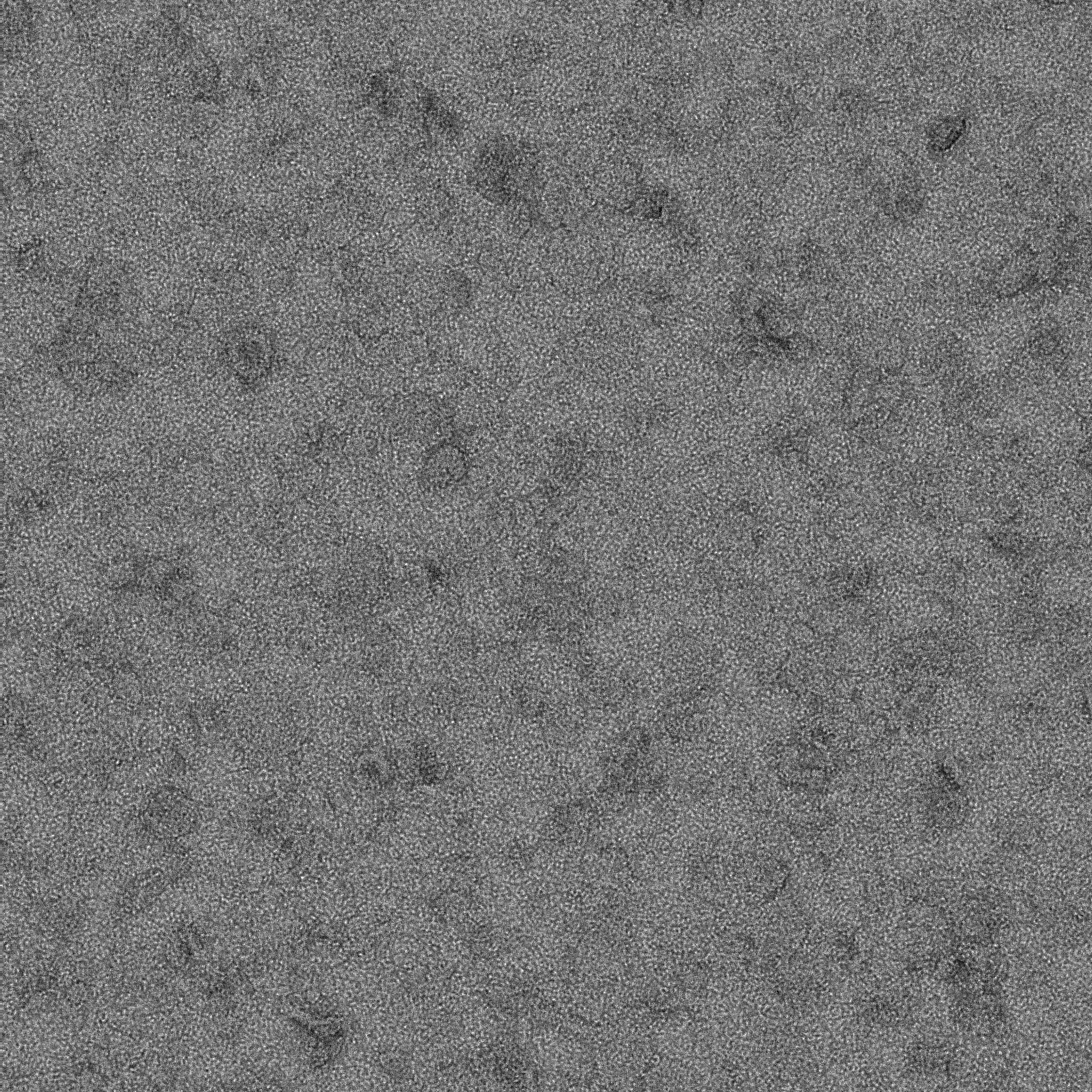

### 12E_3_1.pdf

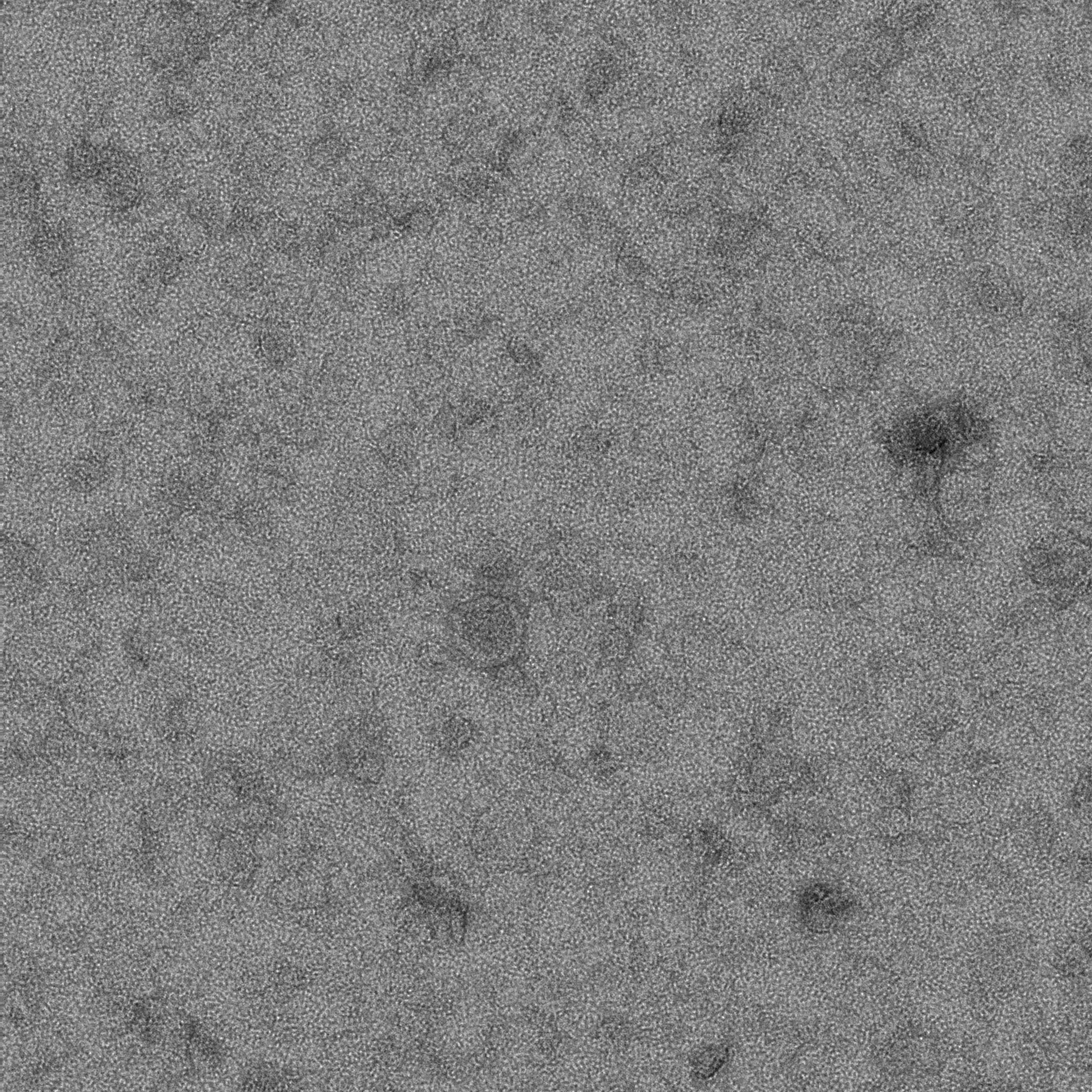

### 12E_4_1.pdf

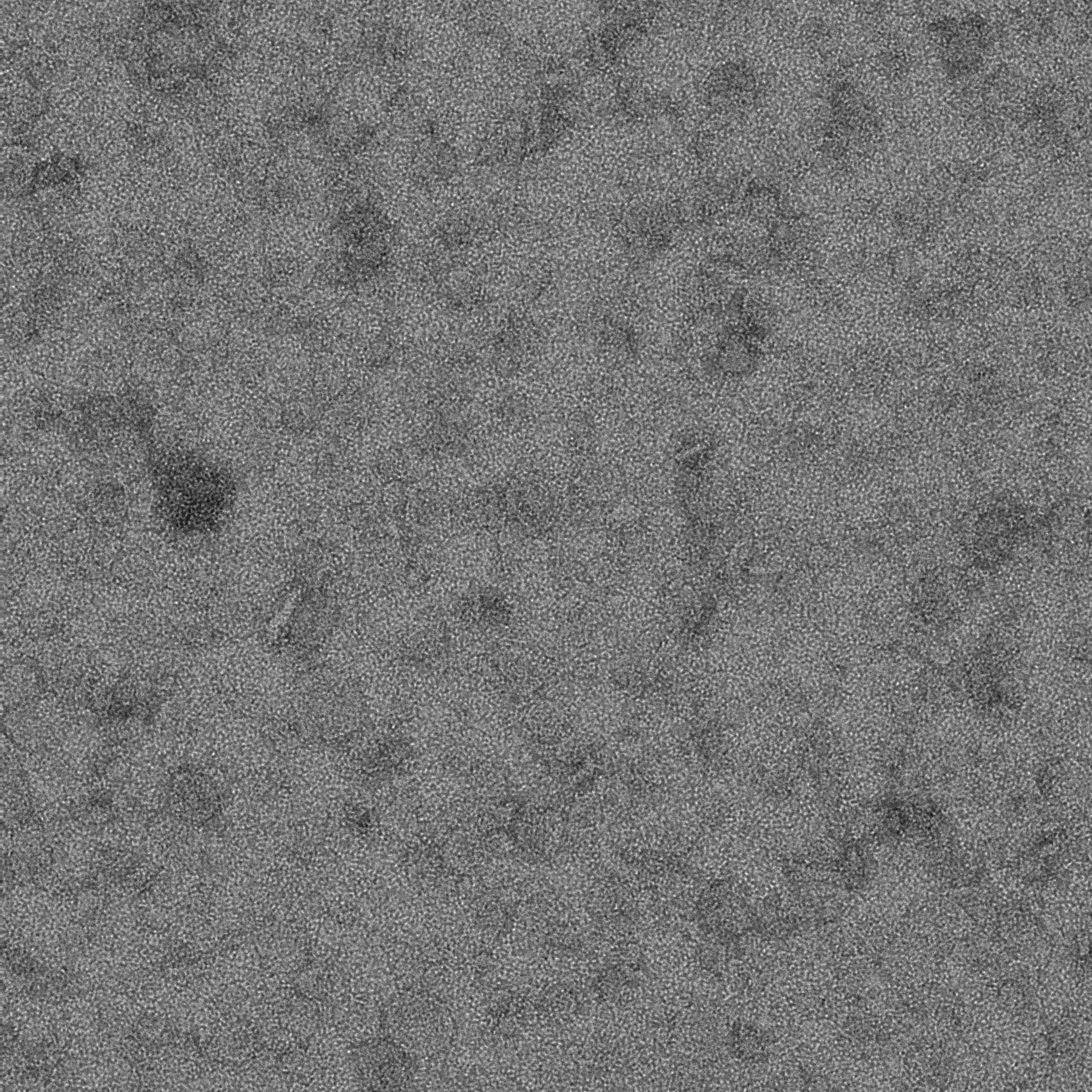

### 12E_5_1.pdf

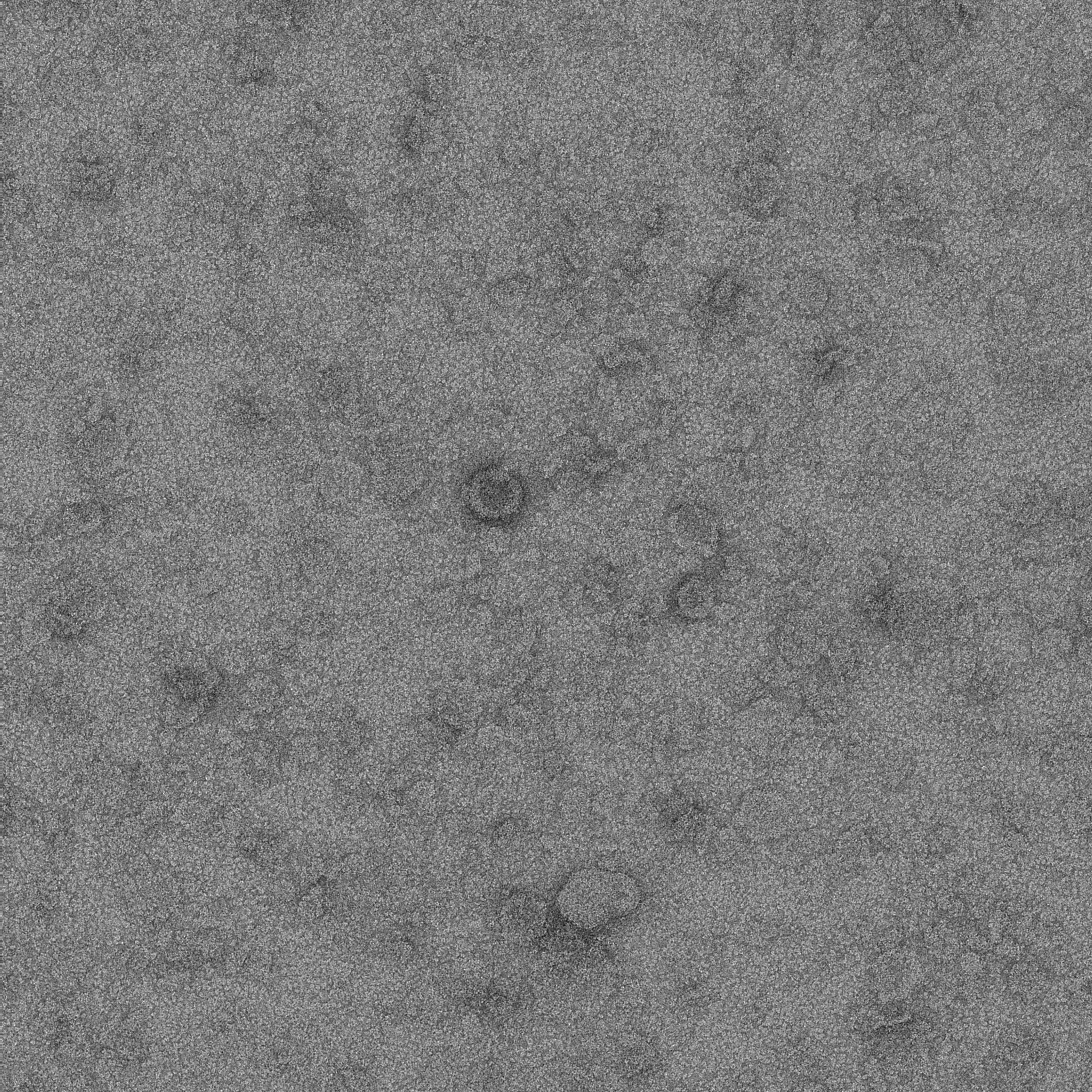

### 12E_6_1.pdf

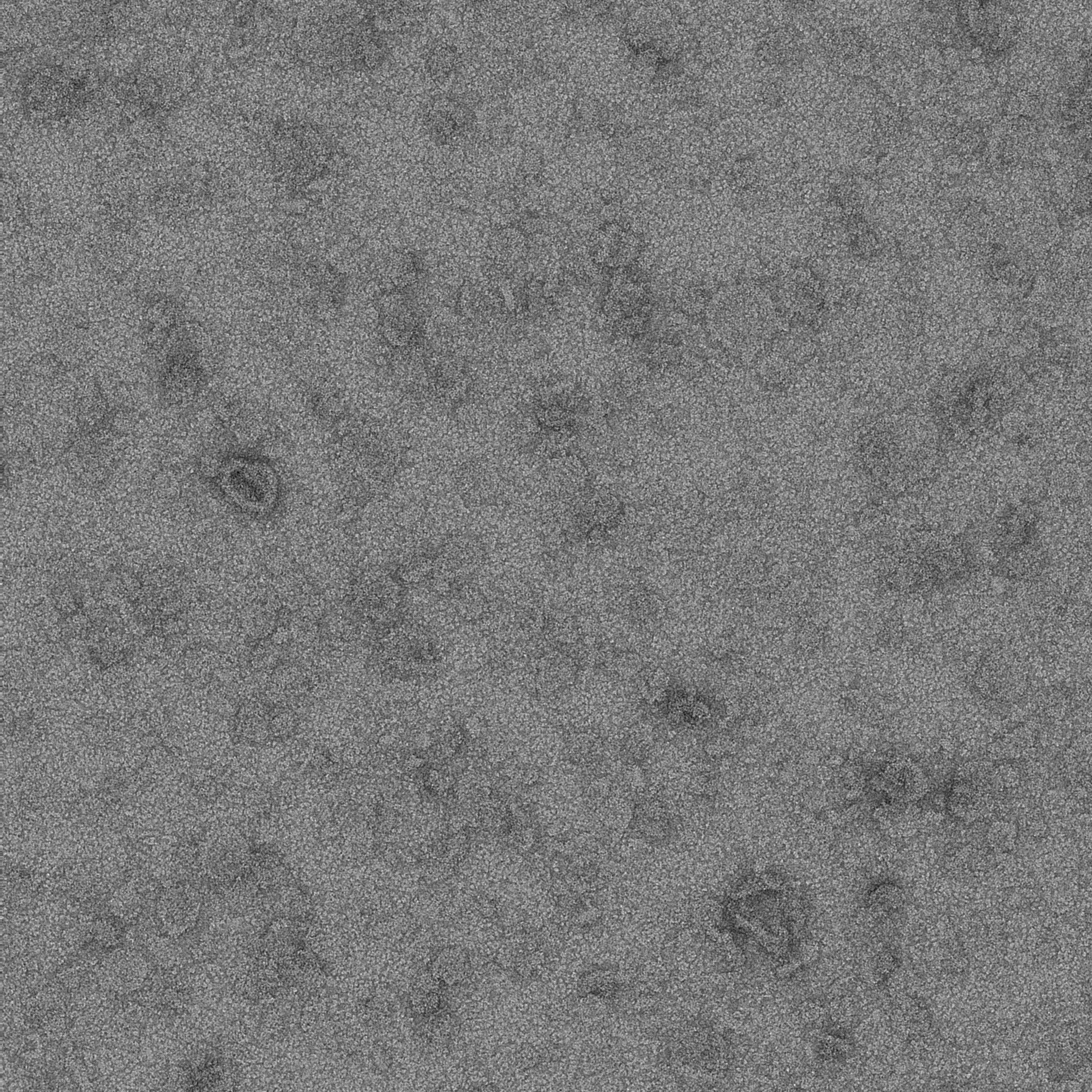

### 12E_7_1.pdf

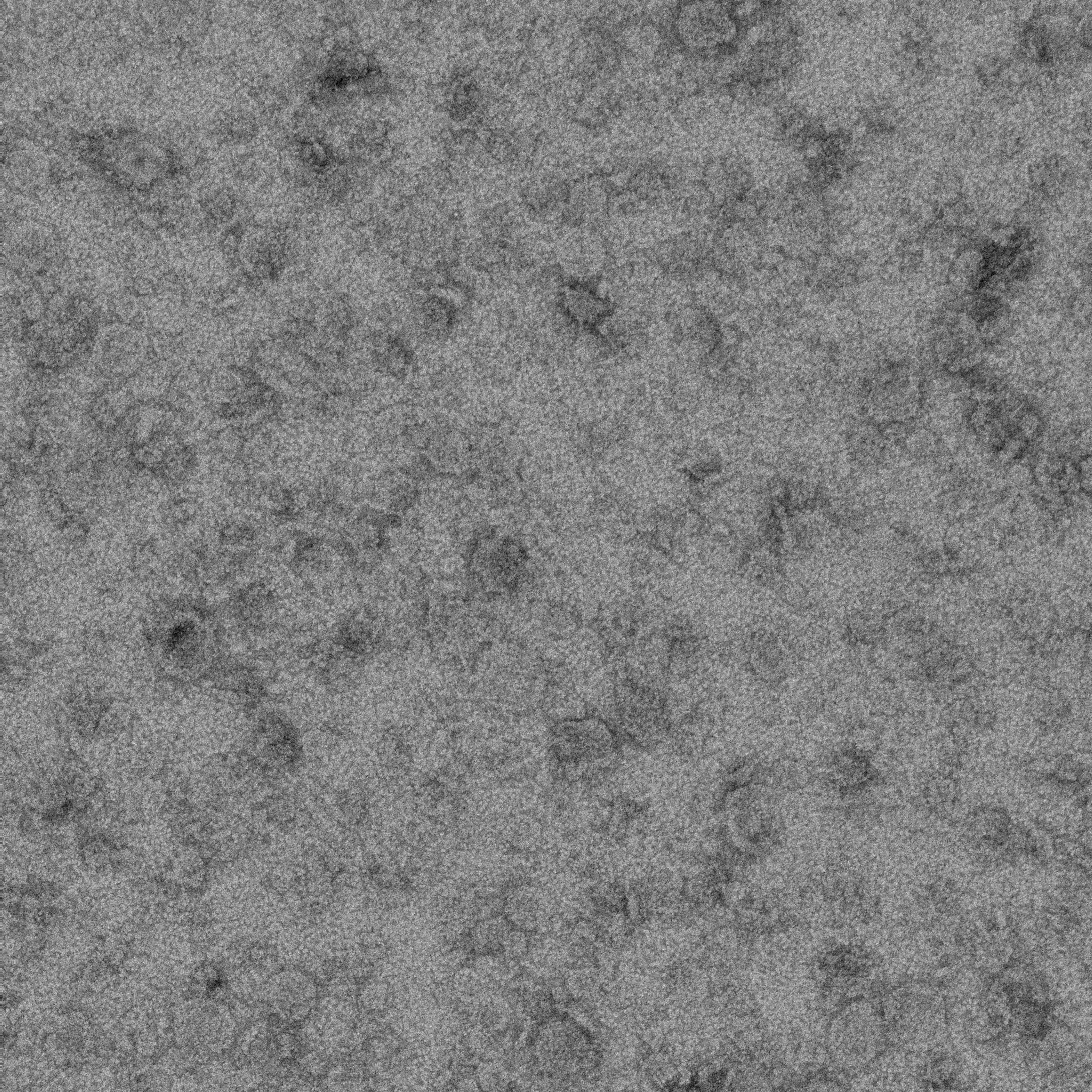

### 12E_8_1.pdf

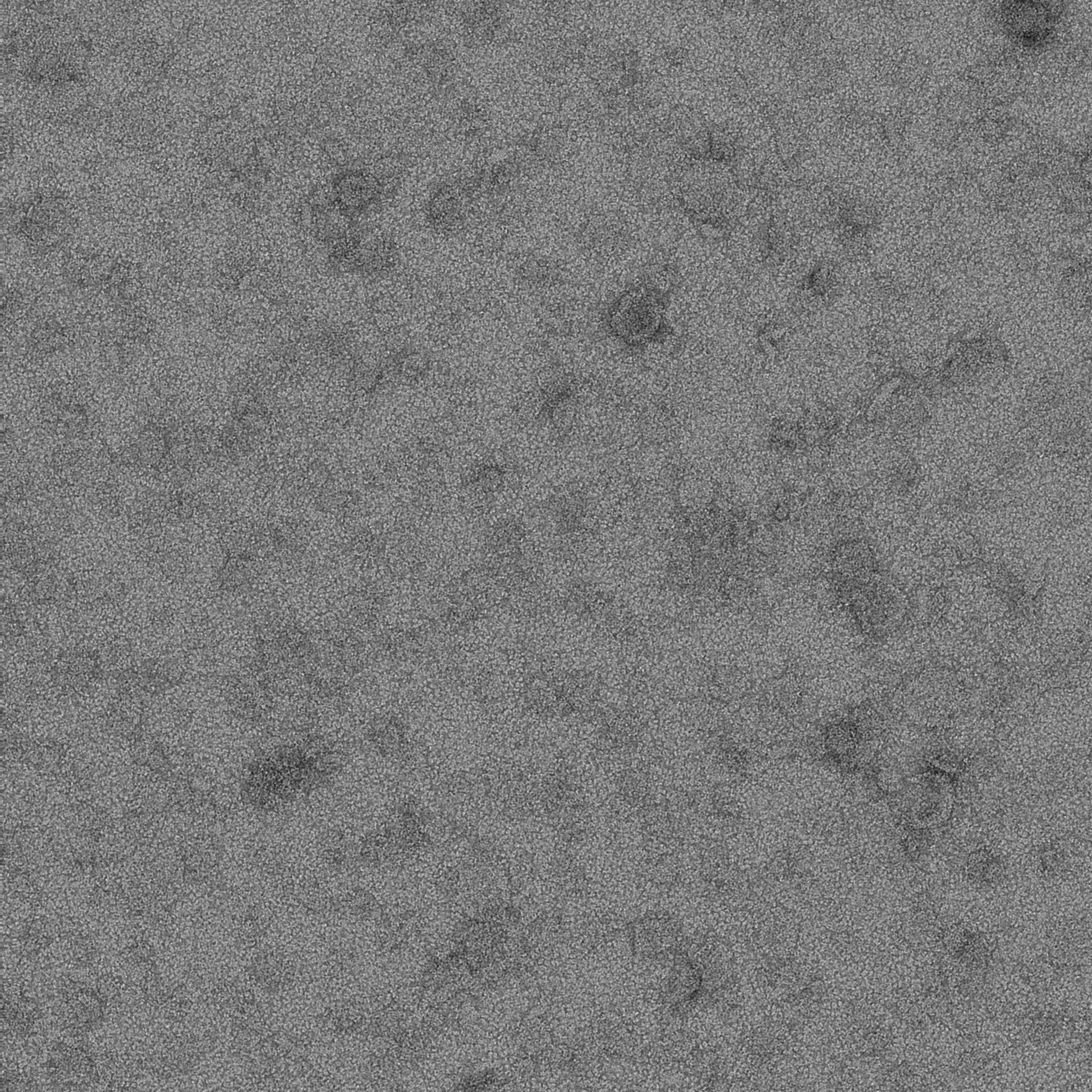

### 12E_9_1.pdf

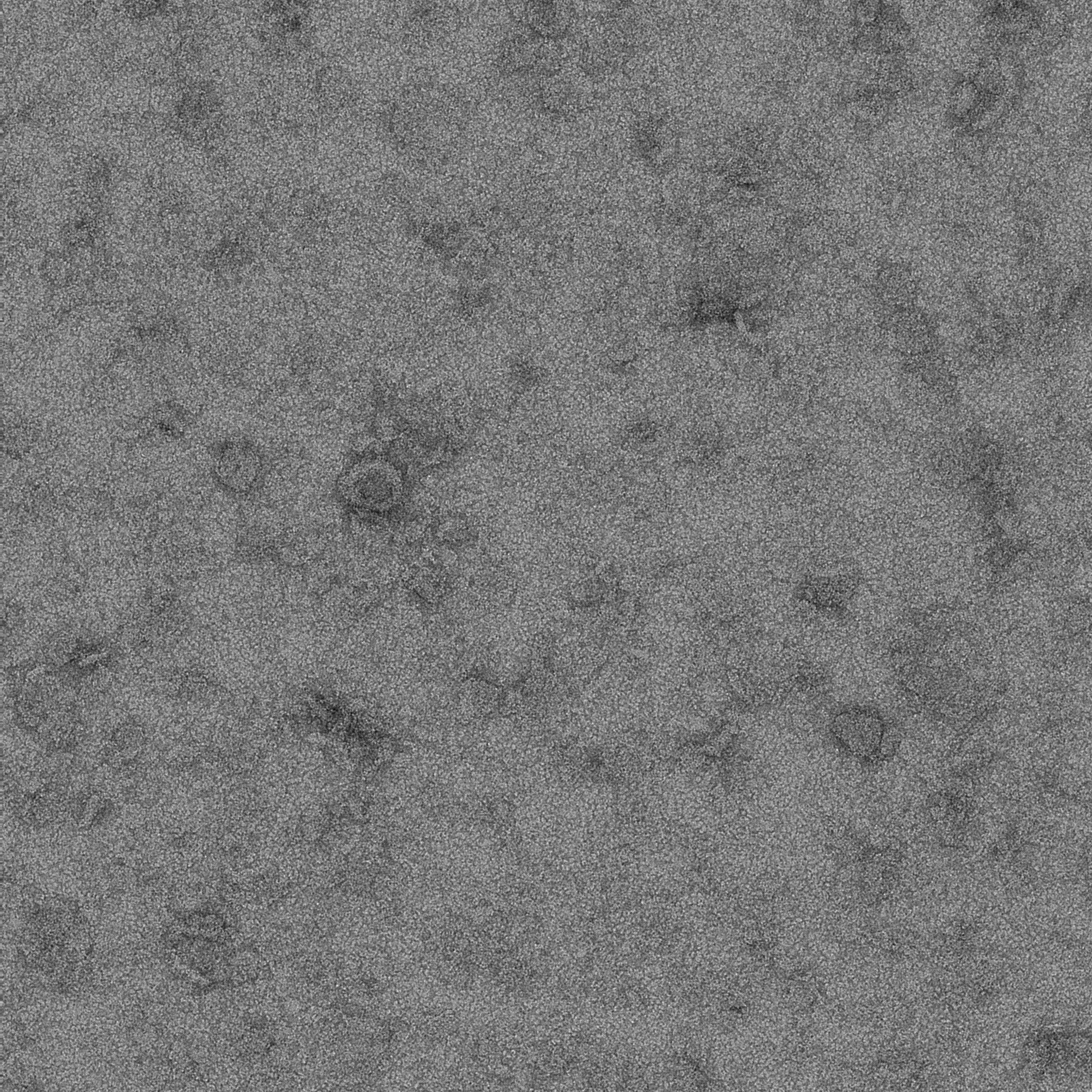

### 12E_10_1.pdf

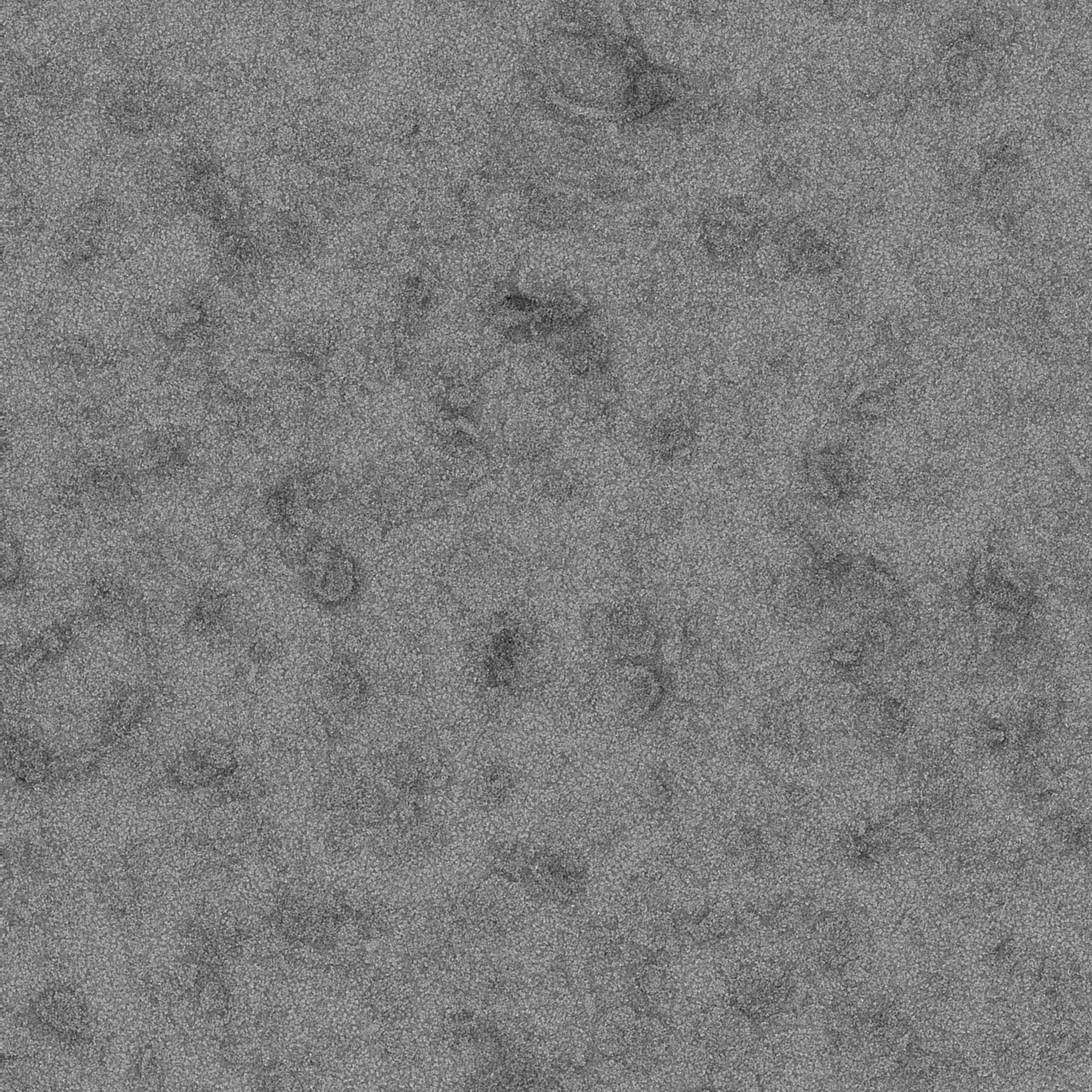

### 12E_11_1.pdf

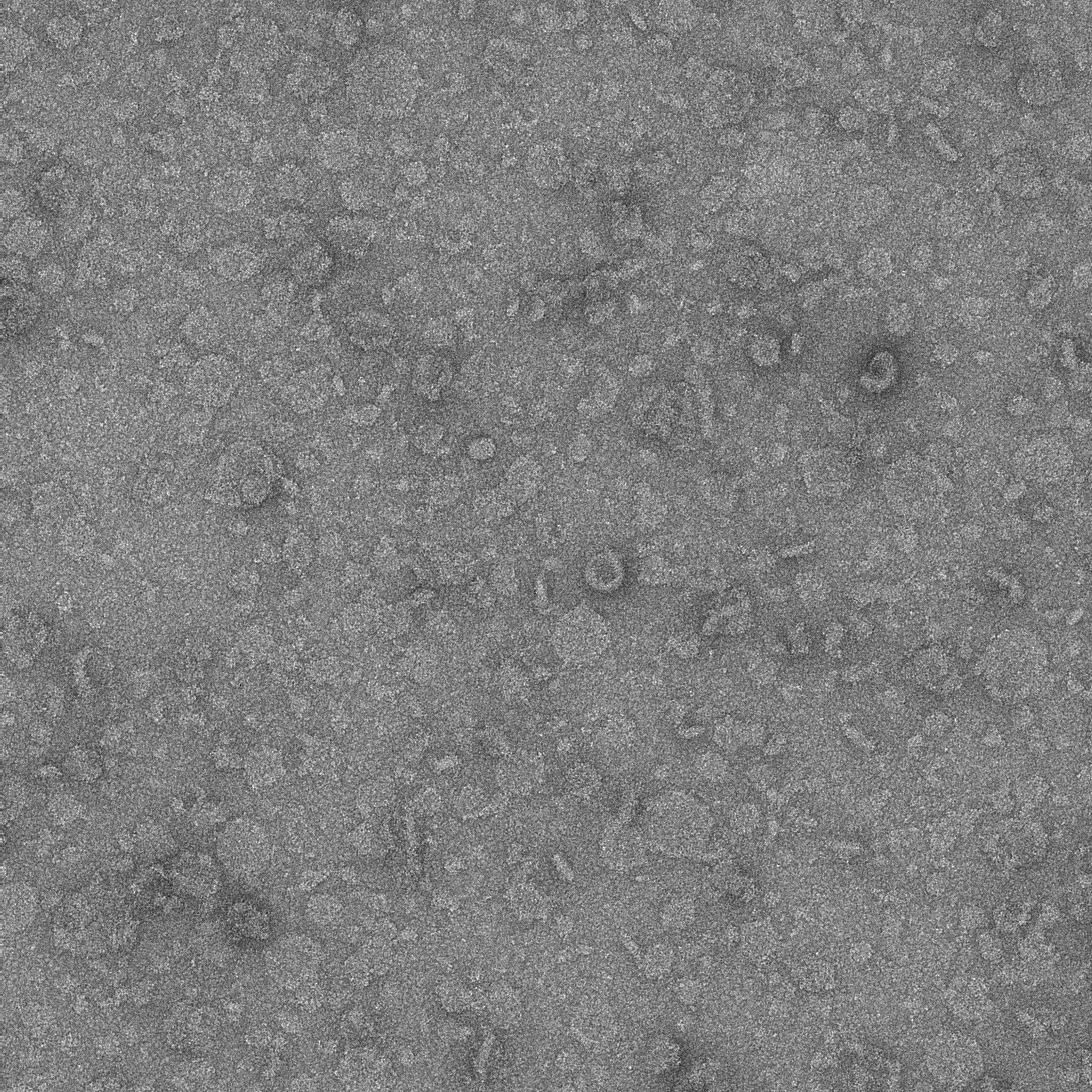

### 12E_12_1.pdf

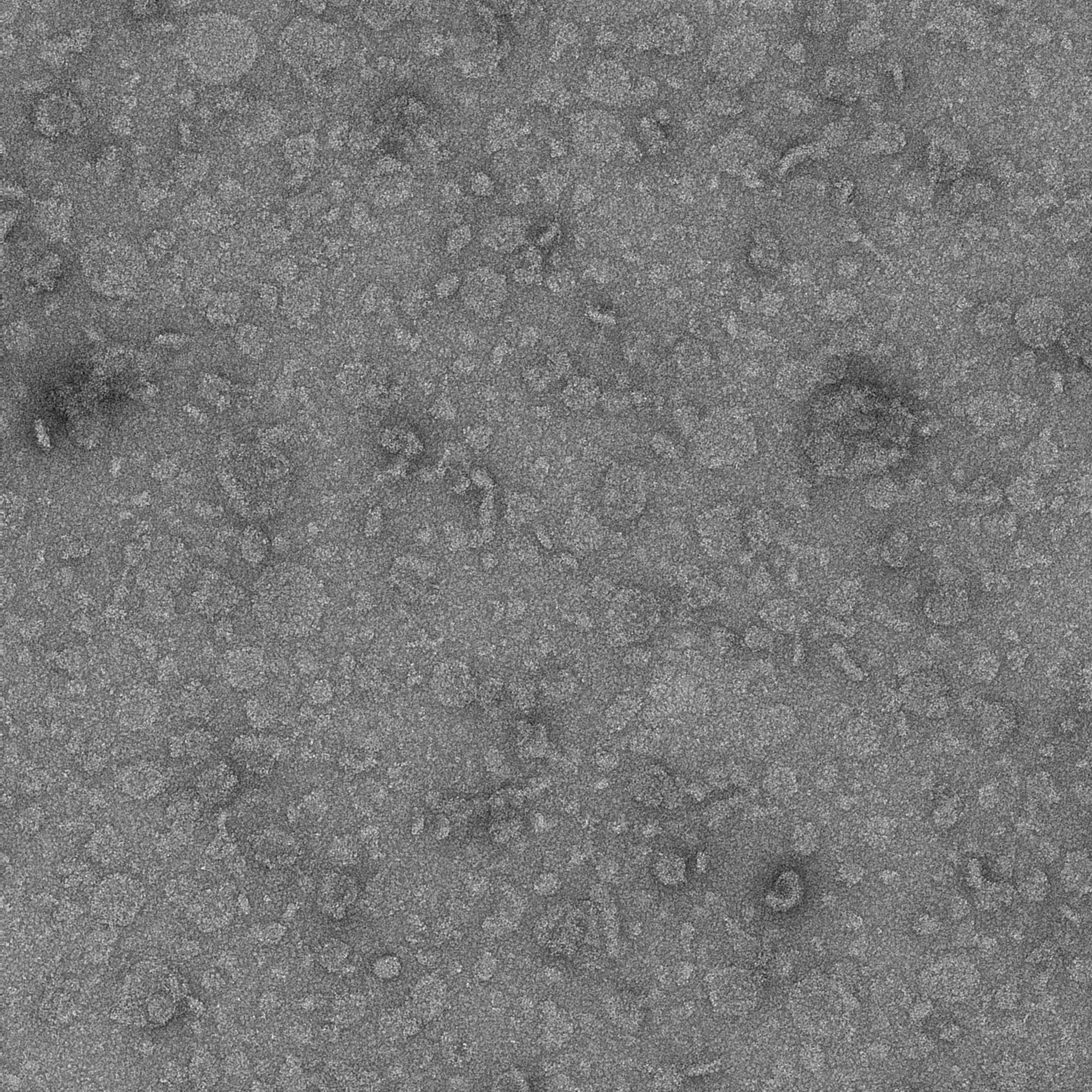

### 12E_13_1.pdf

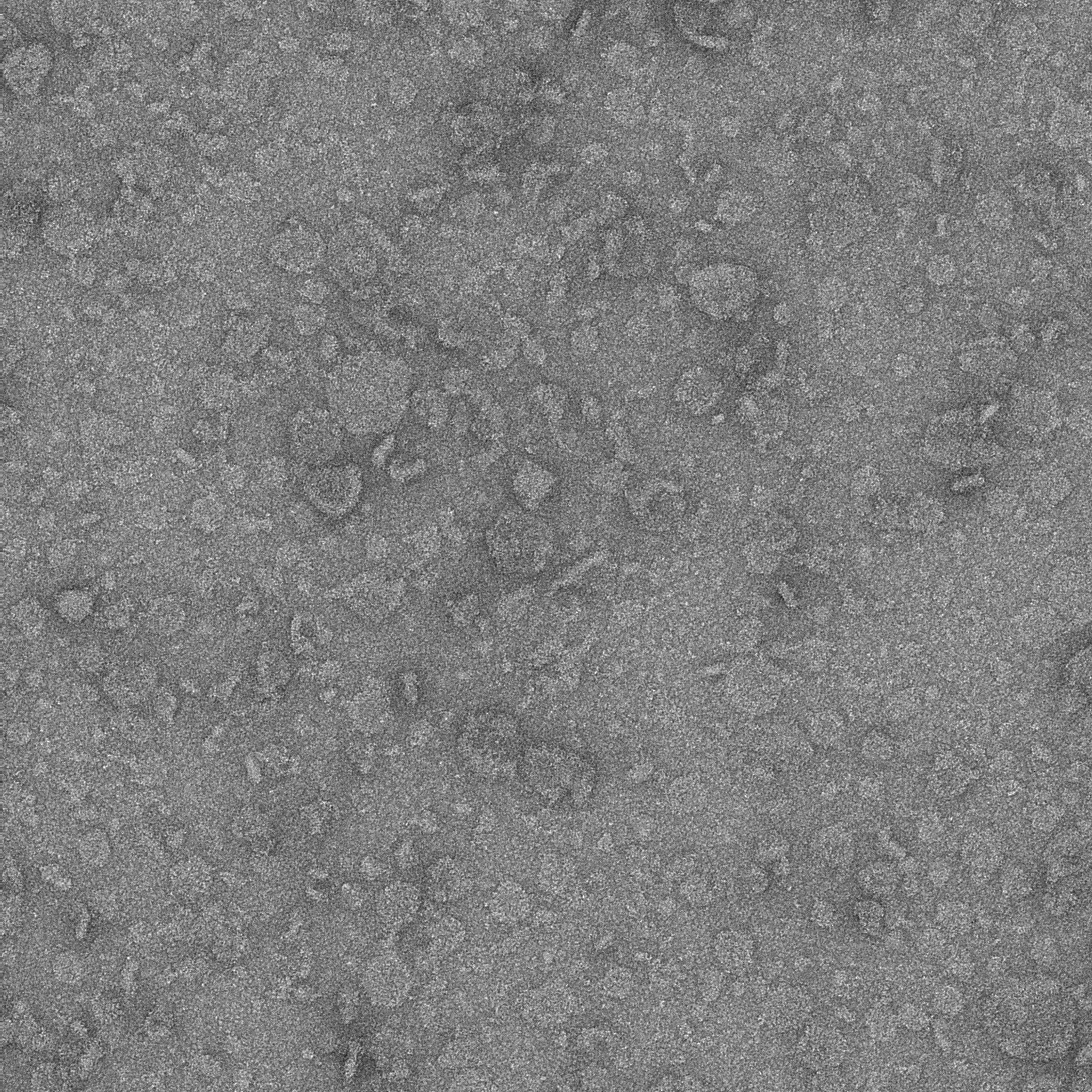

### 12E_14_1.pdf

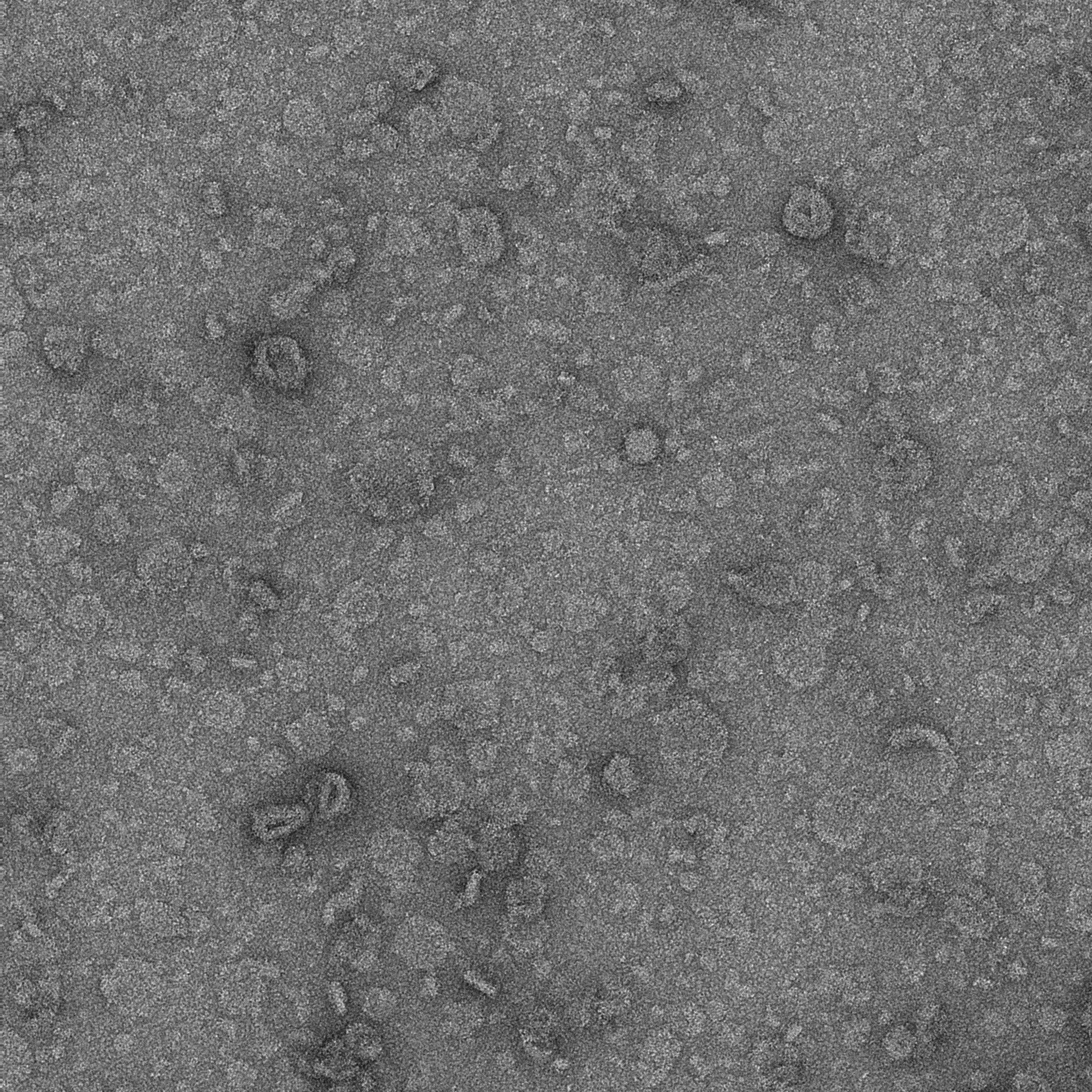

### 12E_15_1.pdf

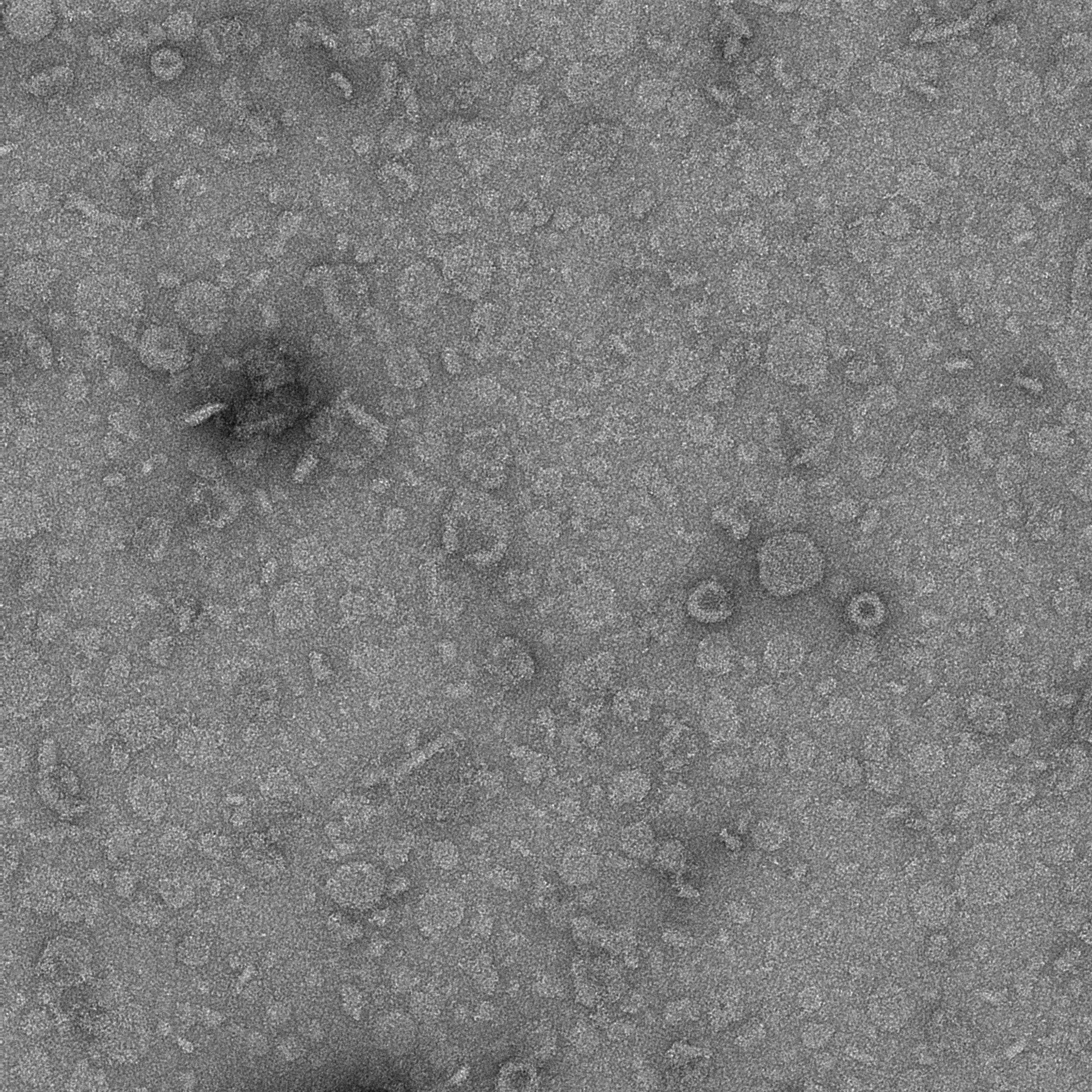

### 12E_16_1.pdf

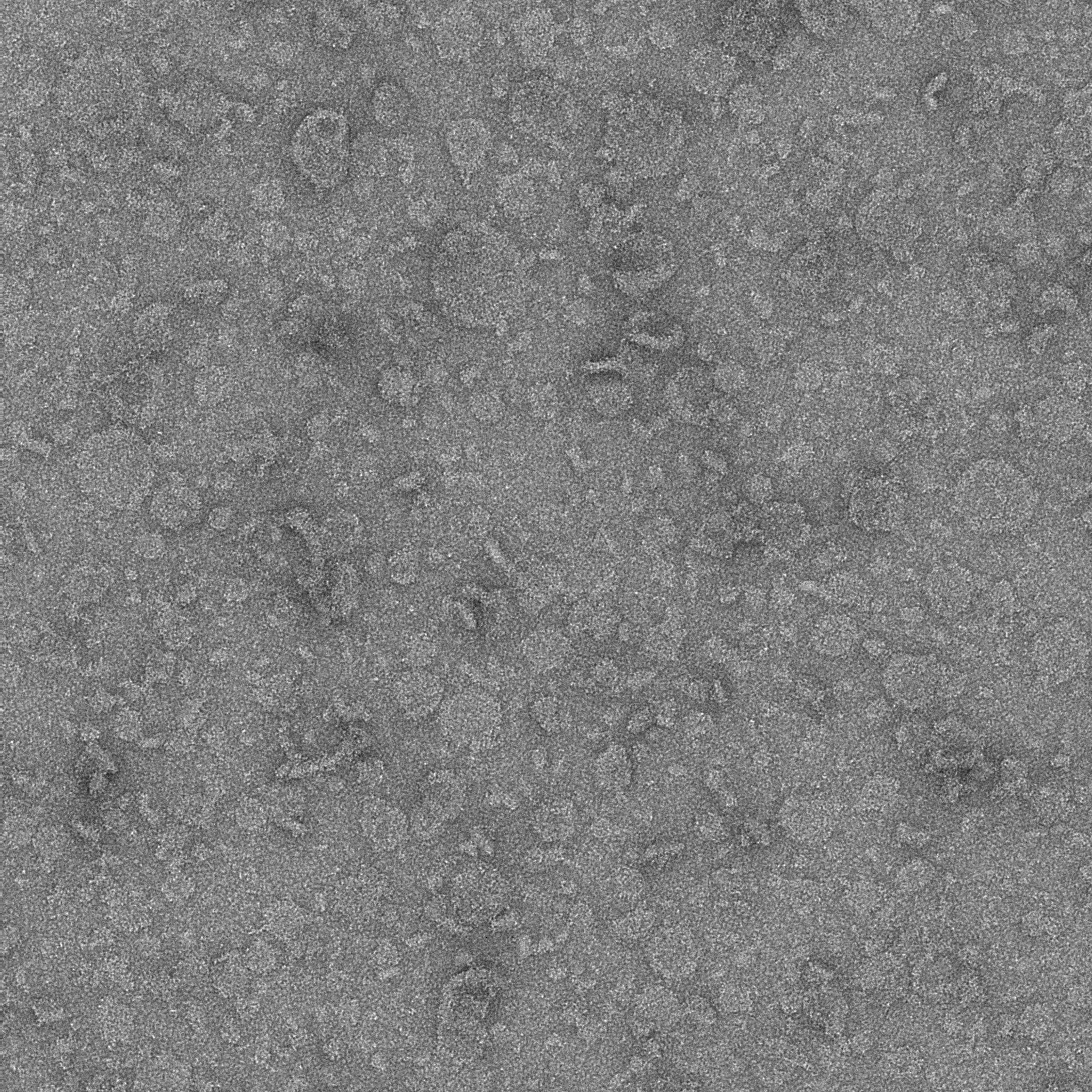

### 12E_17_1.pdf

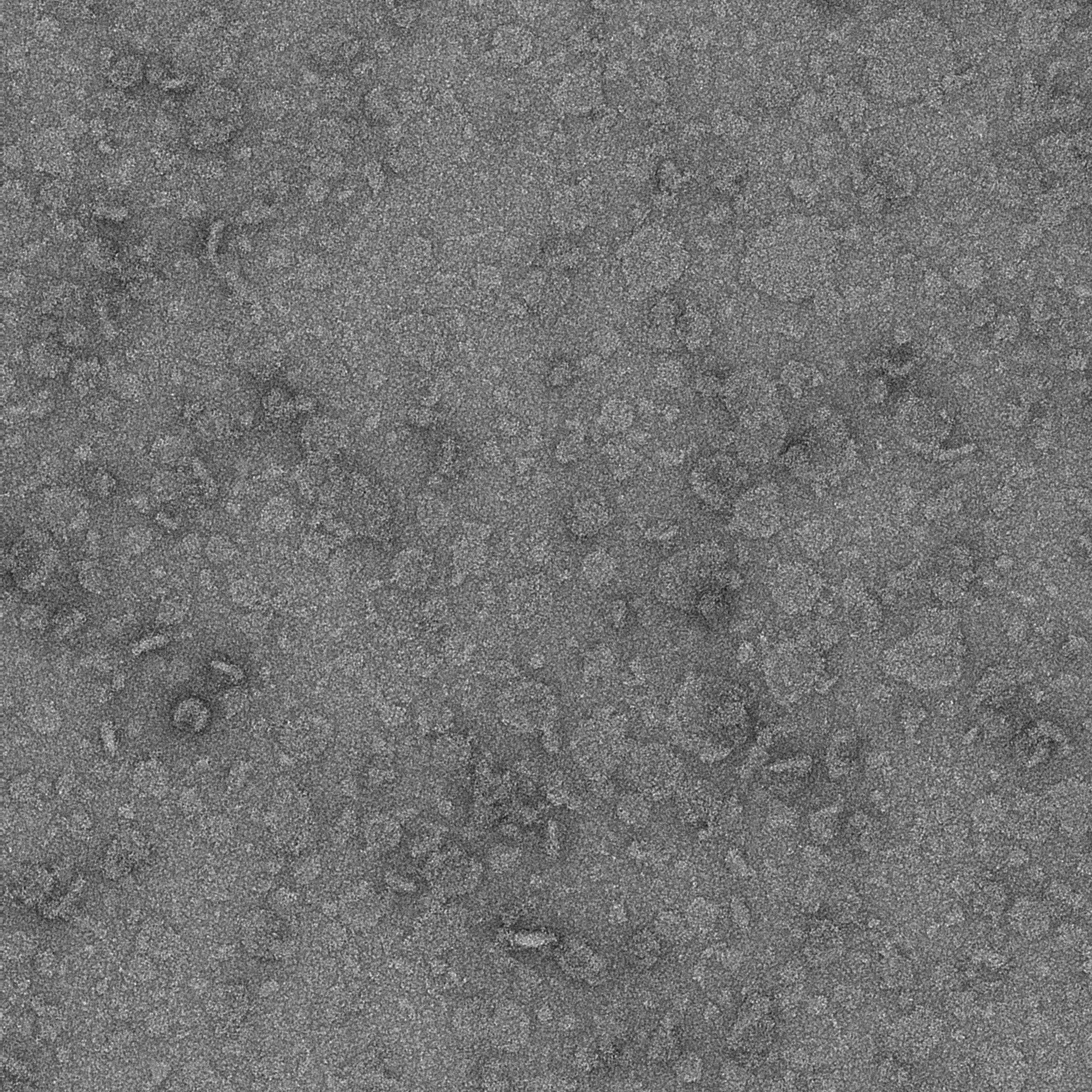

### 12E_18_1.pdf

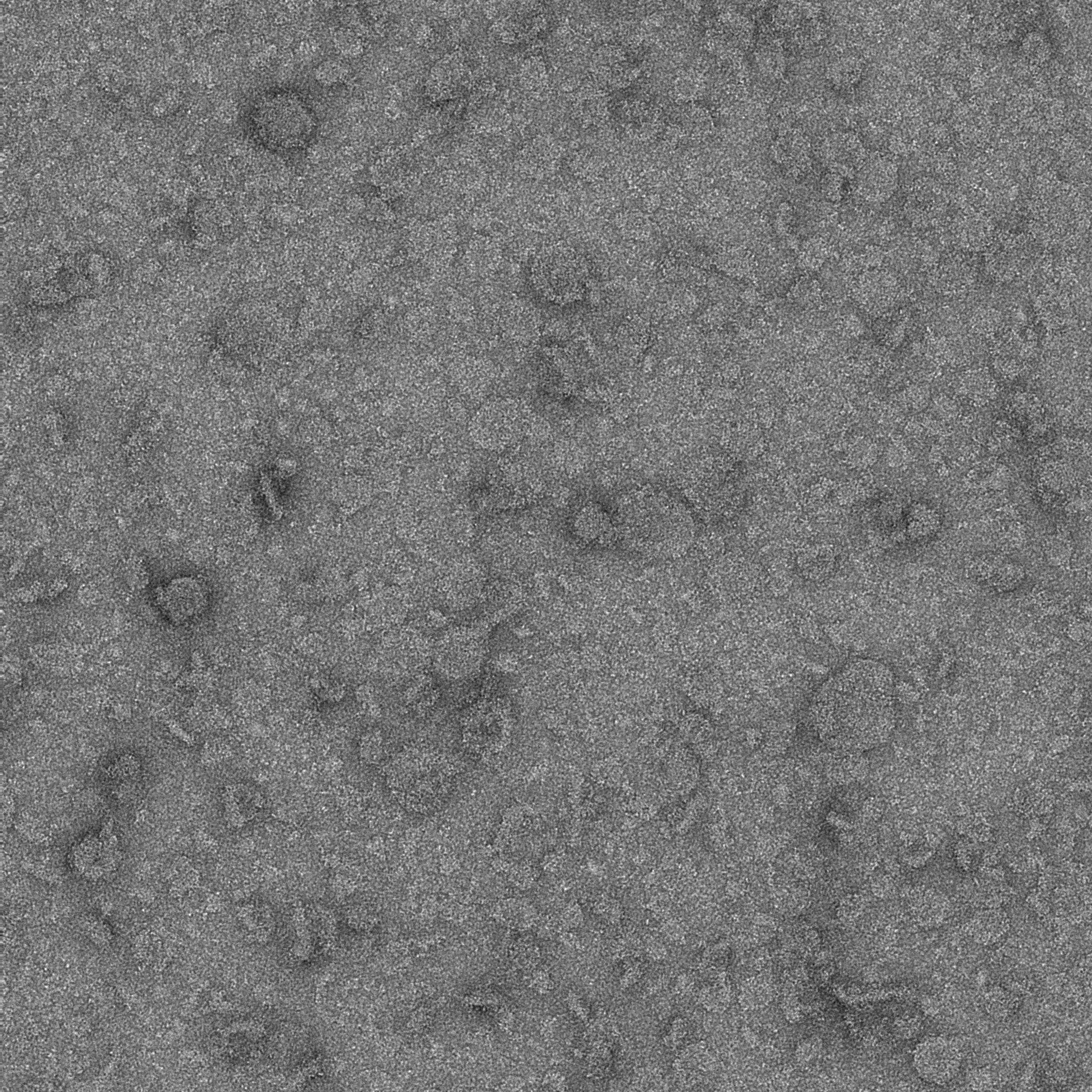

### 12E_19_1.pdf

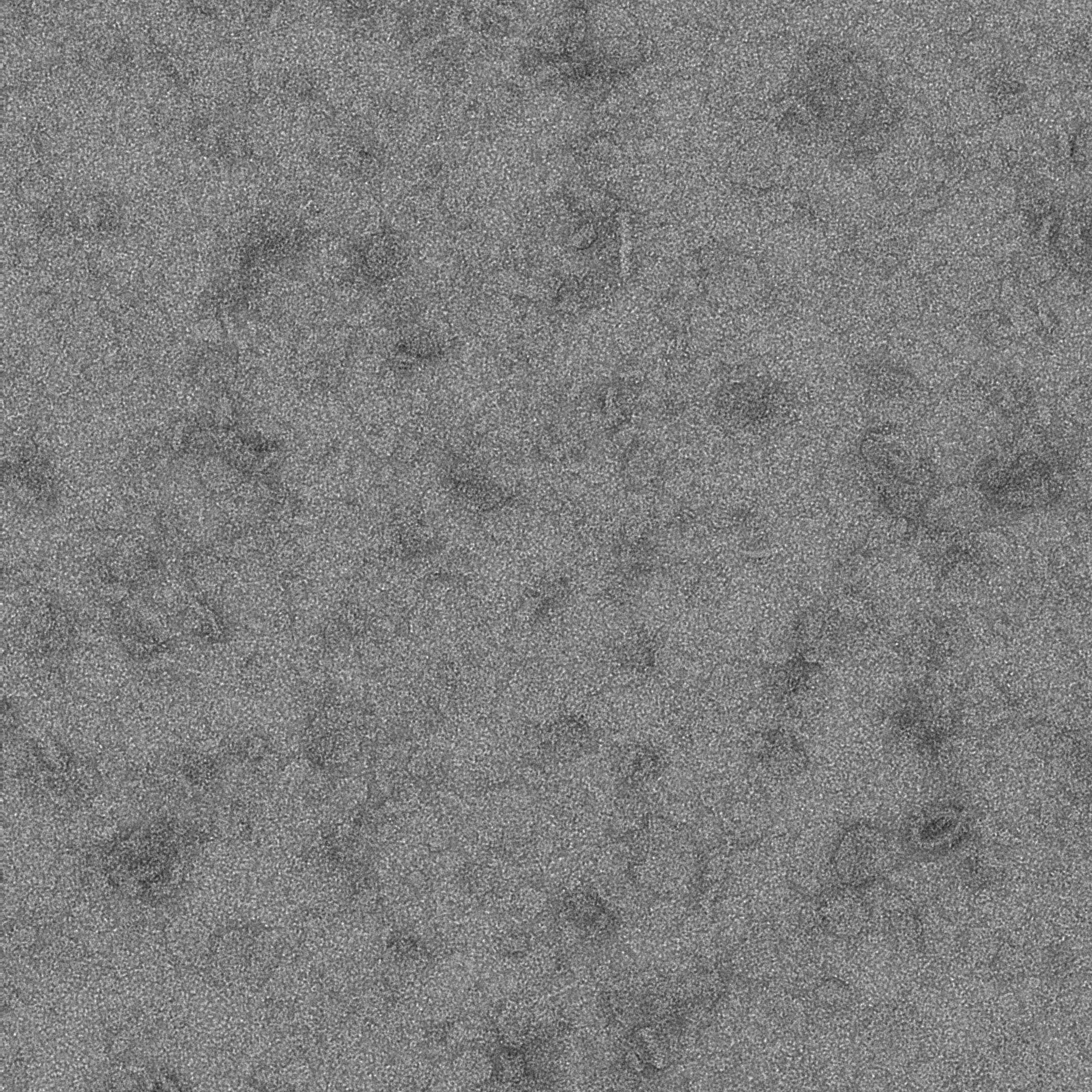

### 12E_20_1.pdf

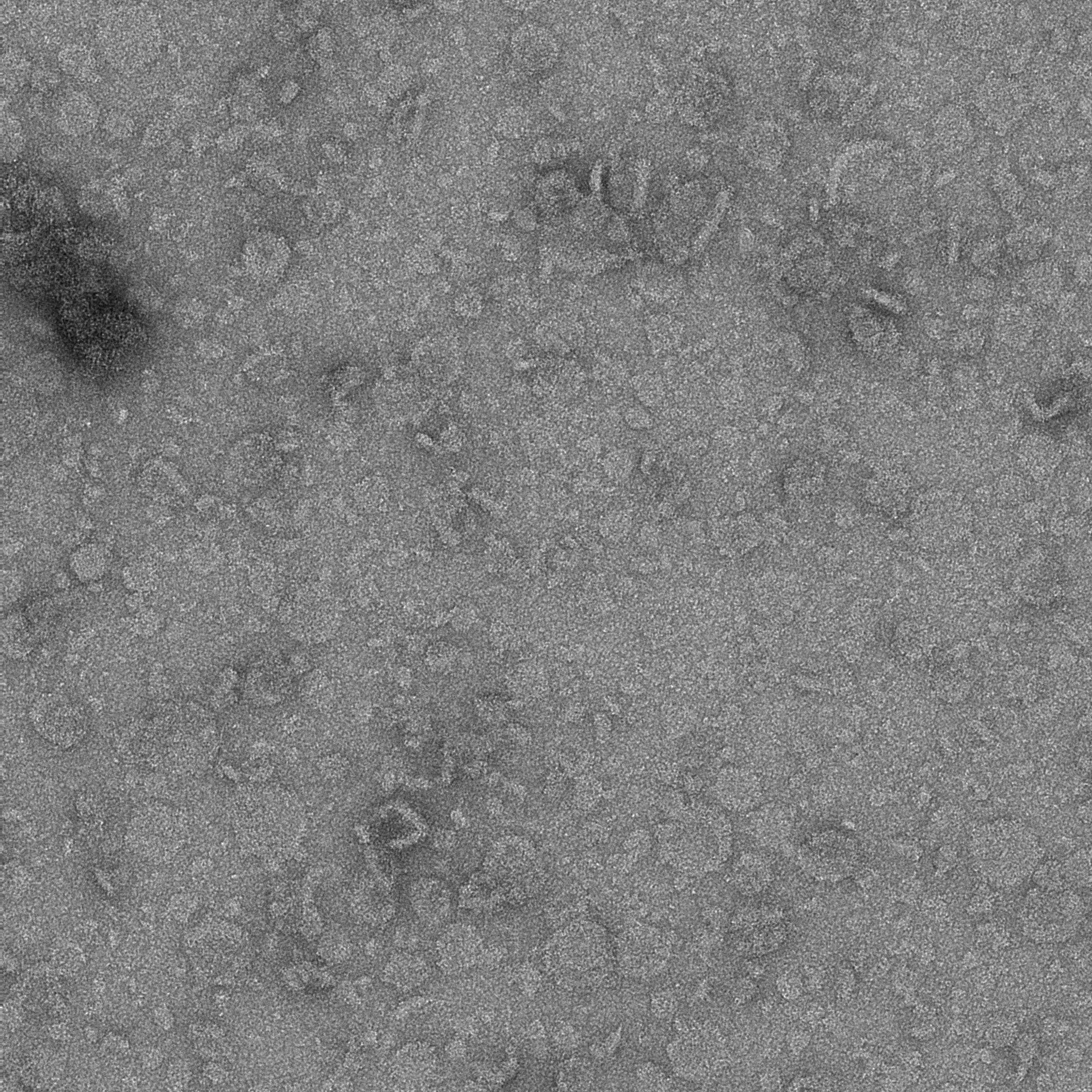

### 12E_21_1.pdf

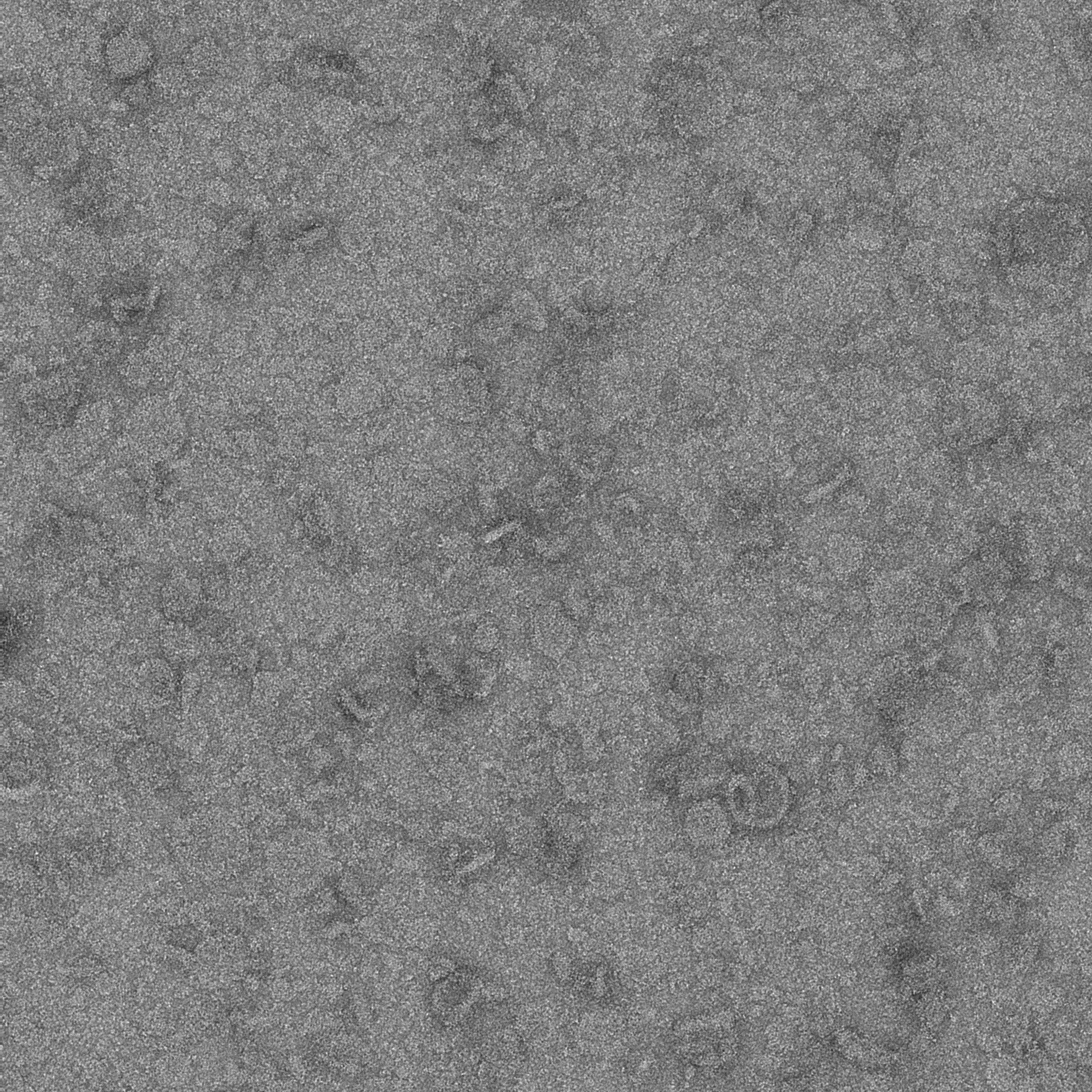

### 13B_1_1.pdf

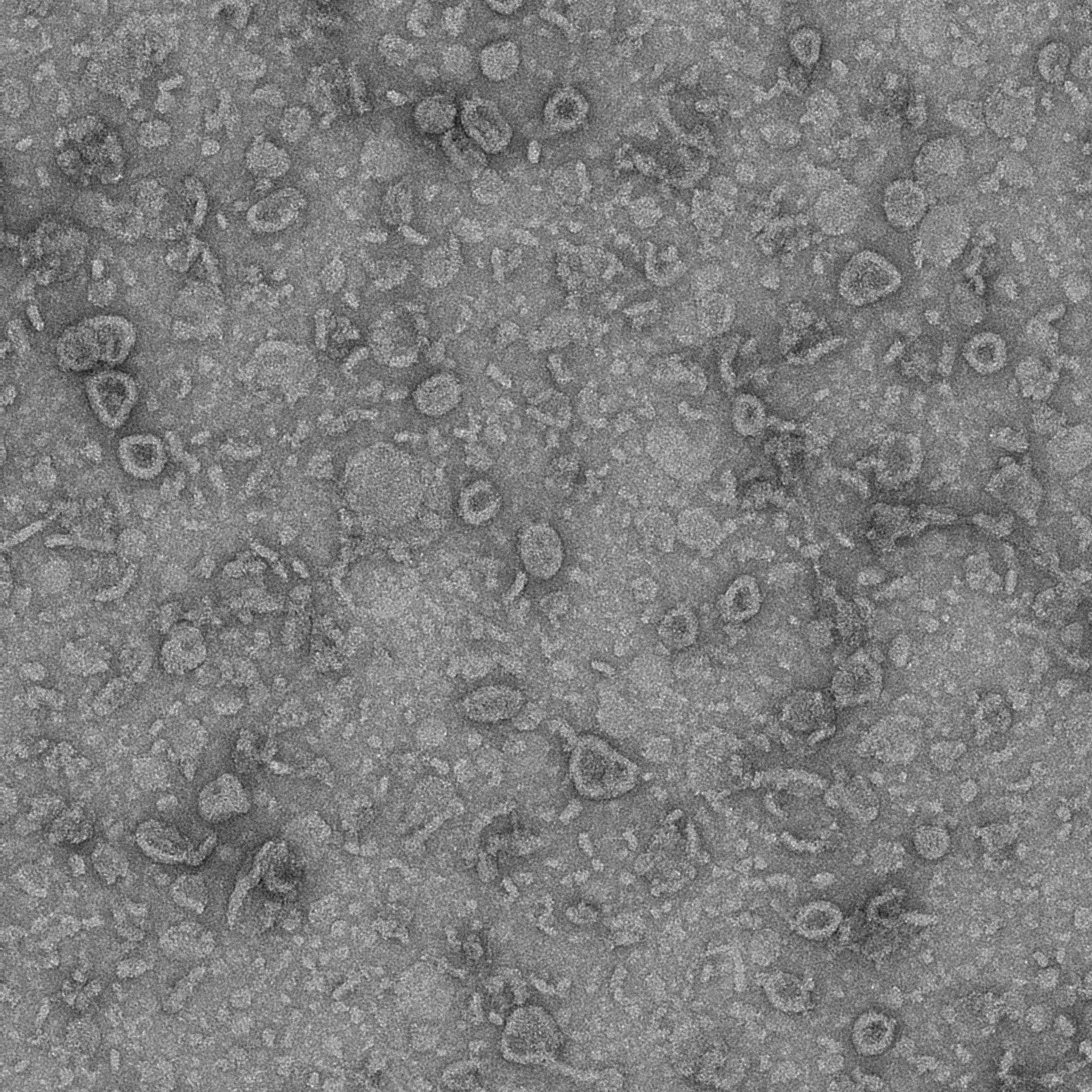

### 13B_2_1.pdf

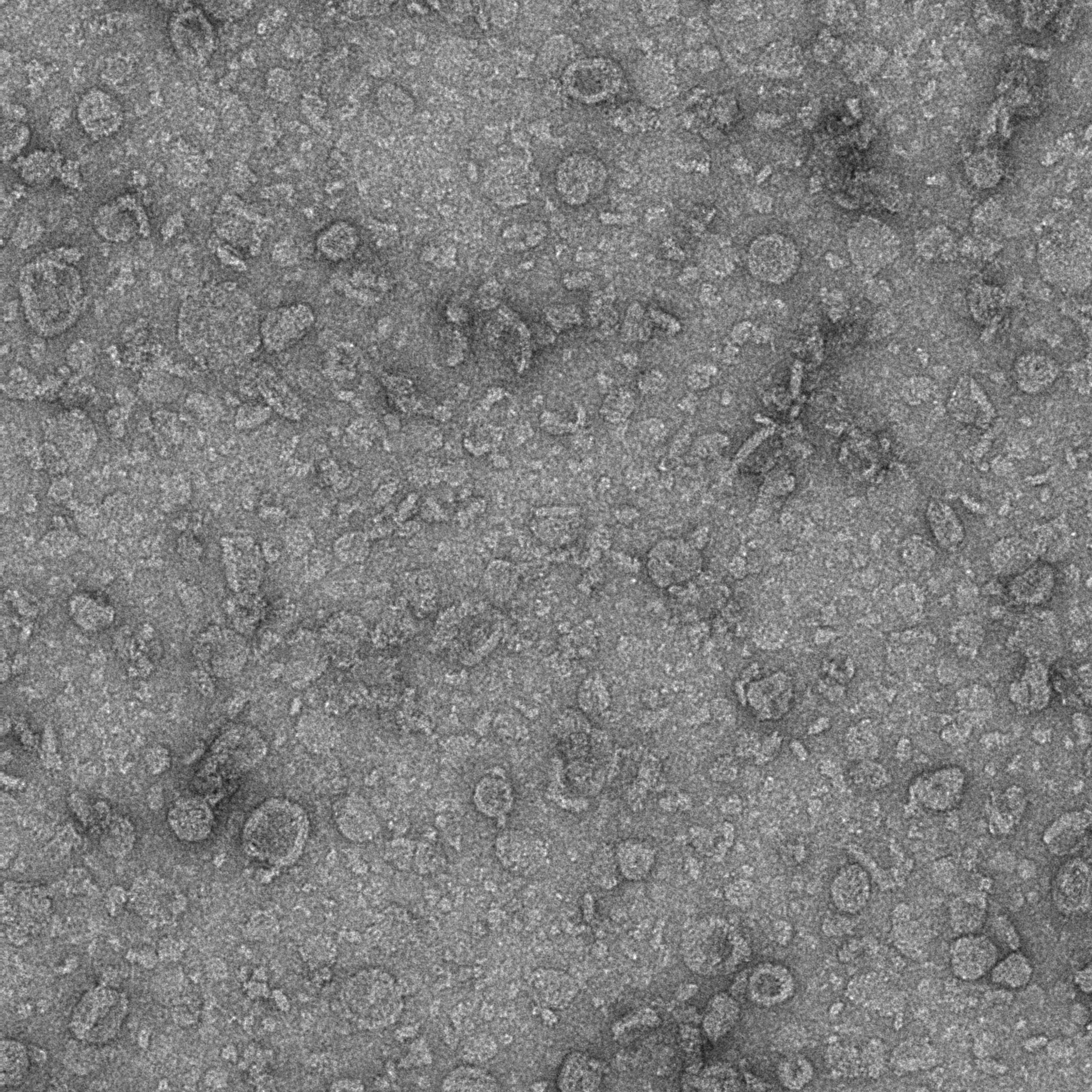

### 16A_0000_1.pdf

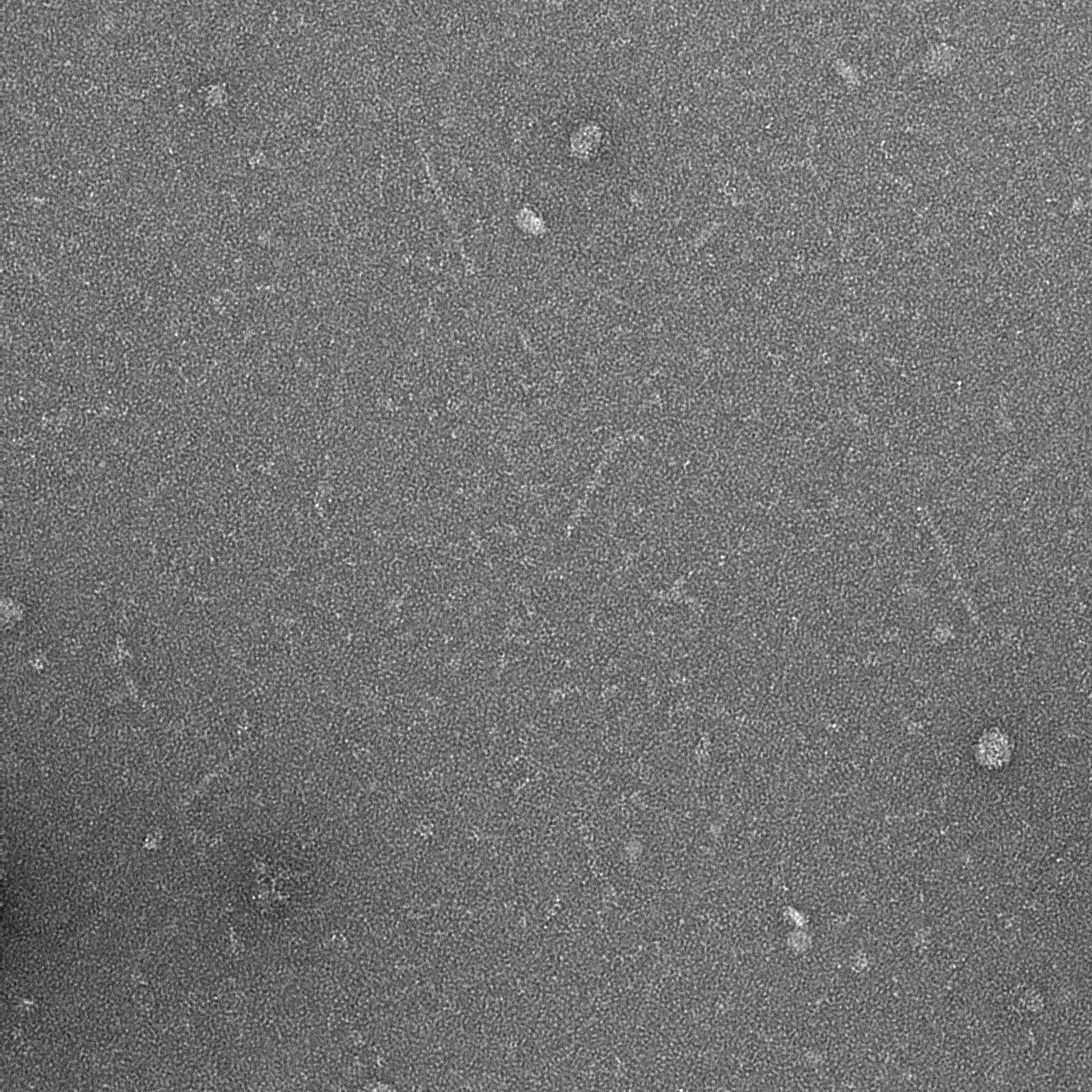

### 16A_0001_1.pdf

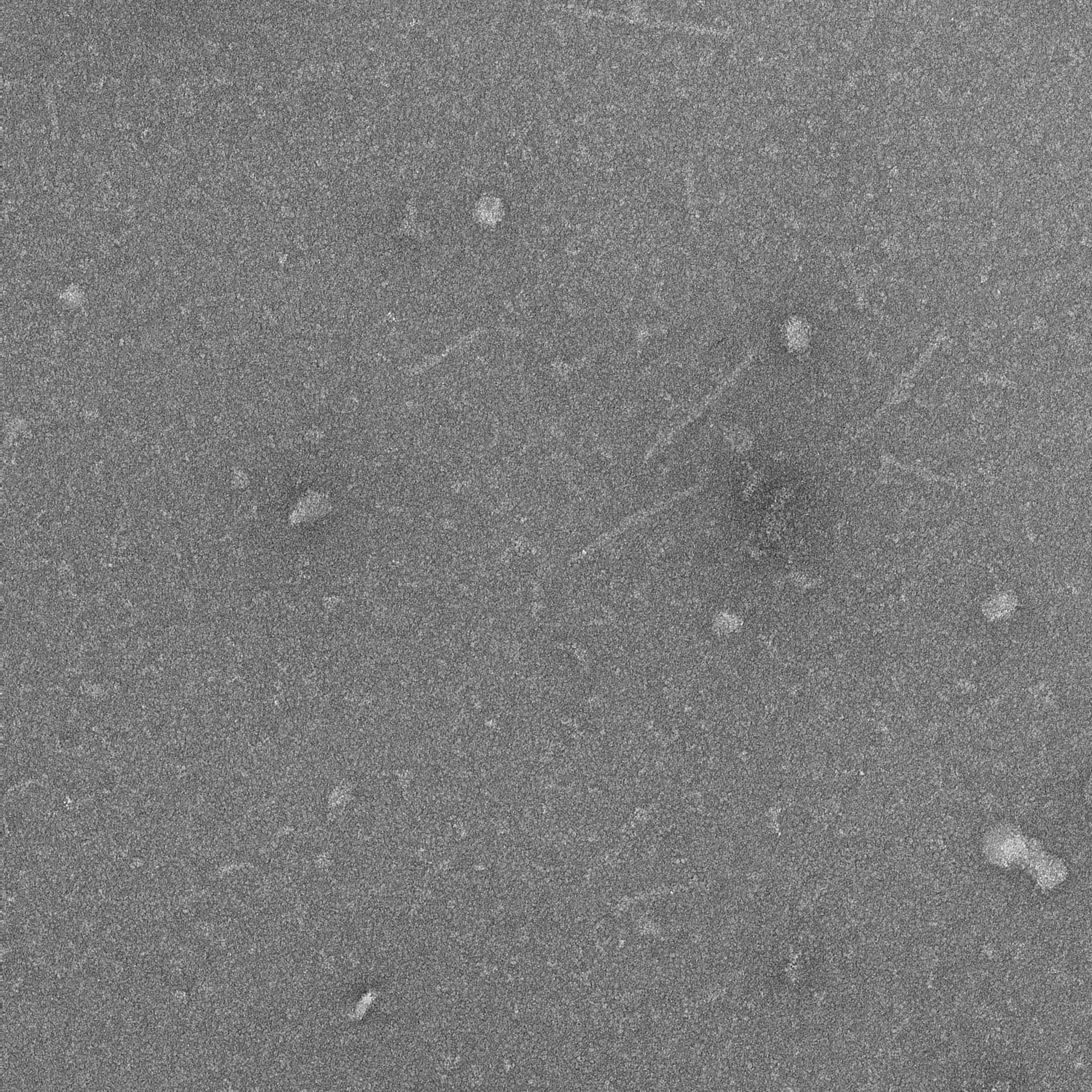

### 16A_0002_1.pdf

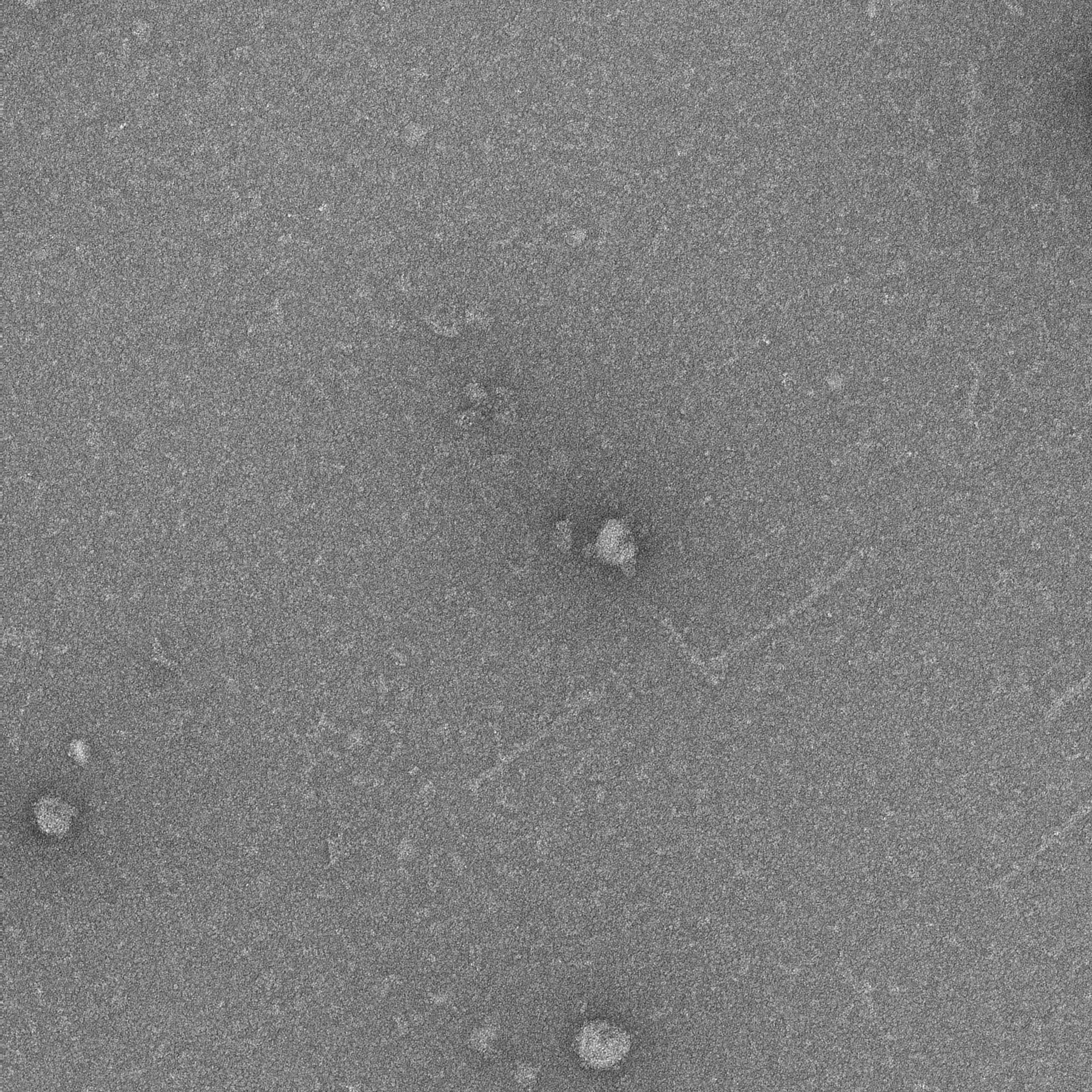

### 16A_0003_1.pdf

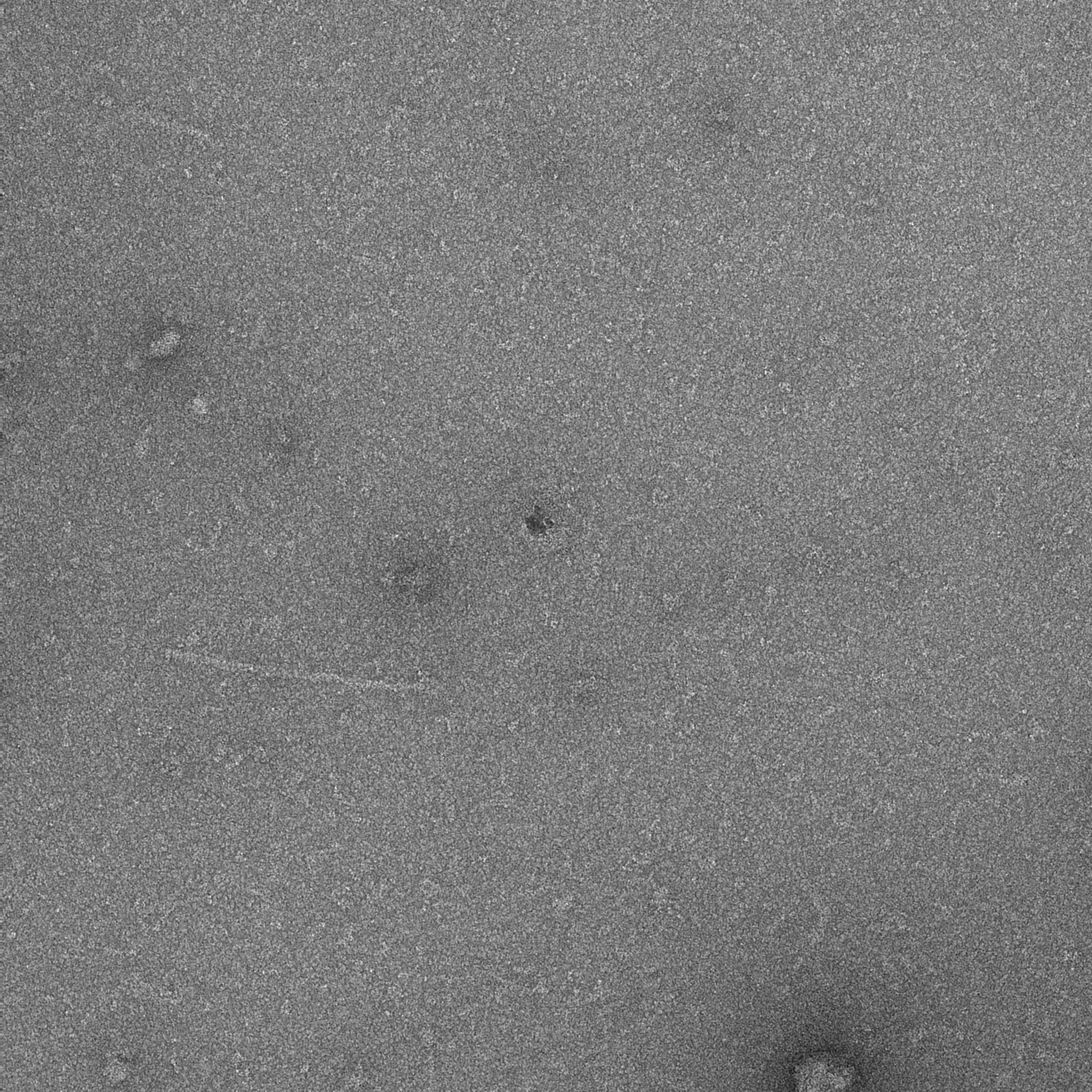

### 16A_0004_1.pdf

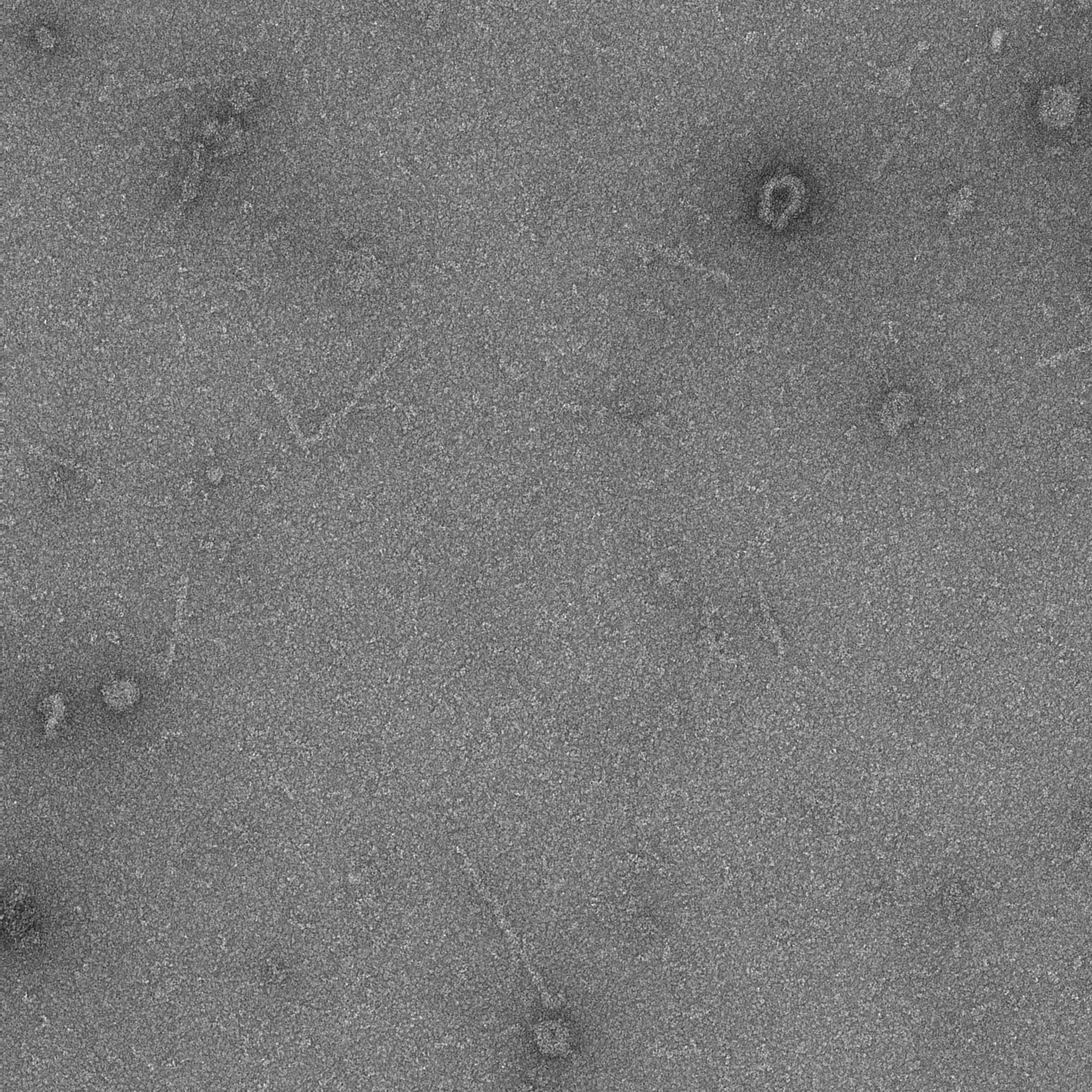

### 16A_0005_1.pdf

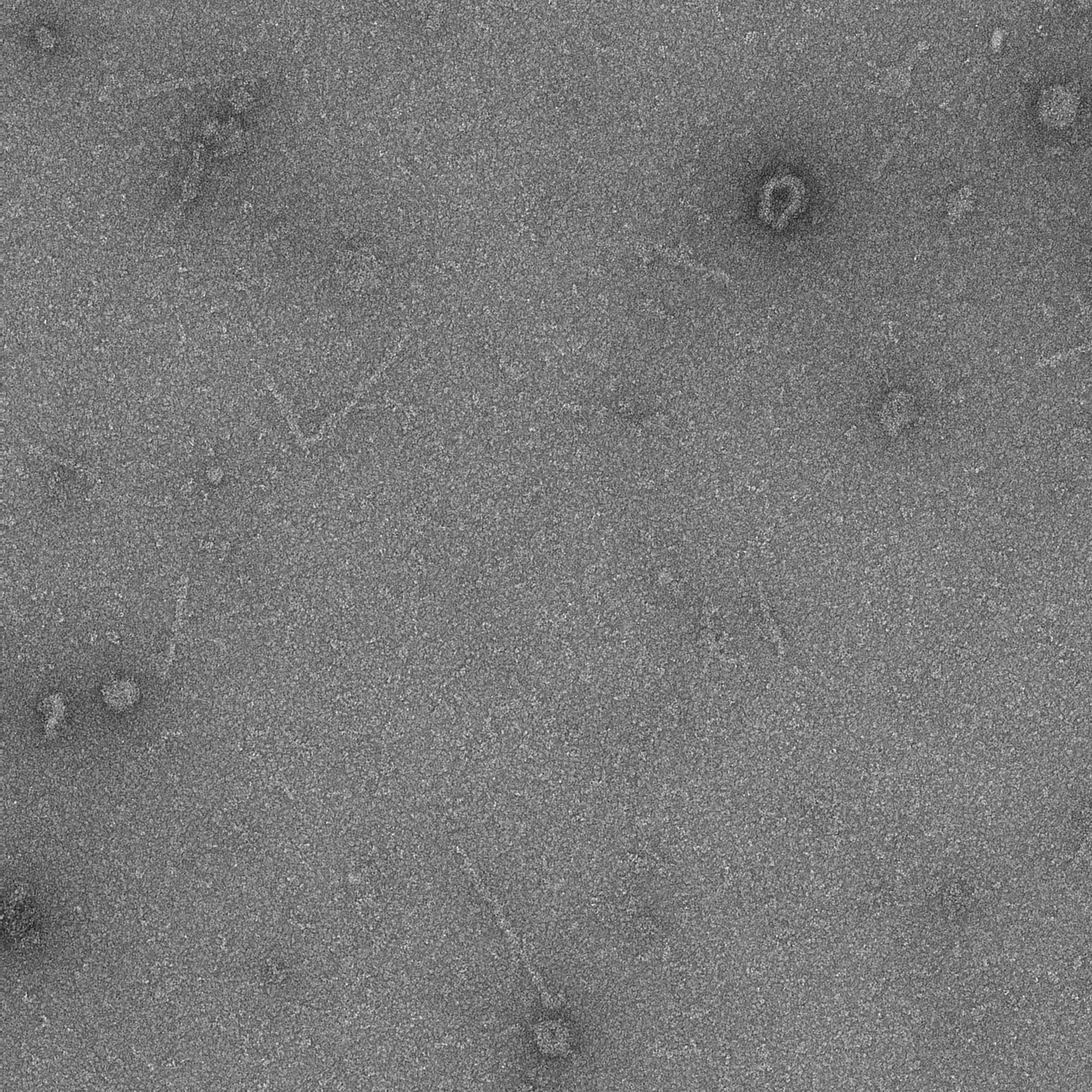

### 16A_0006_1.pdf

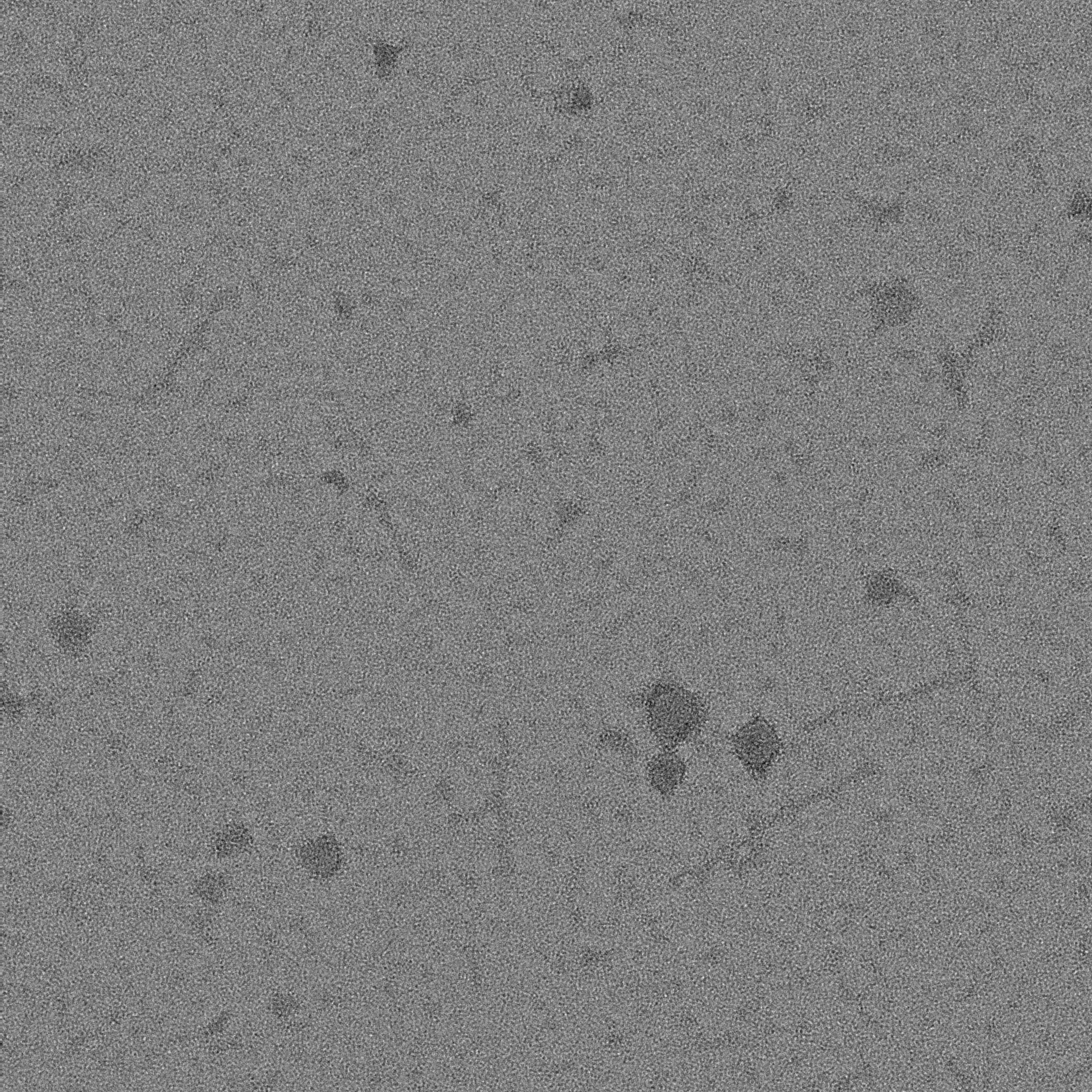

### 16A_0007_1.pdf

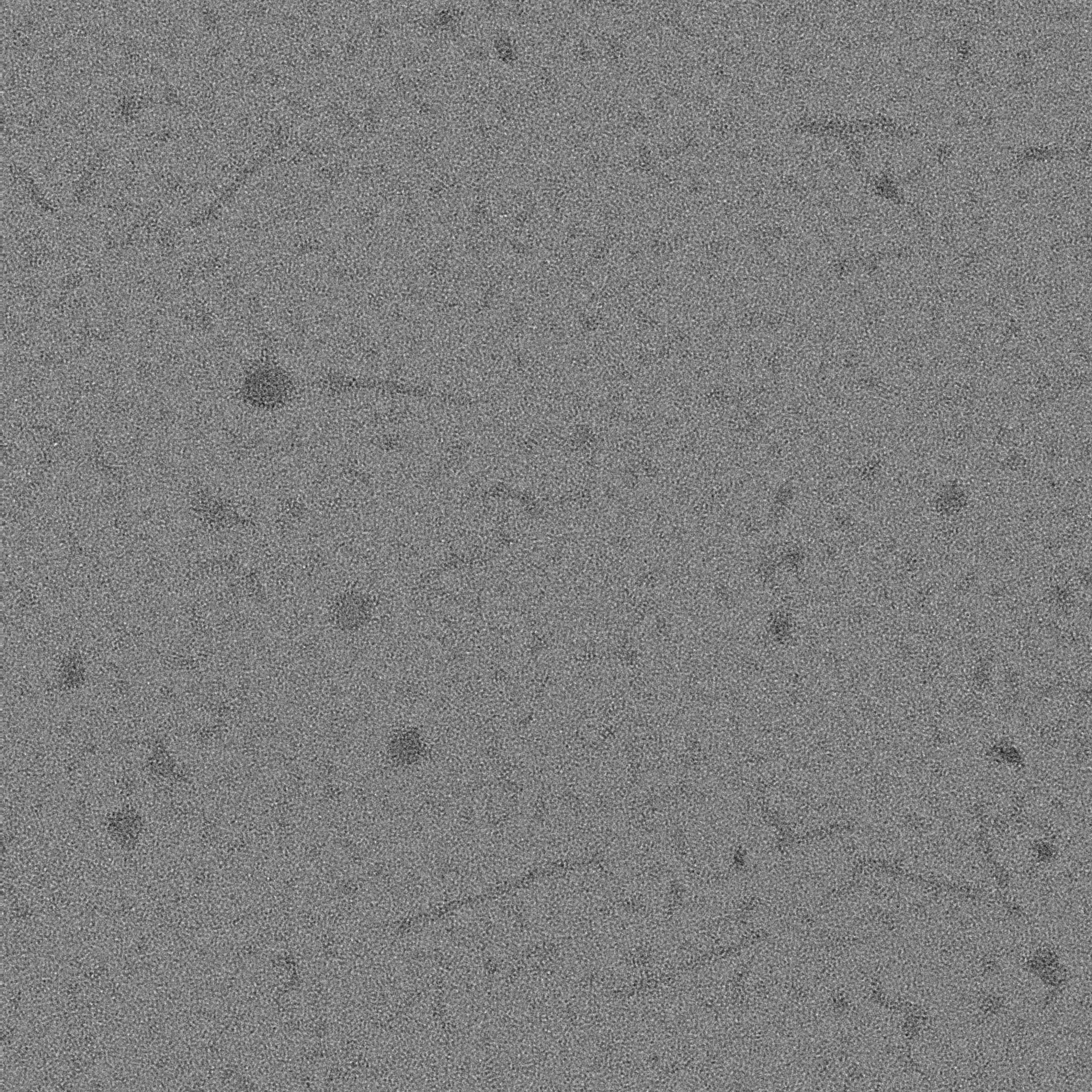

### 16A_0008_1.pdf

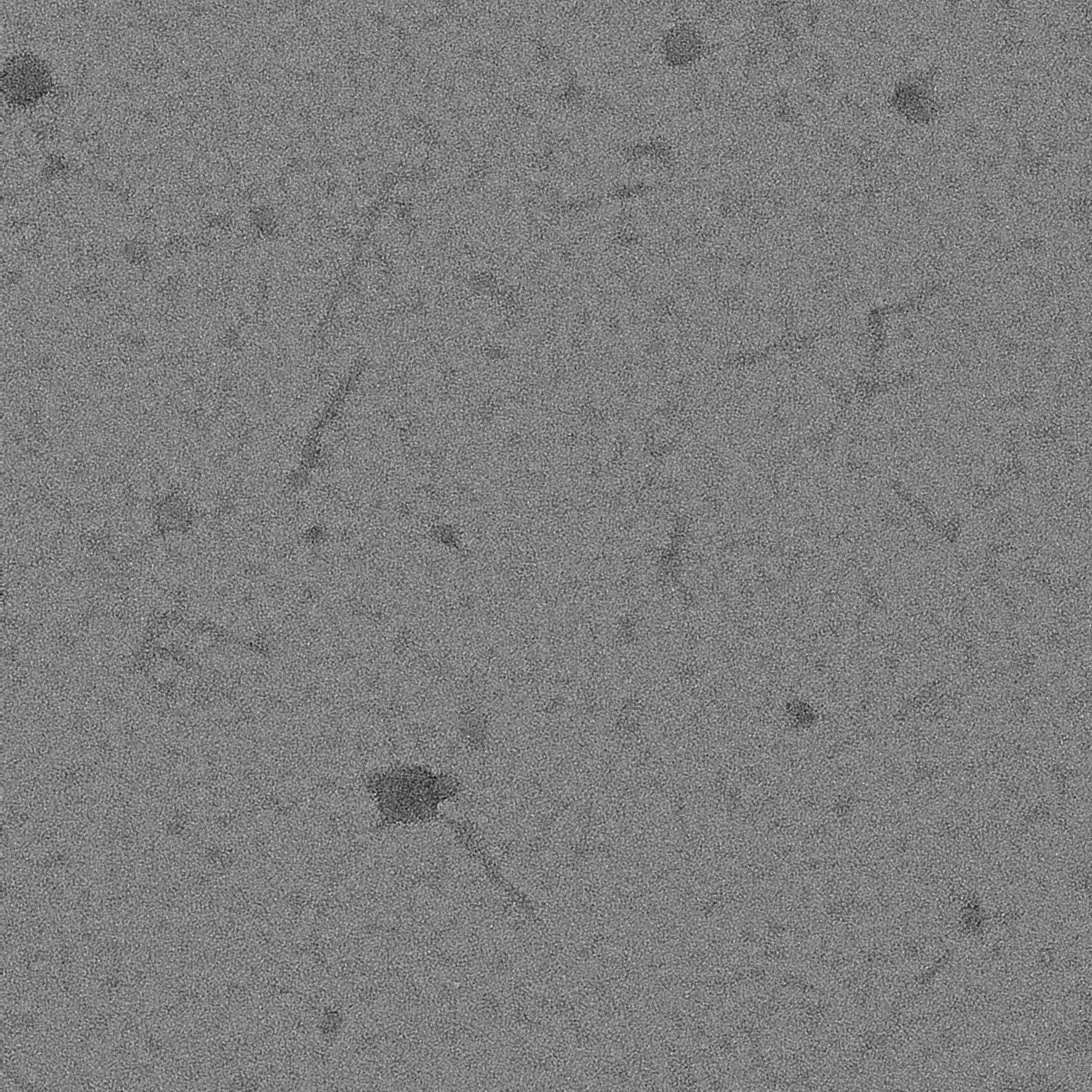

### 16A_0009_1.pdf

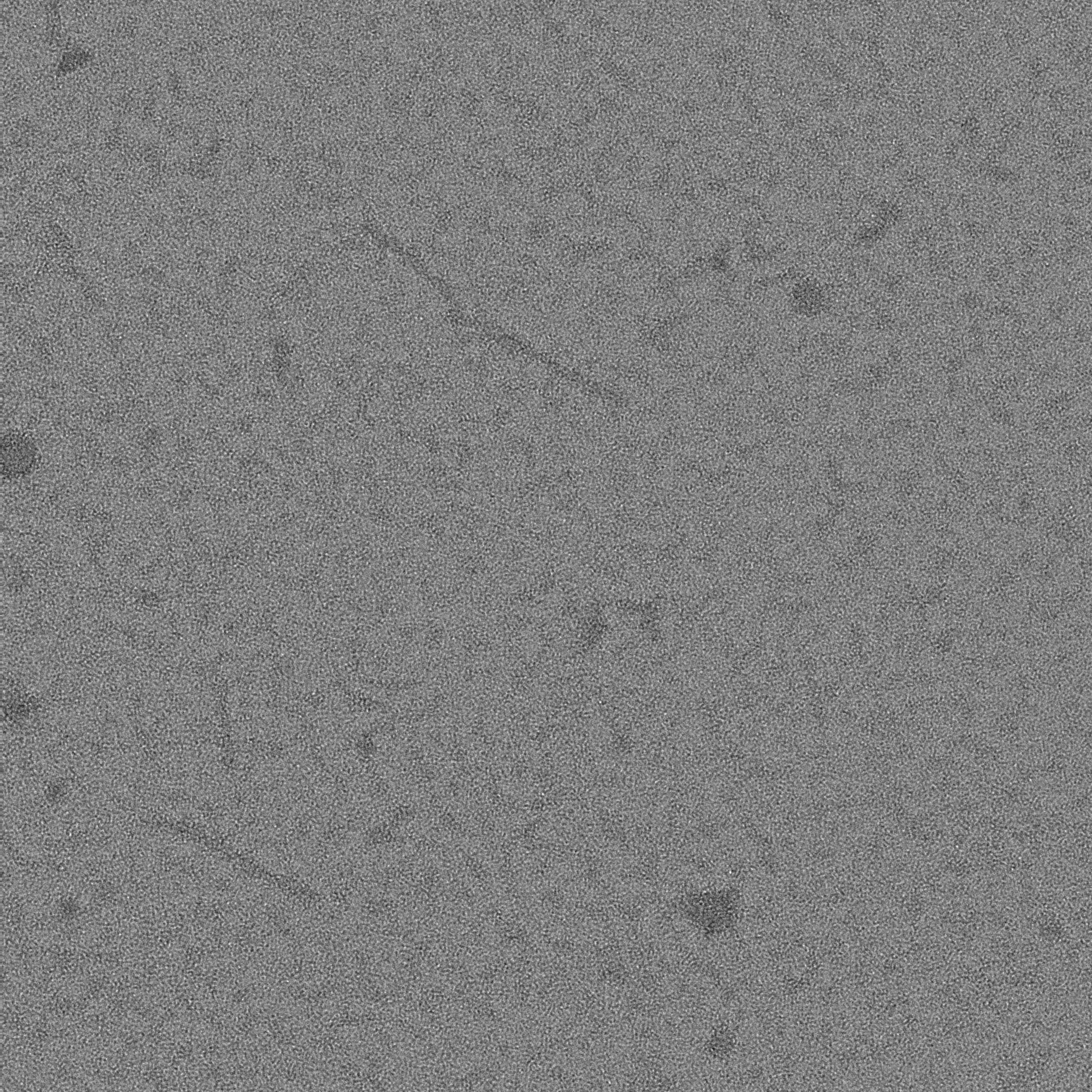

### 16A_0010_1.pdf

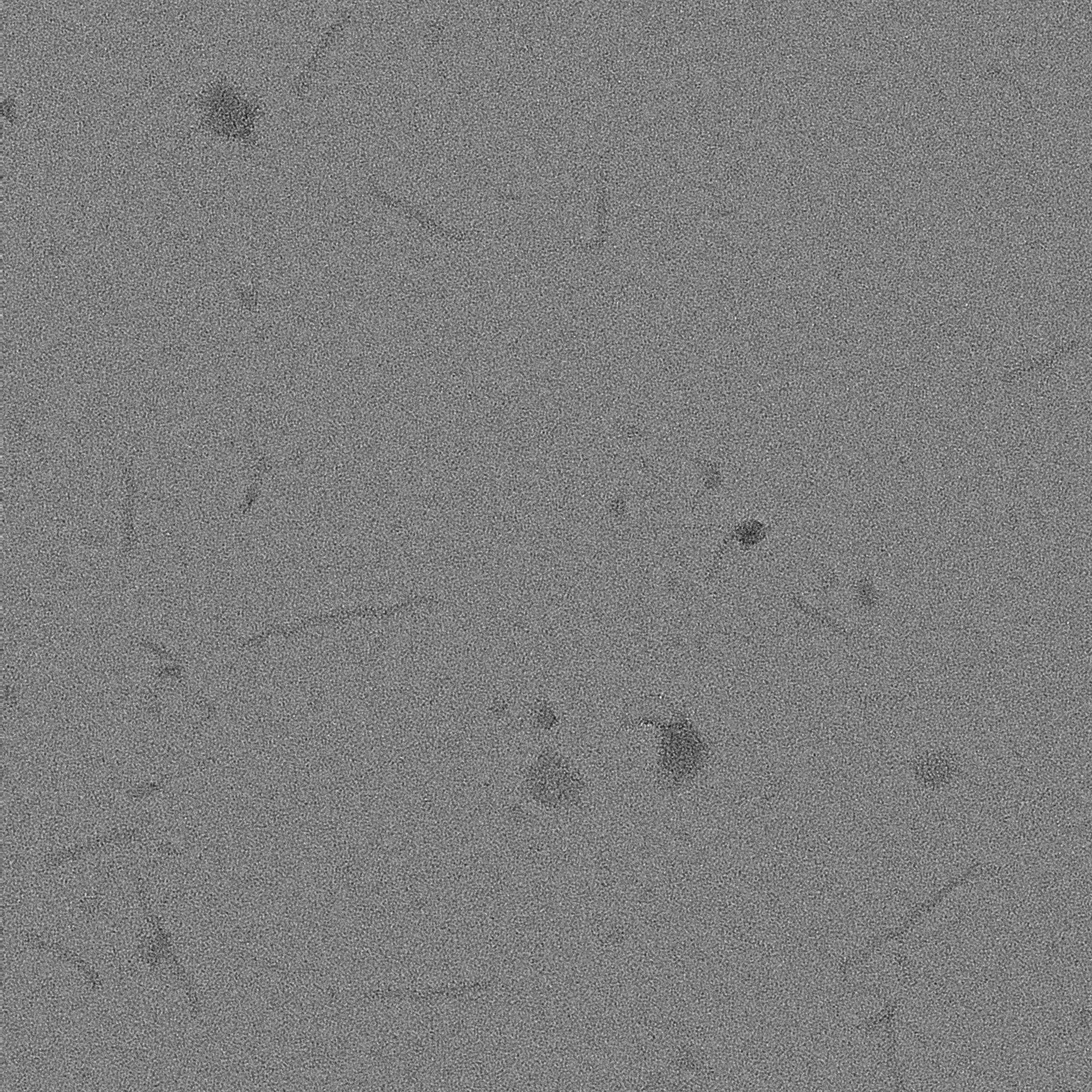

### 16A_0011_1.pdf

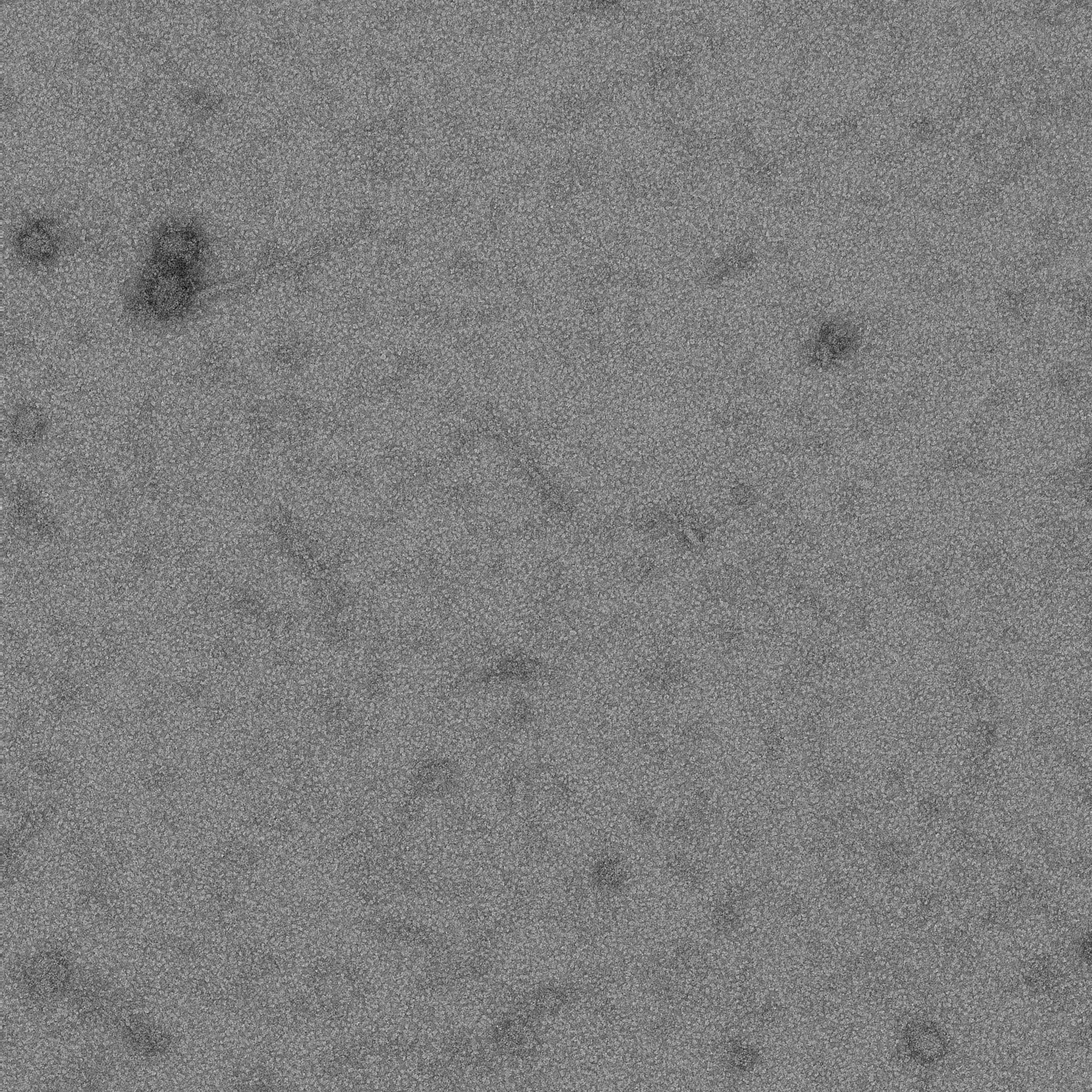

### 16A_0012_1.pdf

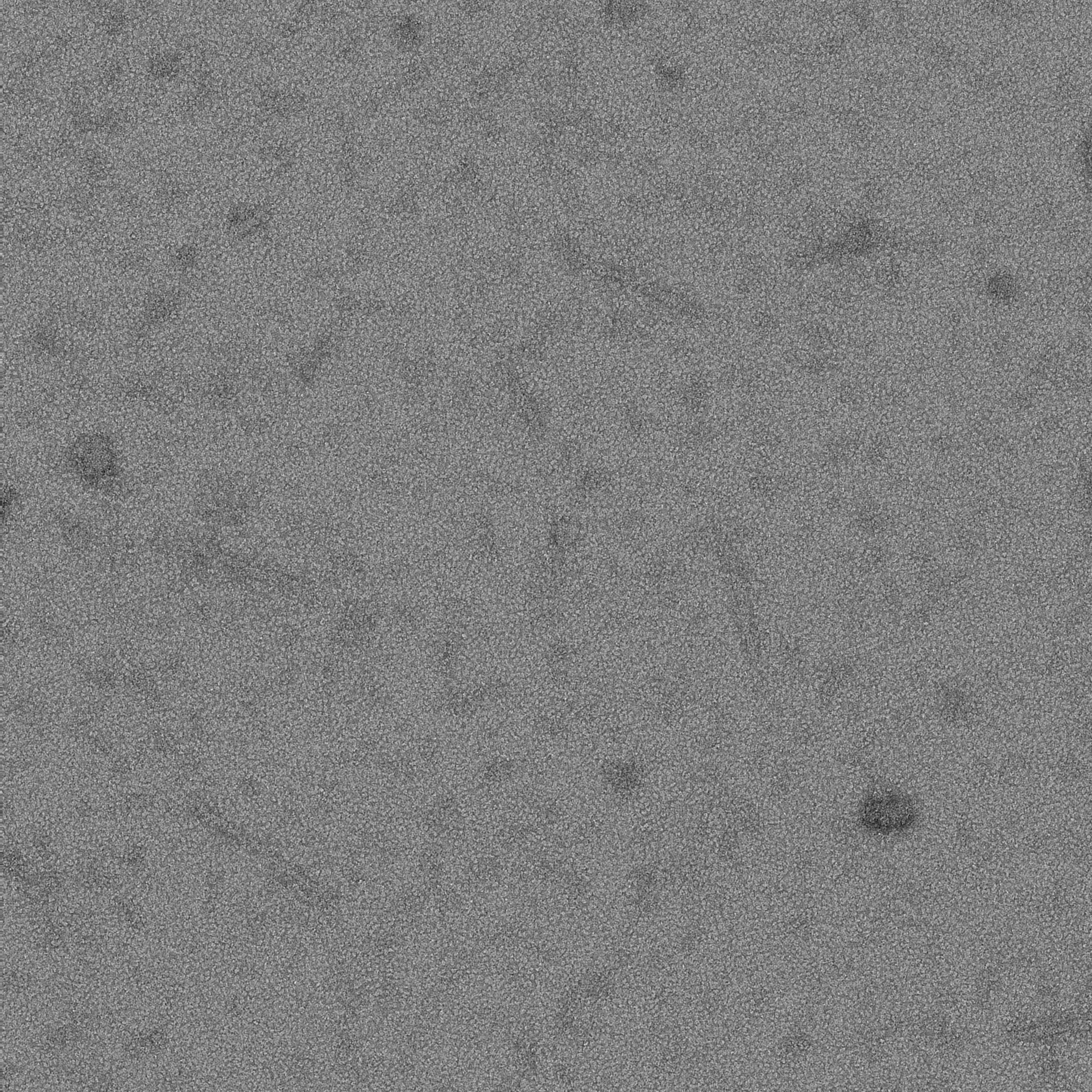

### 16A_0013_1.pdf

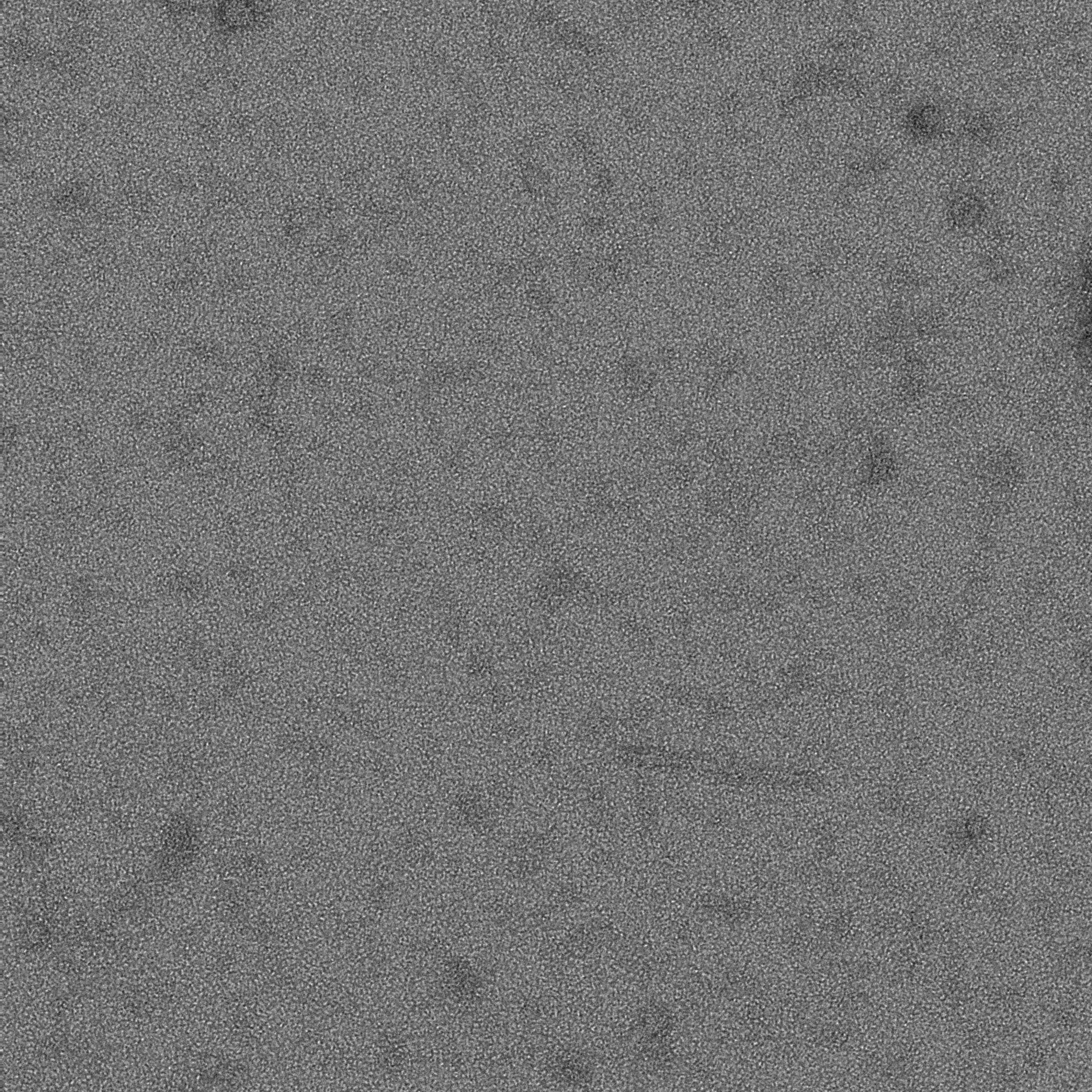

### 16A_0014_1.pdf

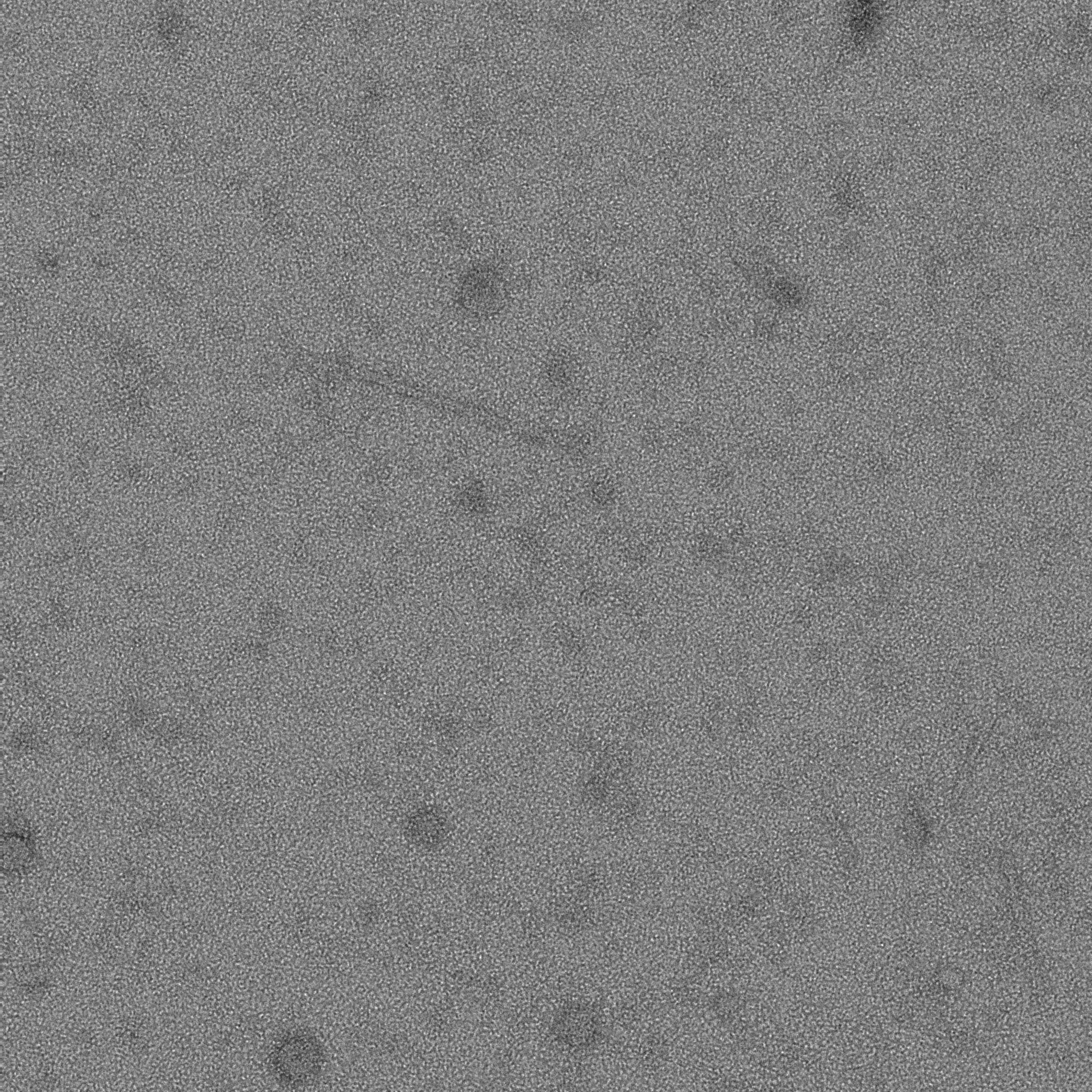

### 16A_0015_1.pdf

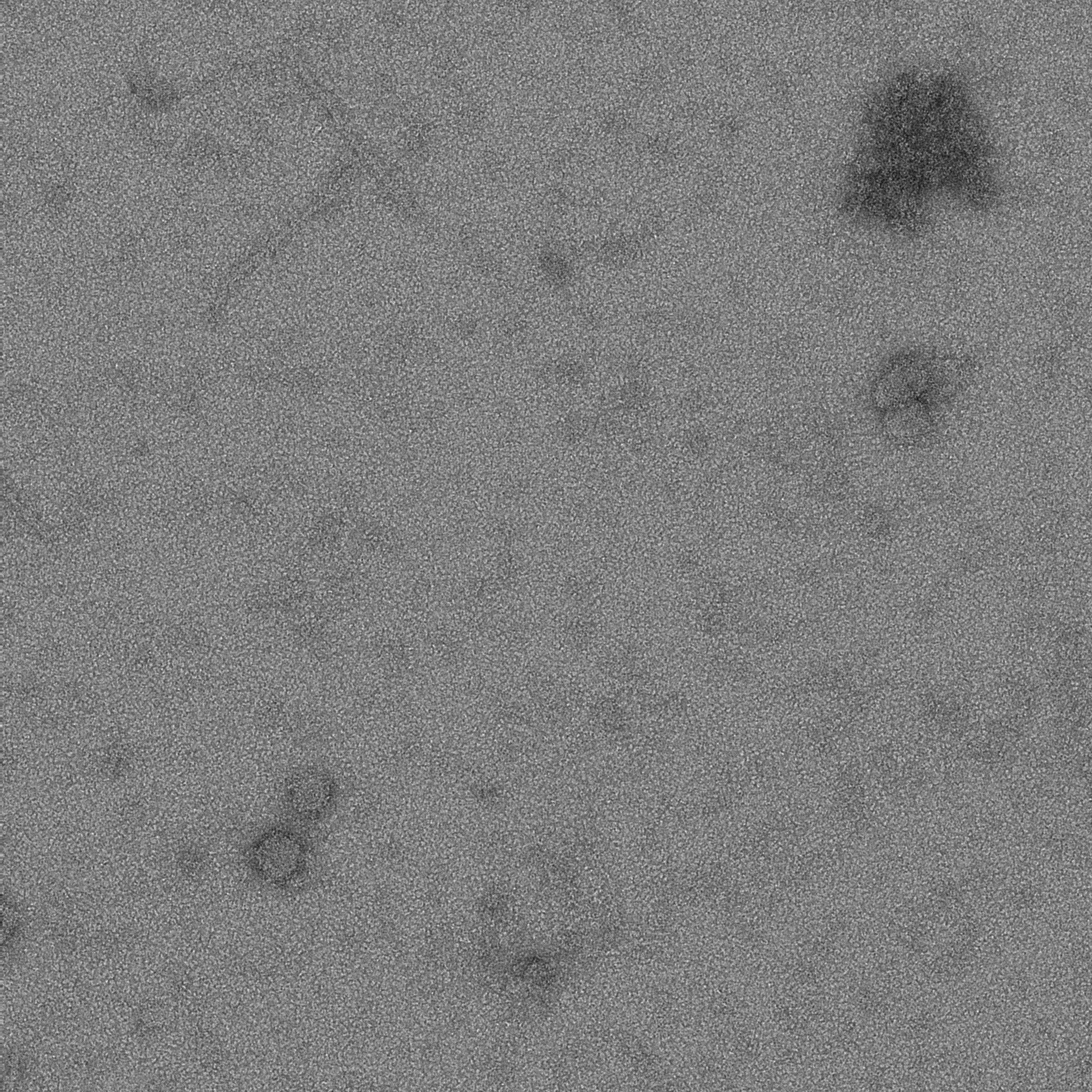

### 16A_0016_1.pdf

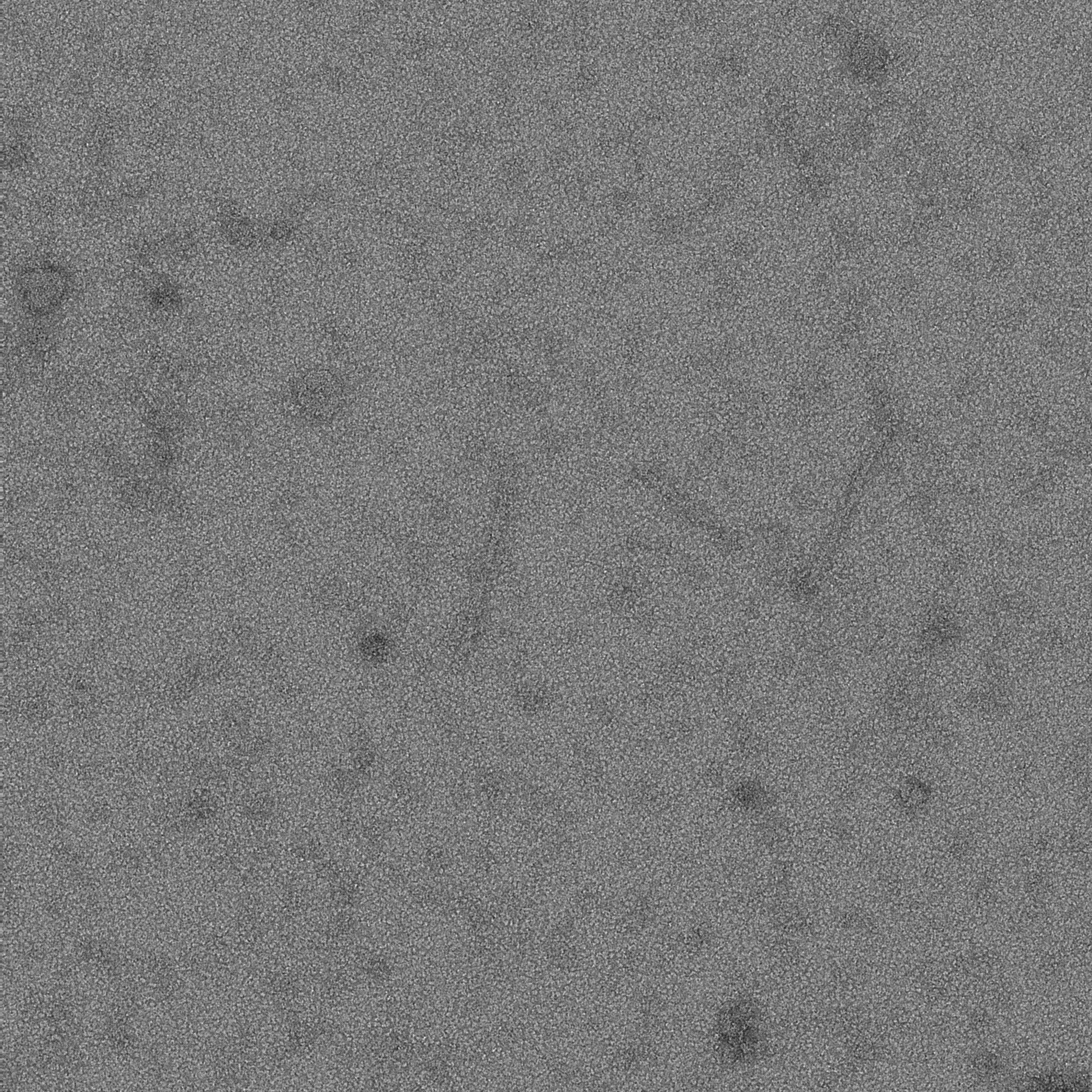

### 16C_0000_1.pdf

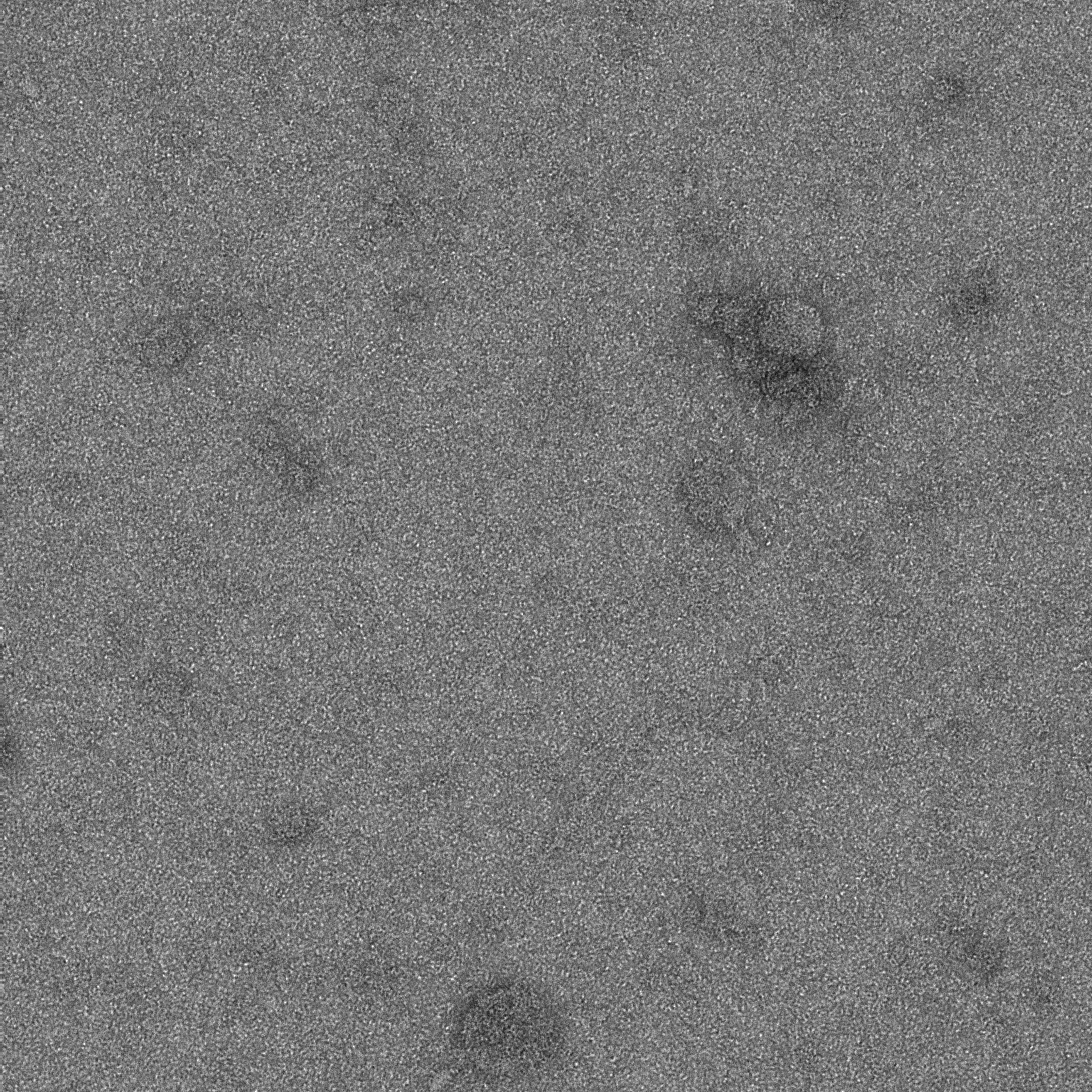

### 16C_0001_1.pdf

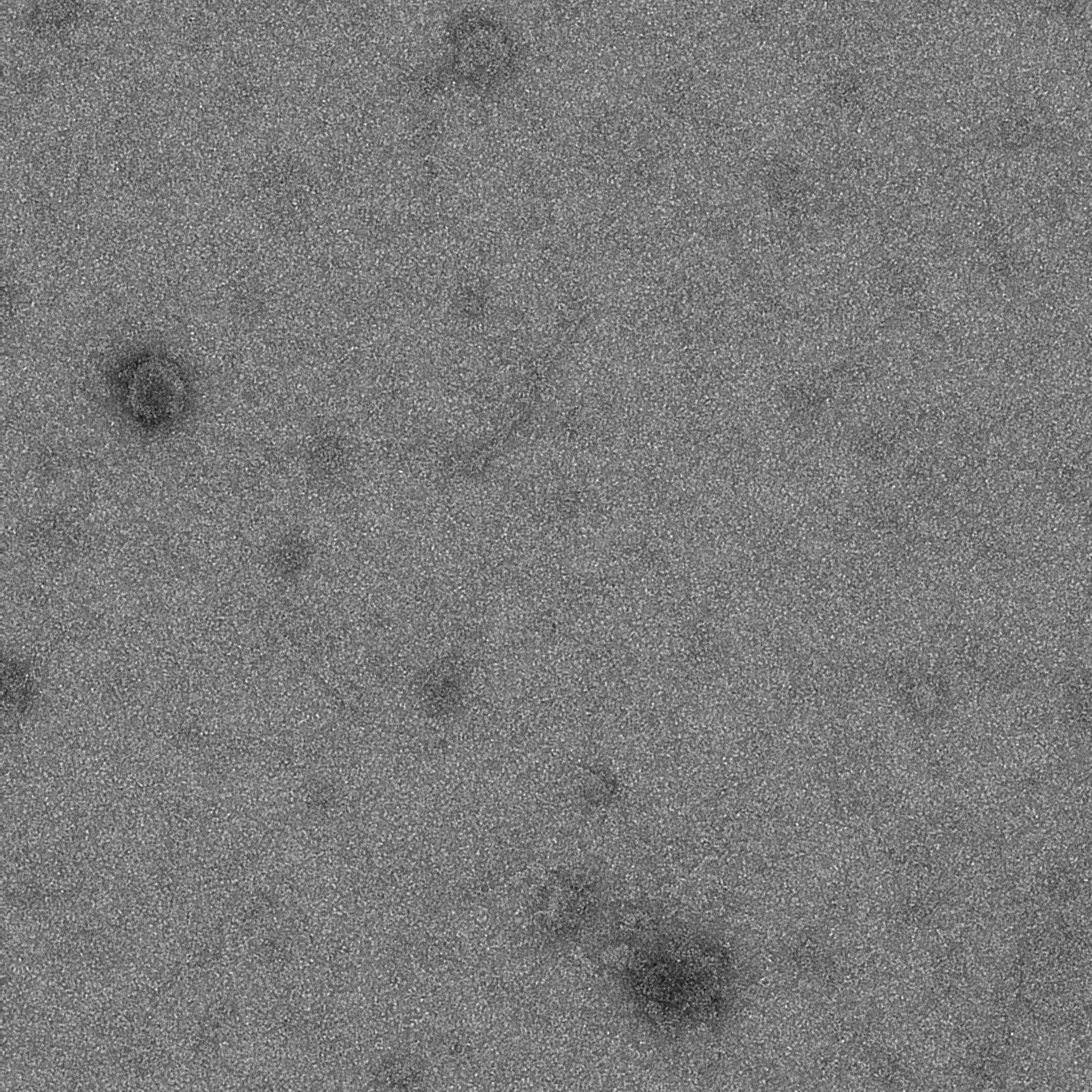

### 16C_0002_1.pdf

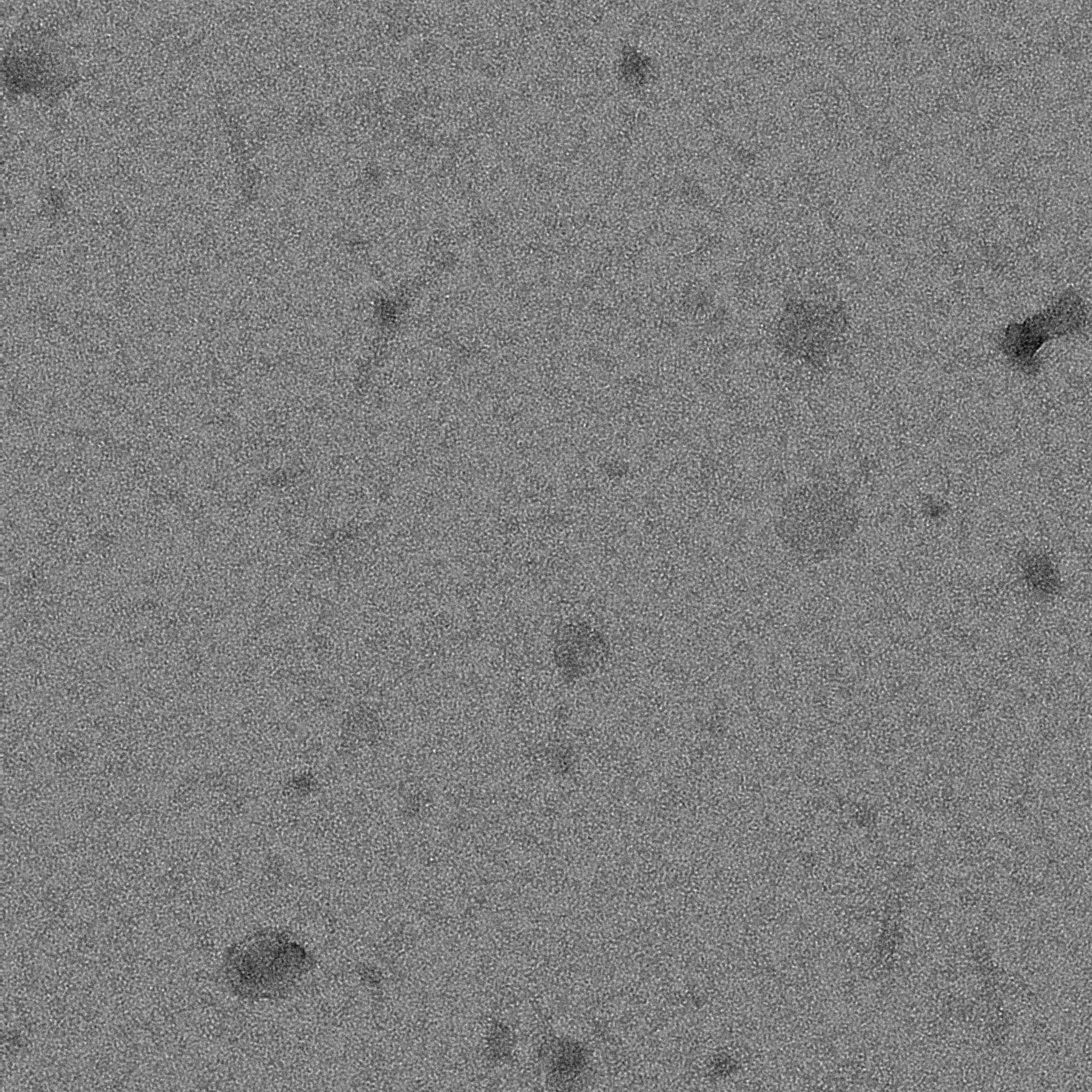

### 16C_0003_1.pdf

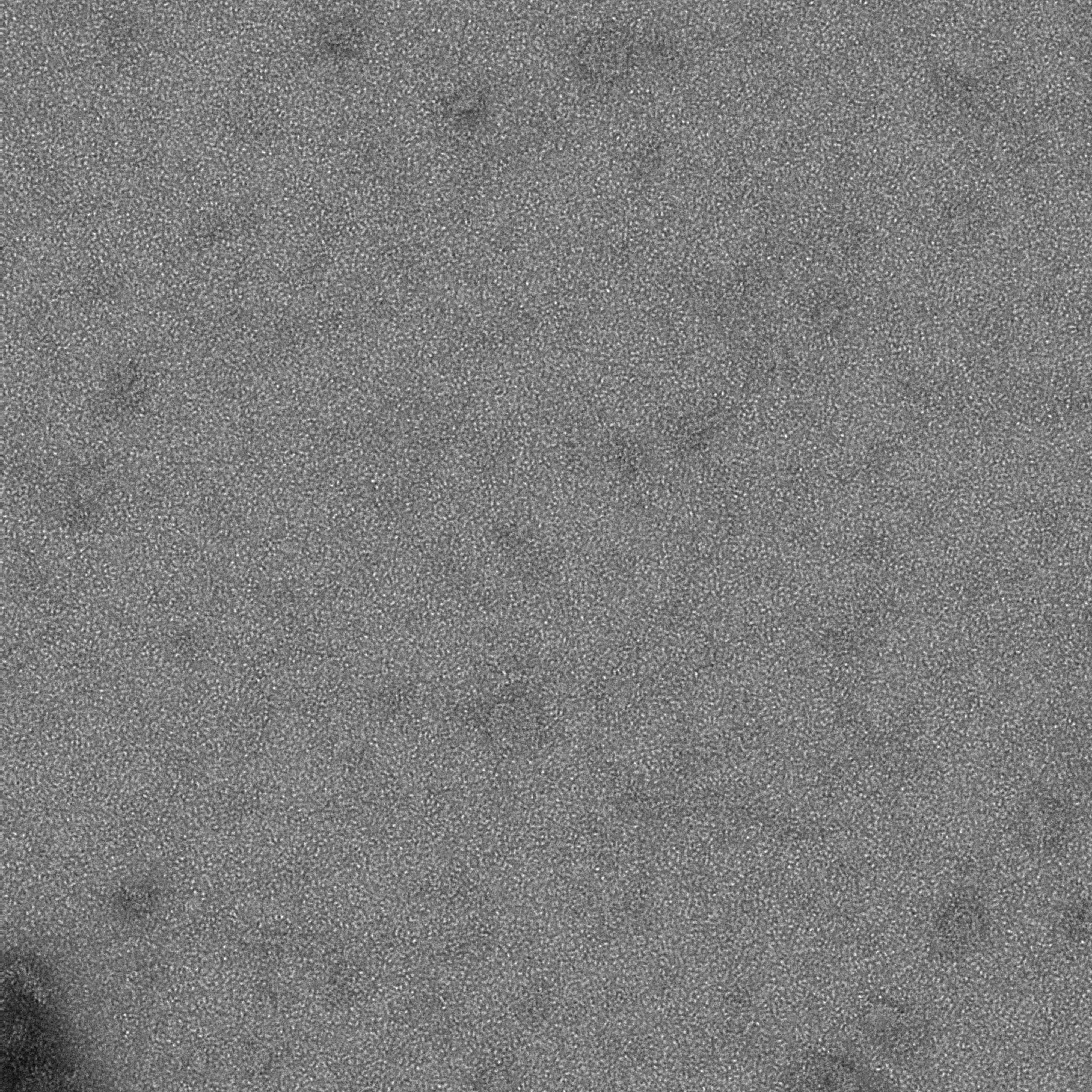

### 16C_0004_1.pdf

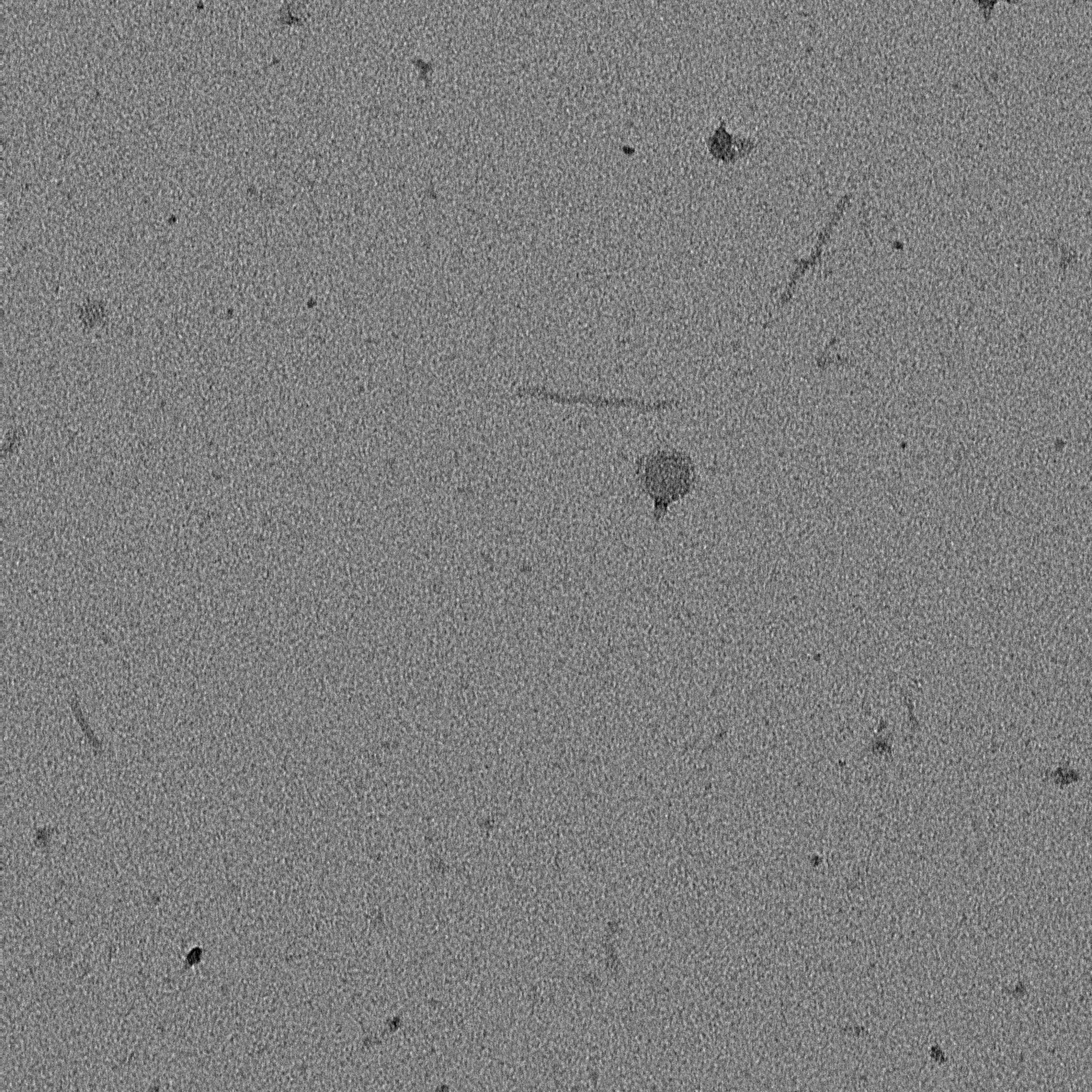

### 16C_0005_1.pdf

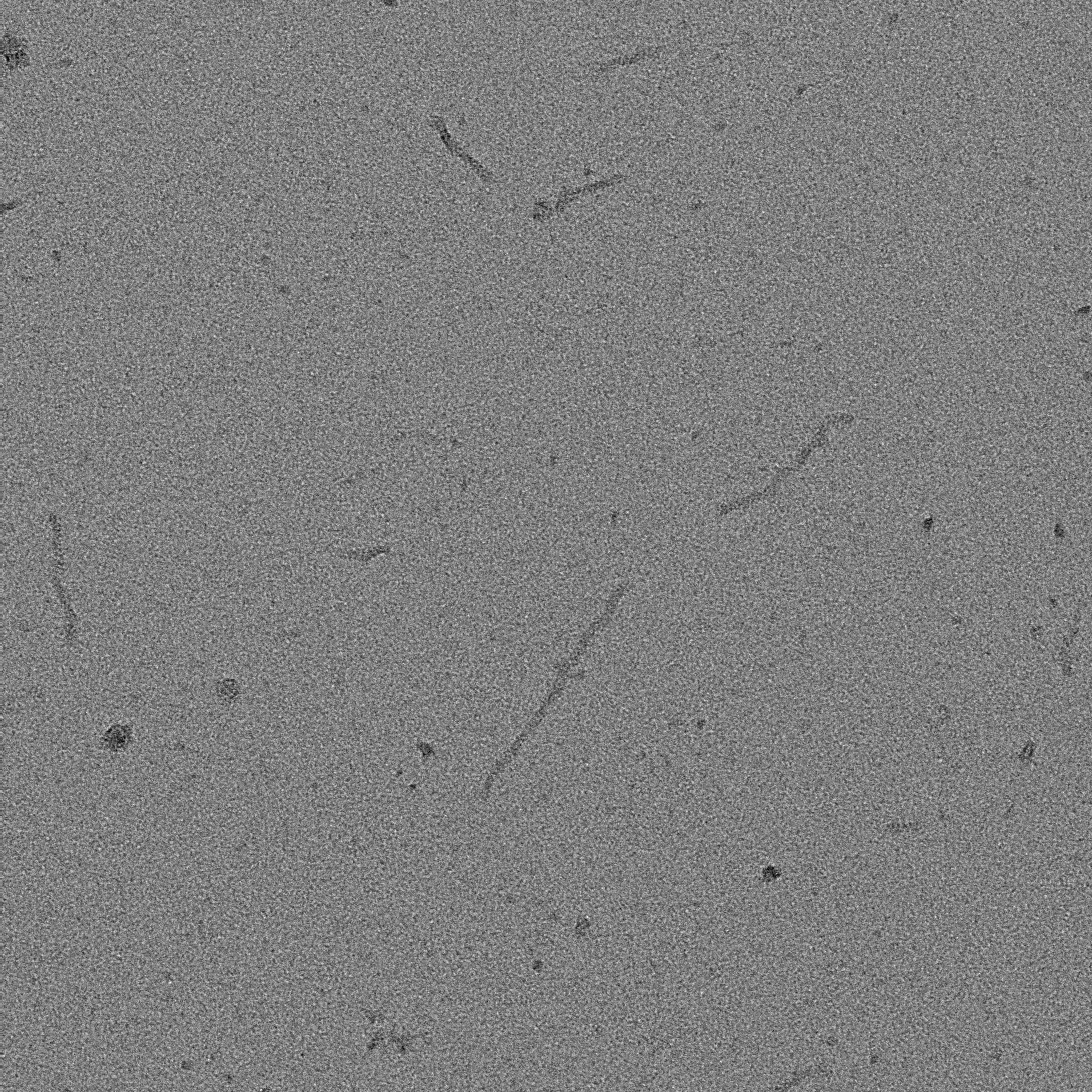

### 16C_0006_1.pdf

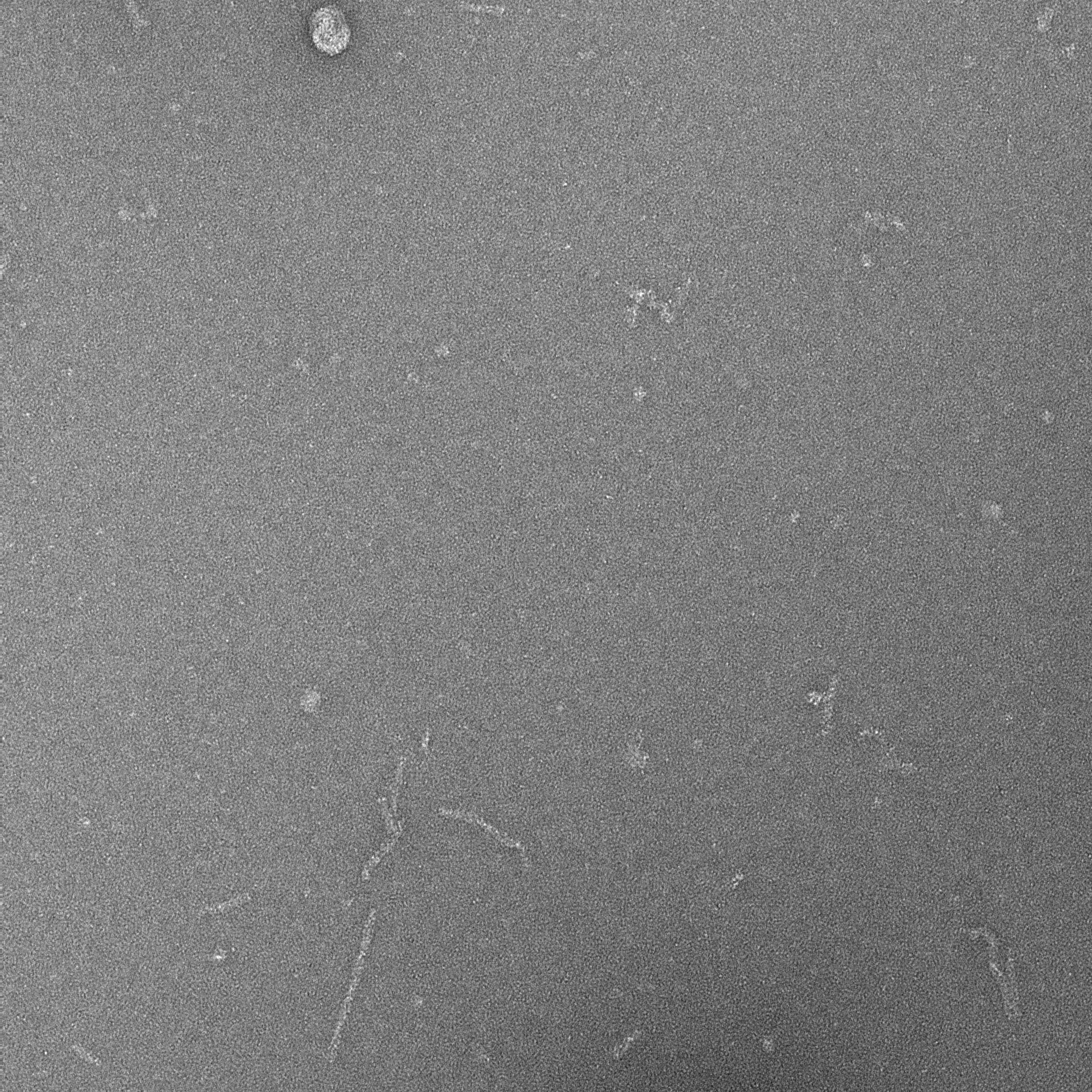

### 16C_0007_1.pdf

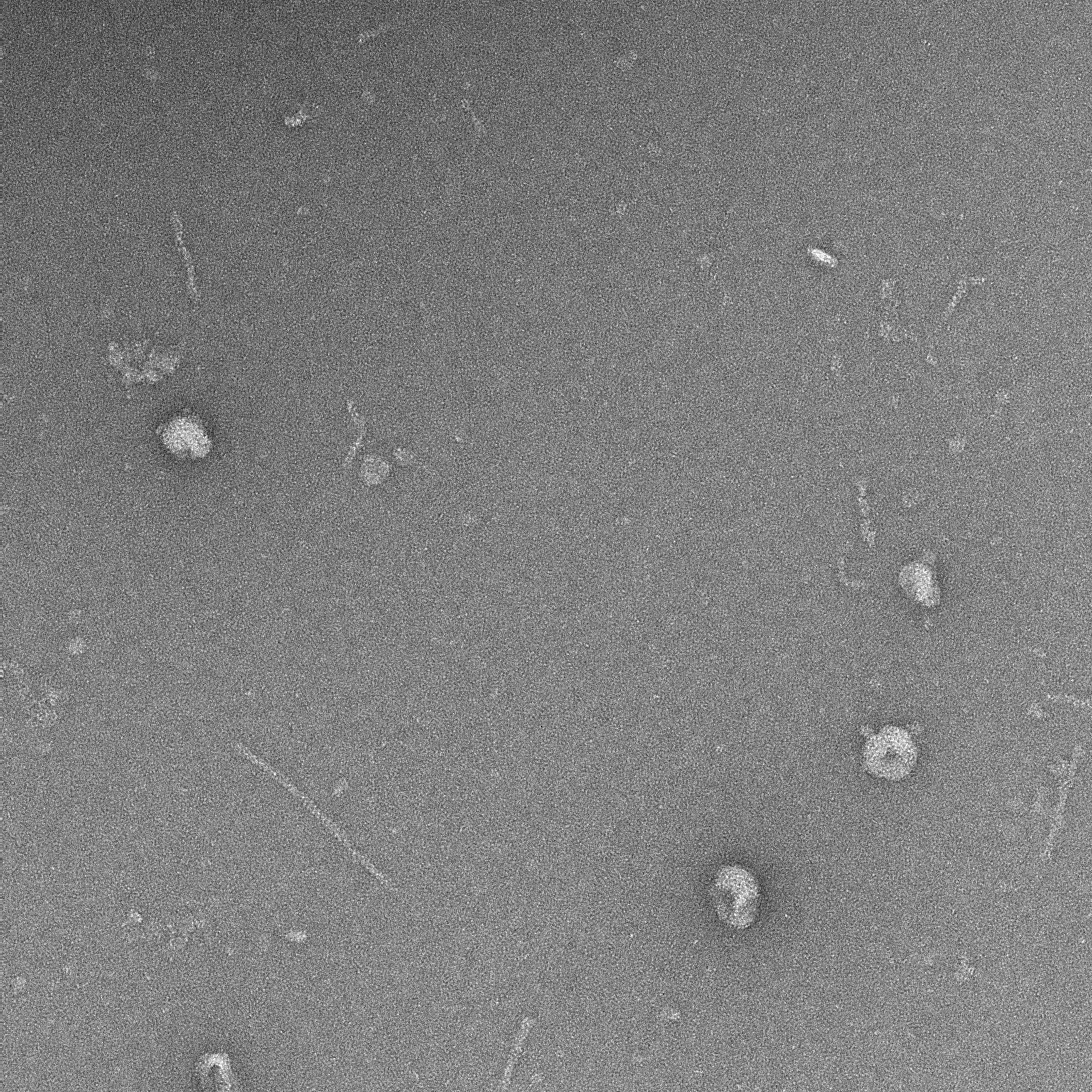

### 16C_0008_1.pdf

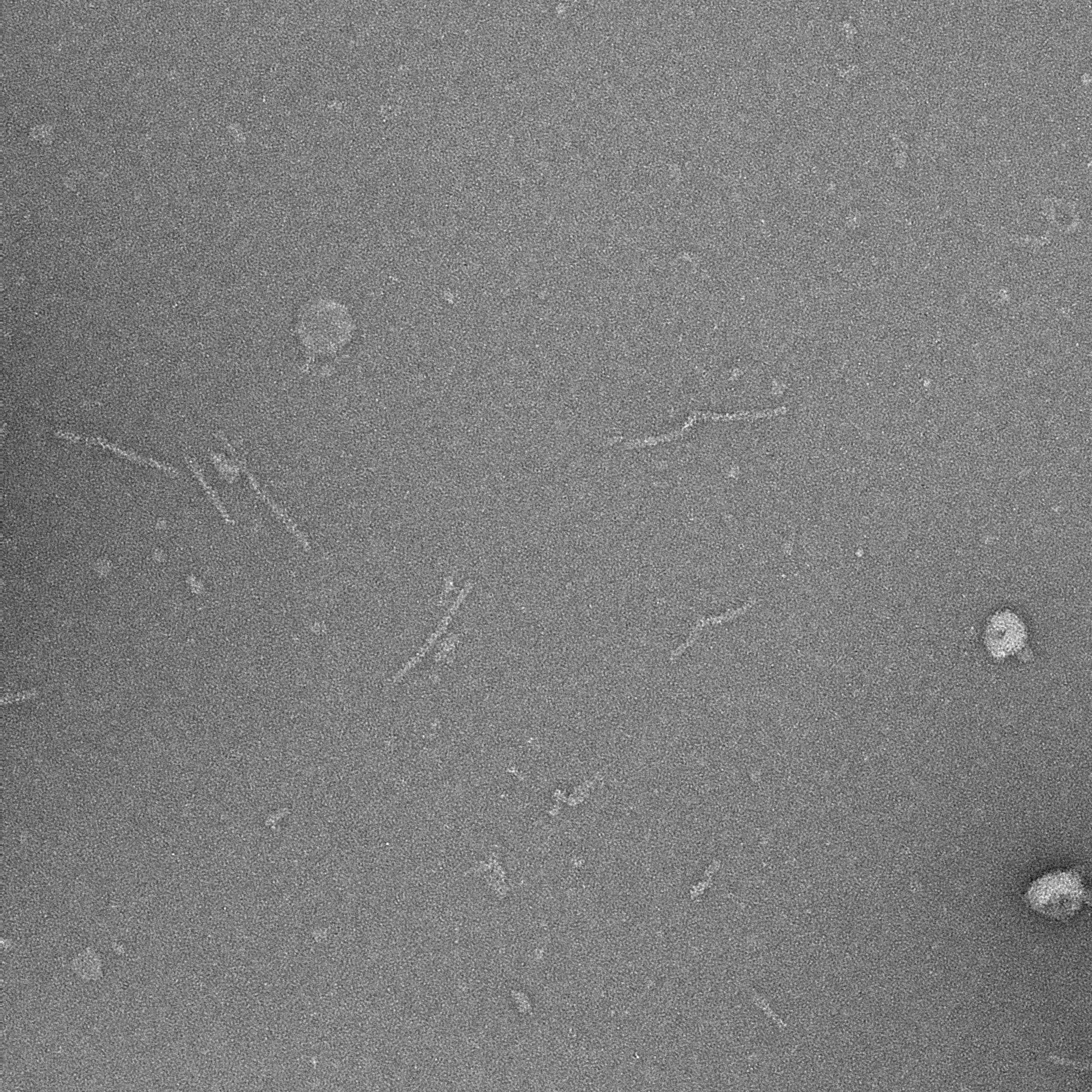

### 16C_0009_1.pdf

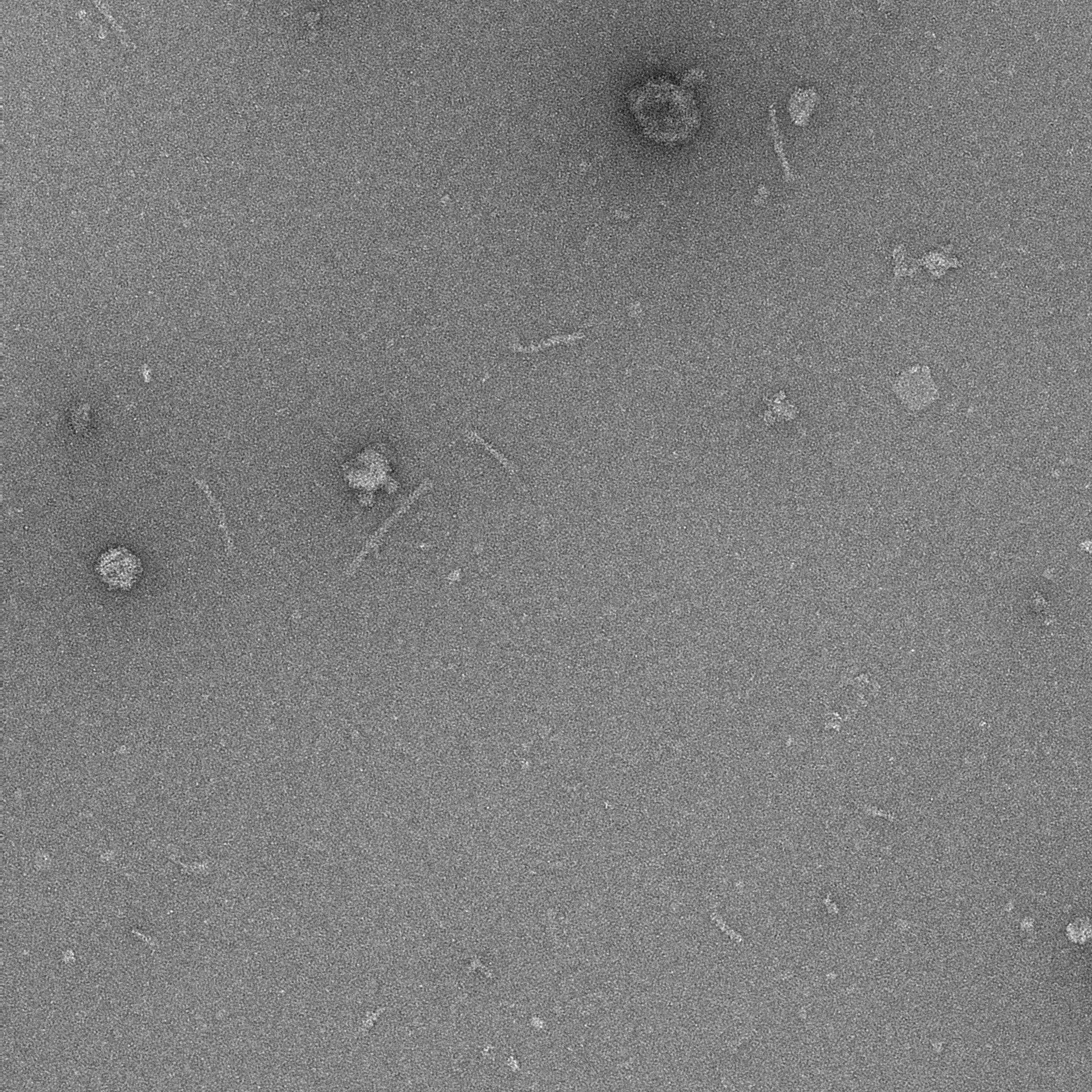

### 16E_0000_1.pdf

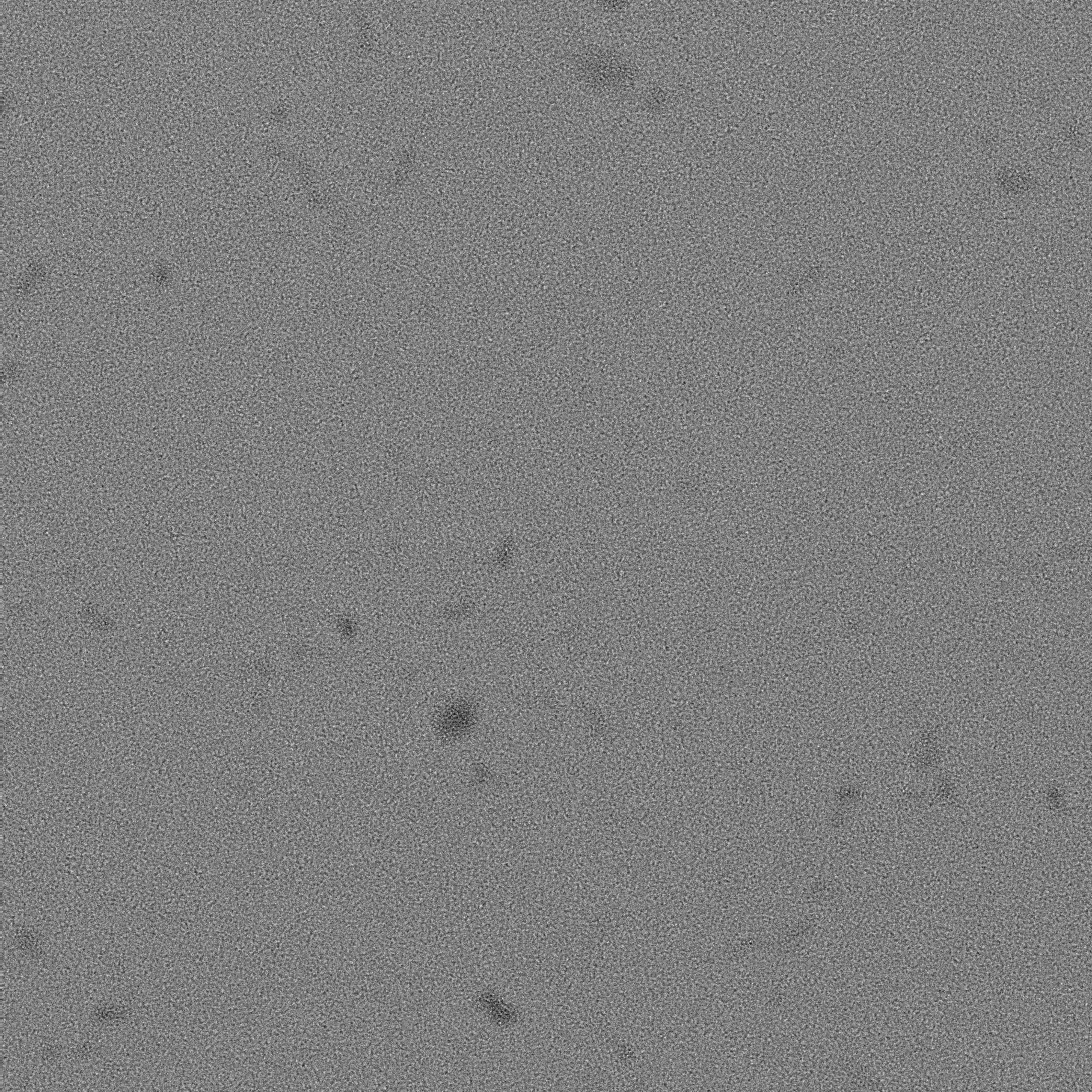

### 16E_0001_1.pdf

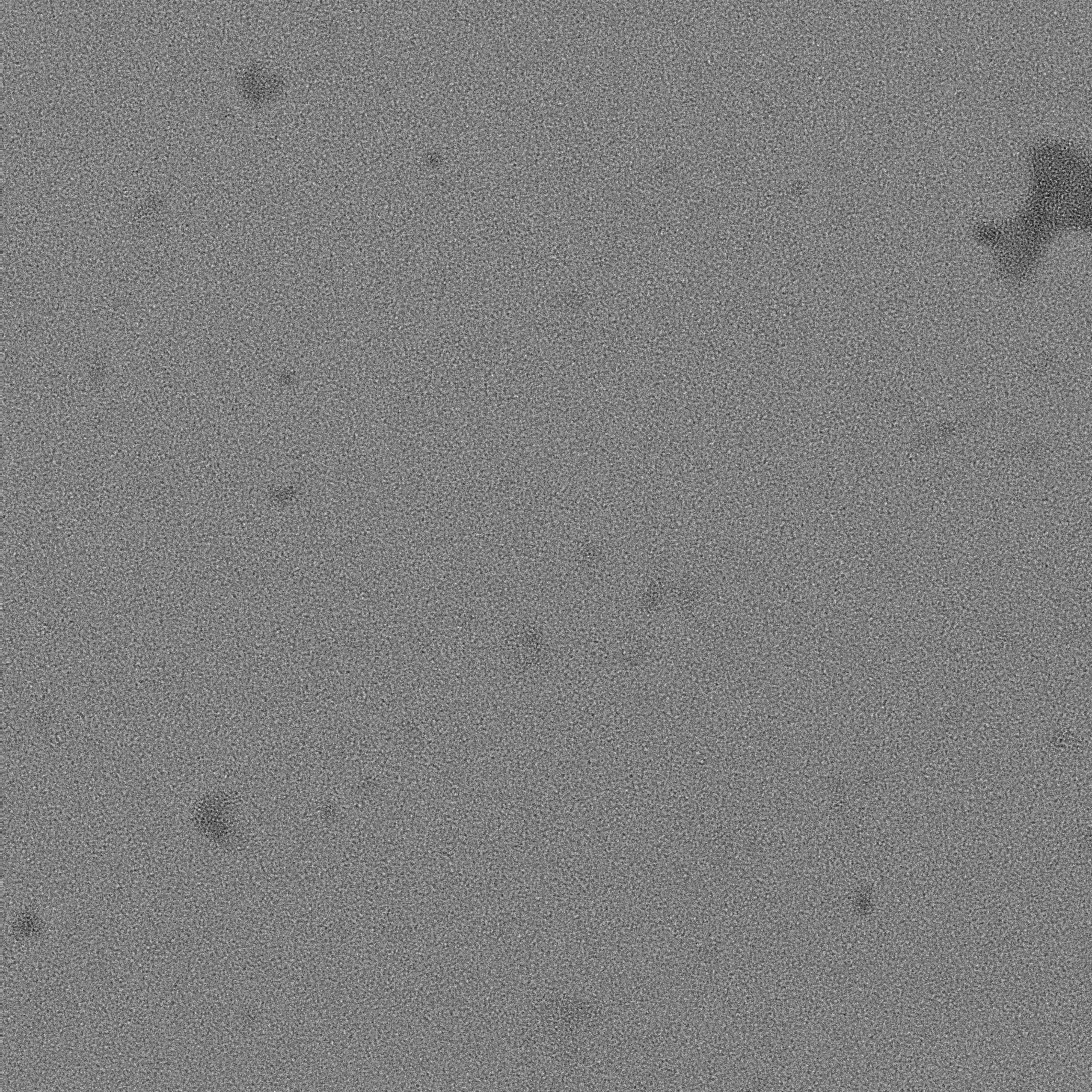

### 16E_0002_1.pdf

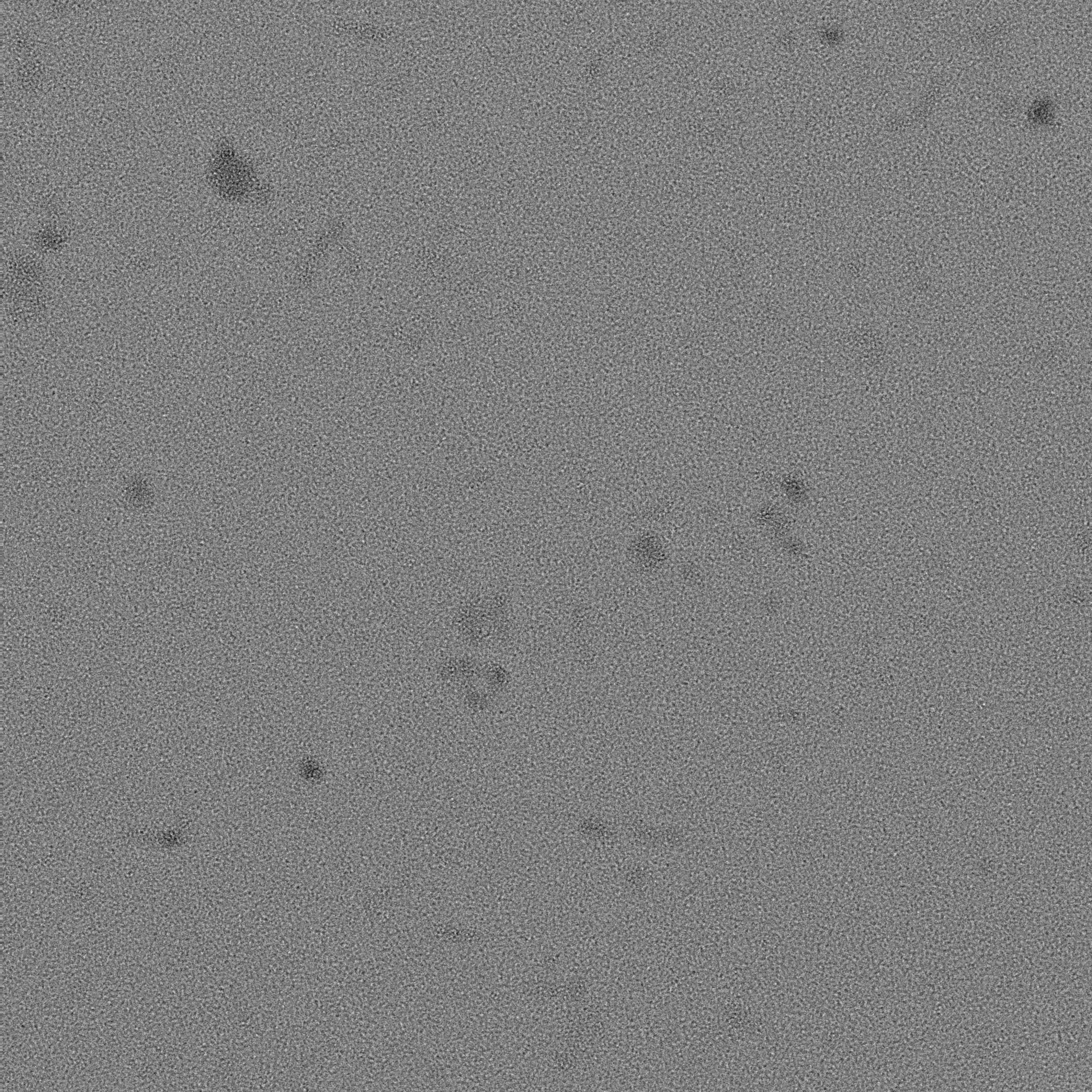

### 16E_0003_1.pdf

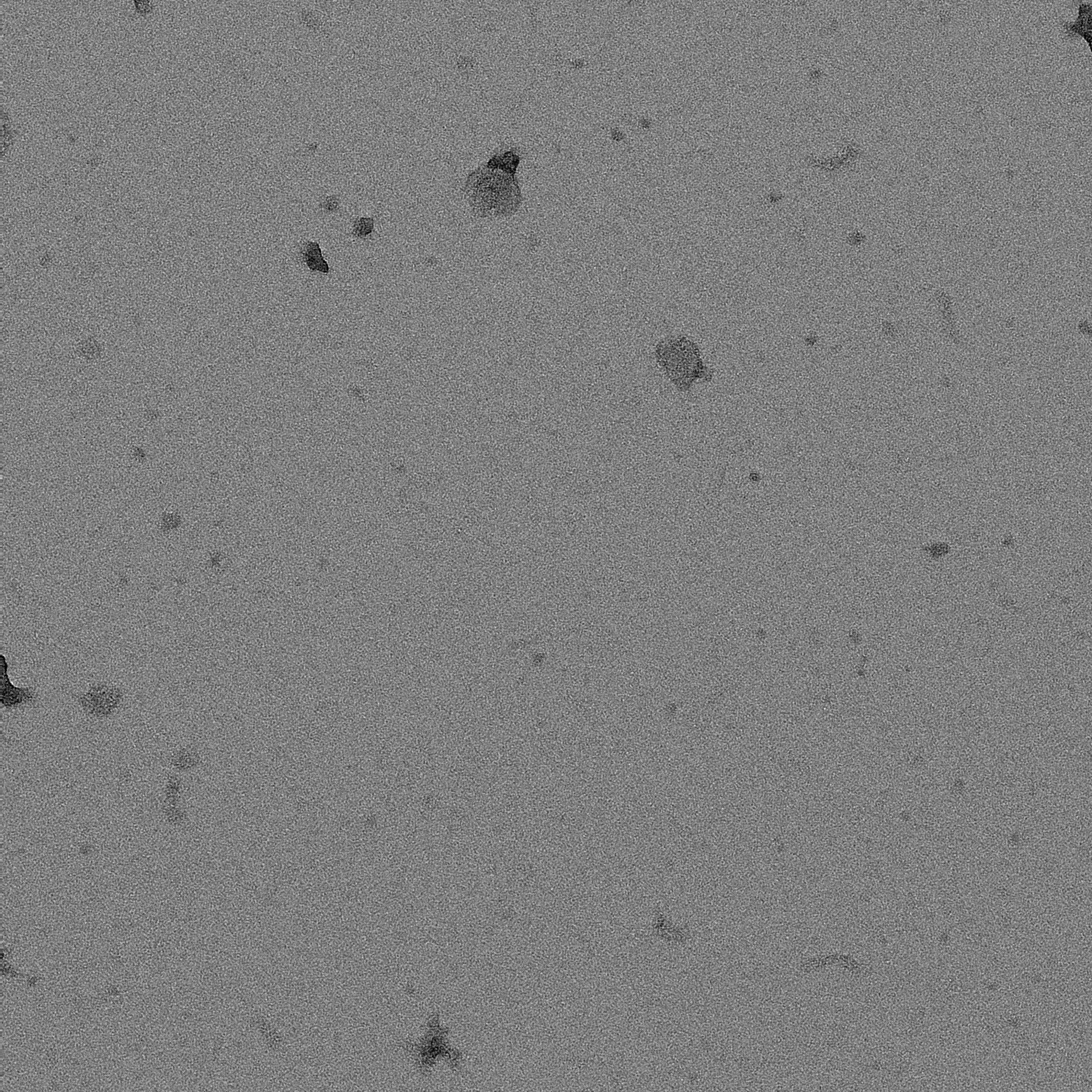

### 16E_0004_1.pdf

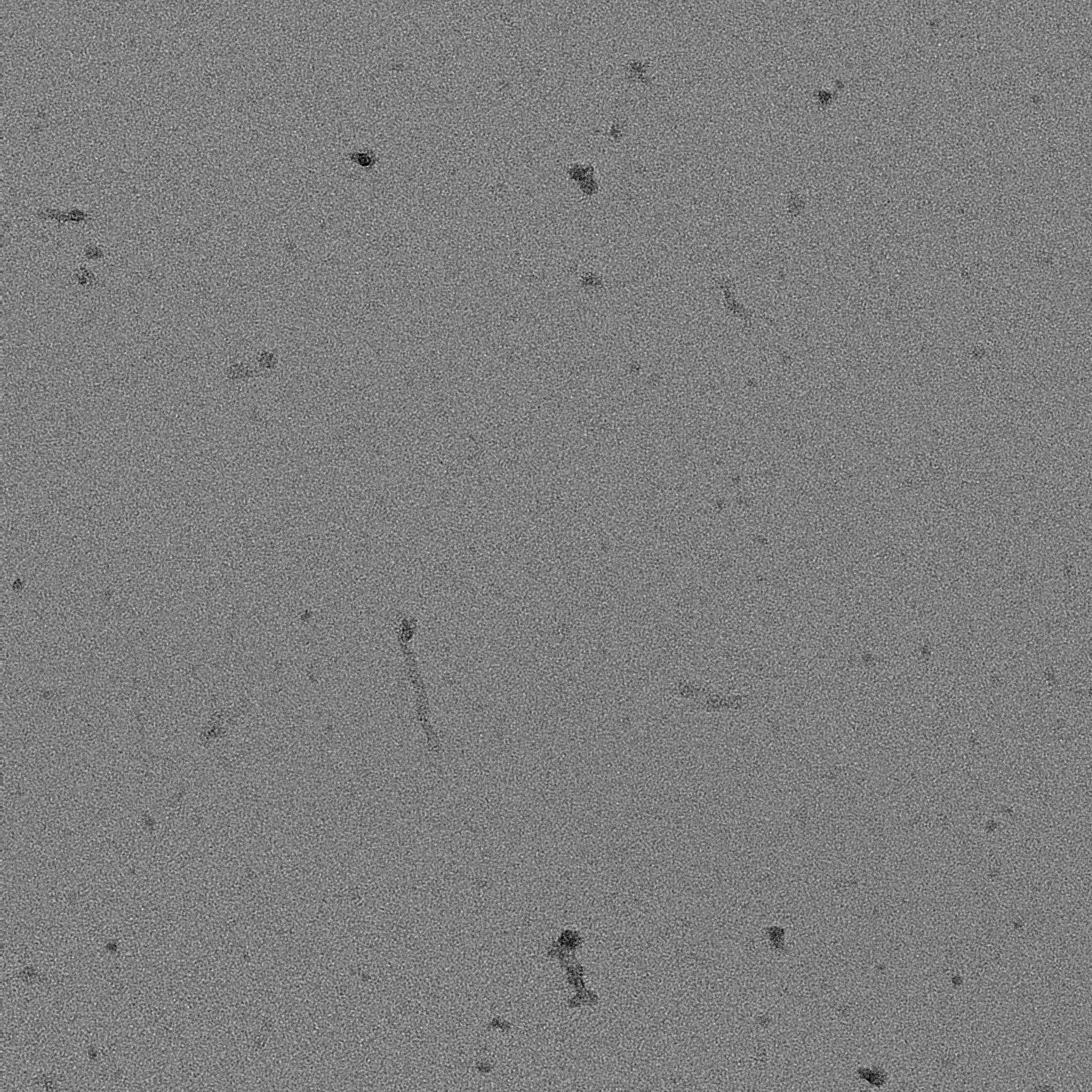

### 16E_0005_1.pdf

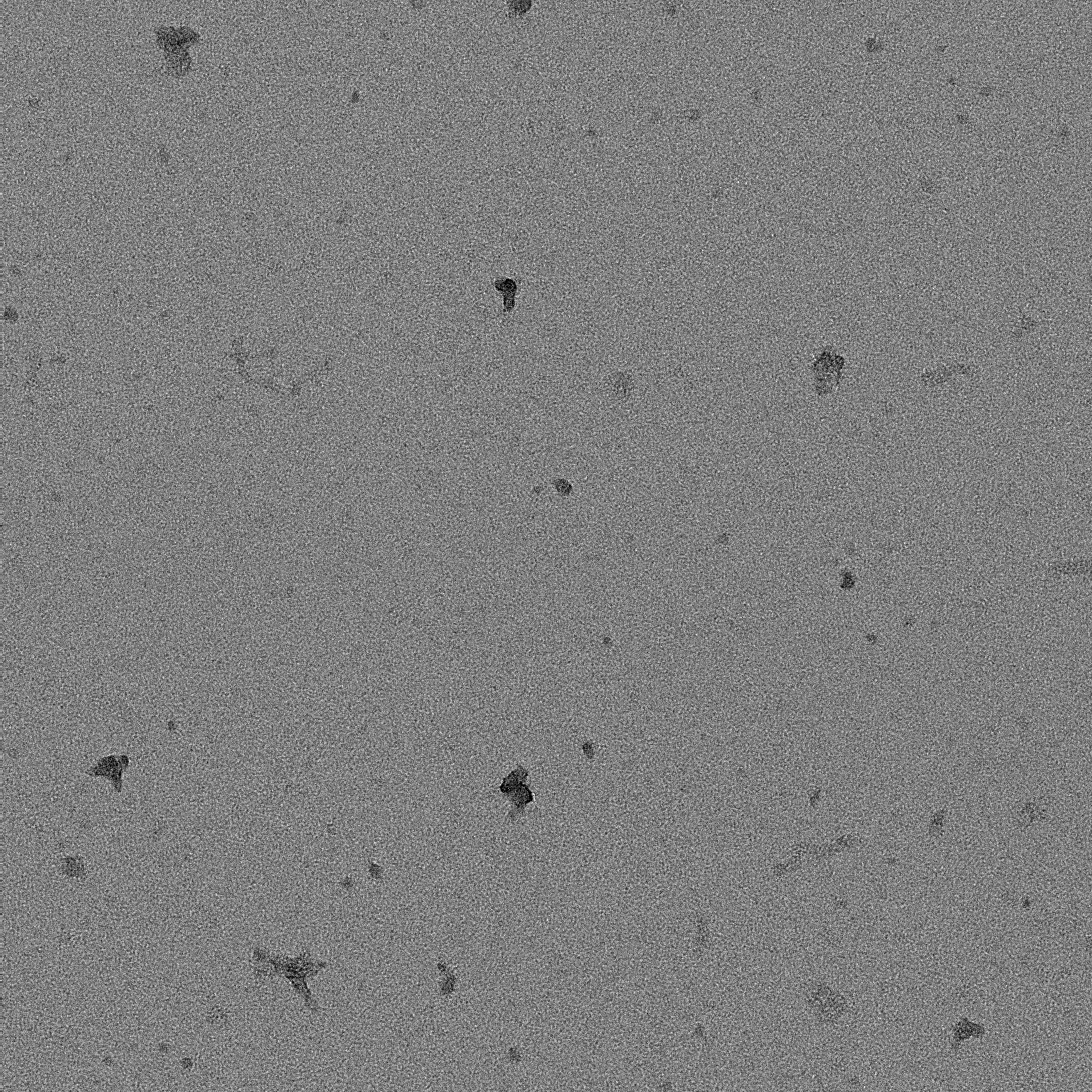

### 16E_0006_1.pdf

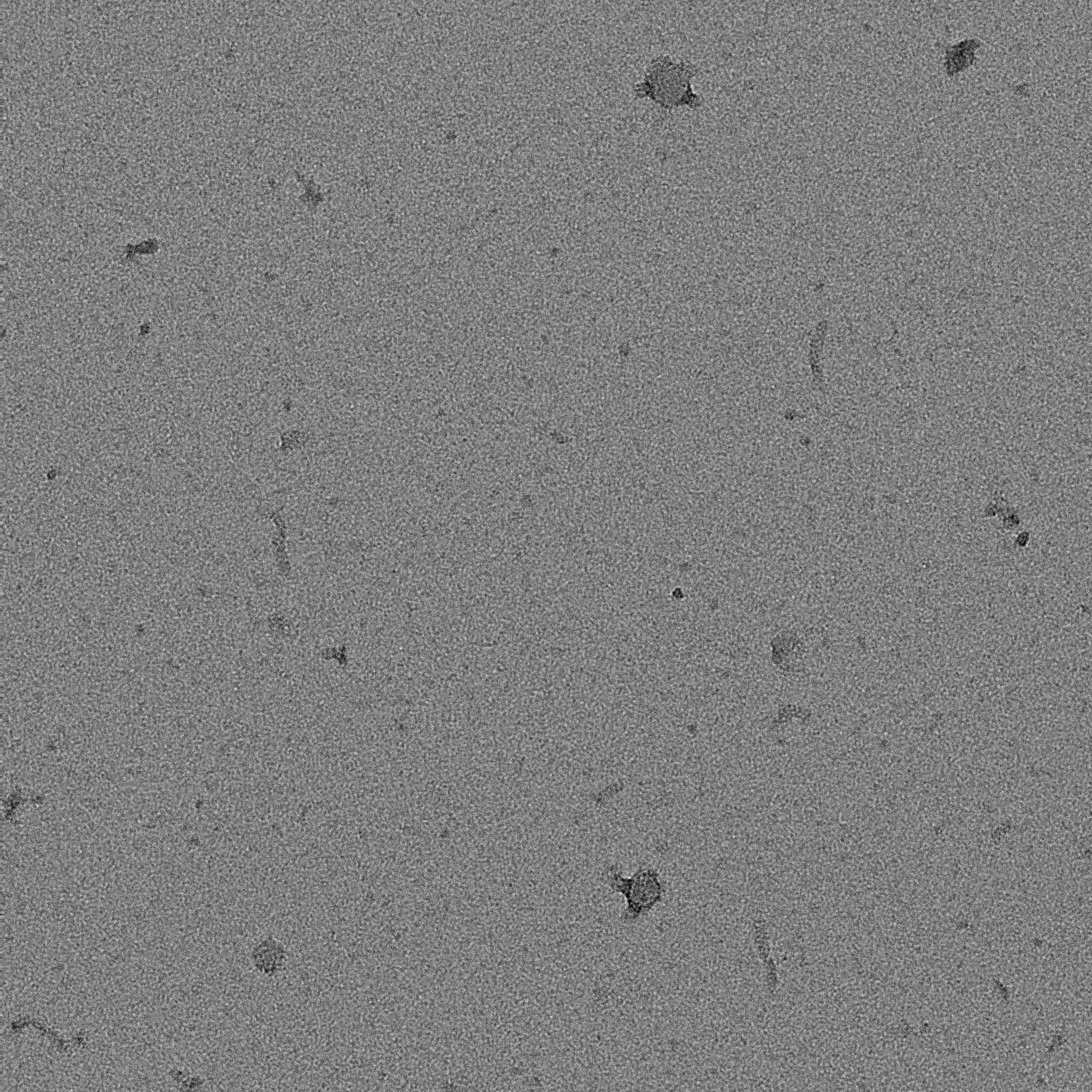

### 16E_0007_1.pdf

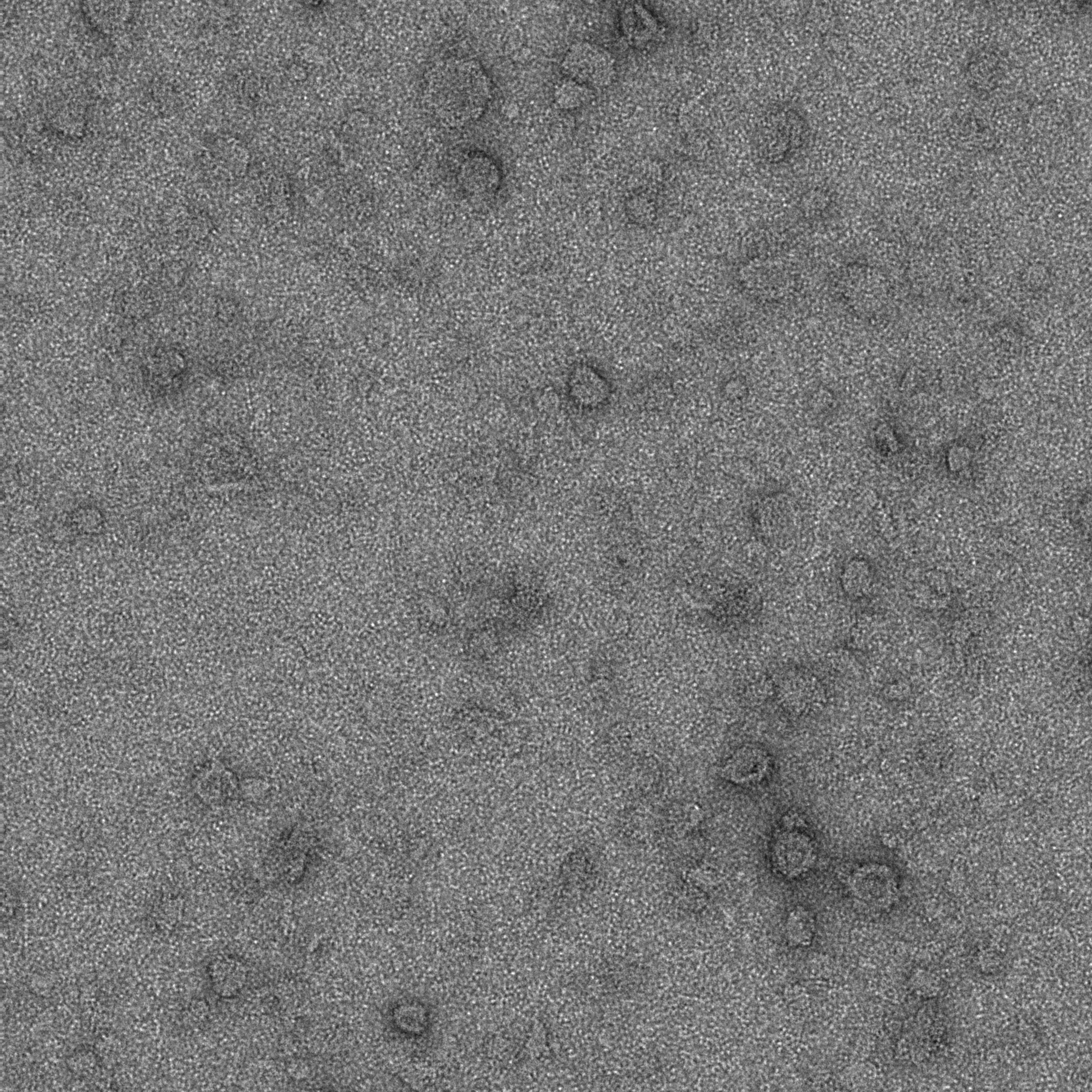

### S3 EL2 overnight_0000_1.pdf

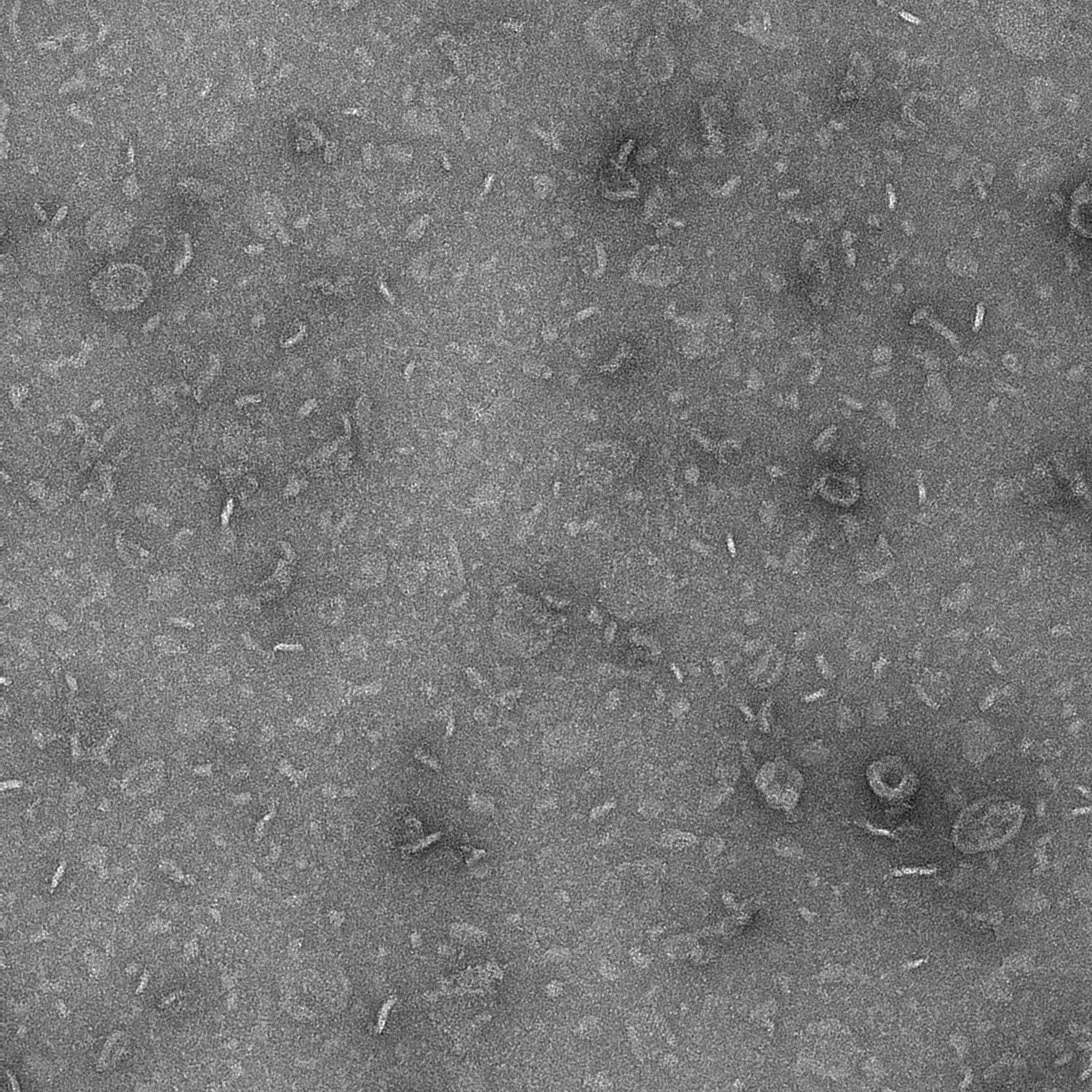

### S3 EL2 overnight_0001_1.pdf

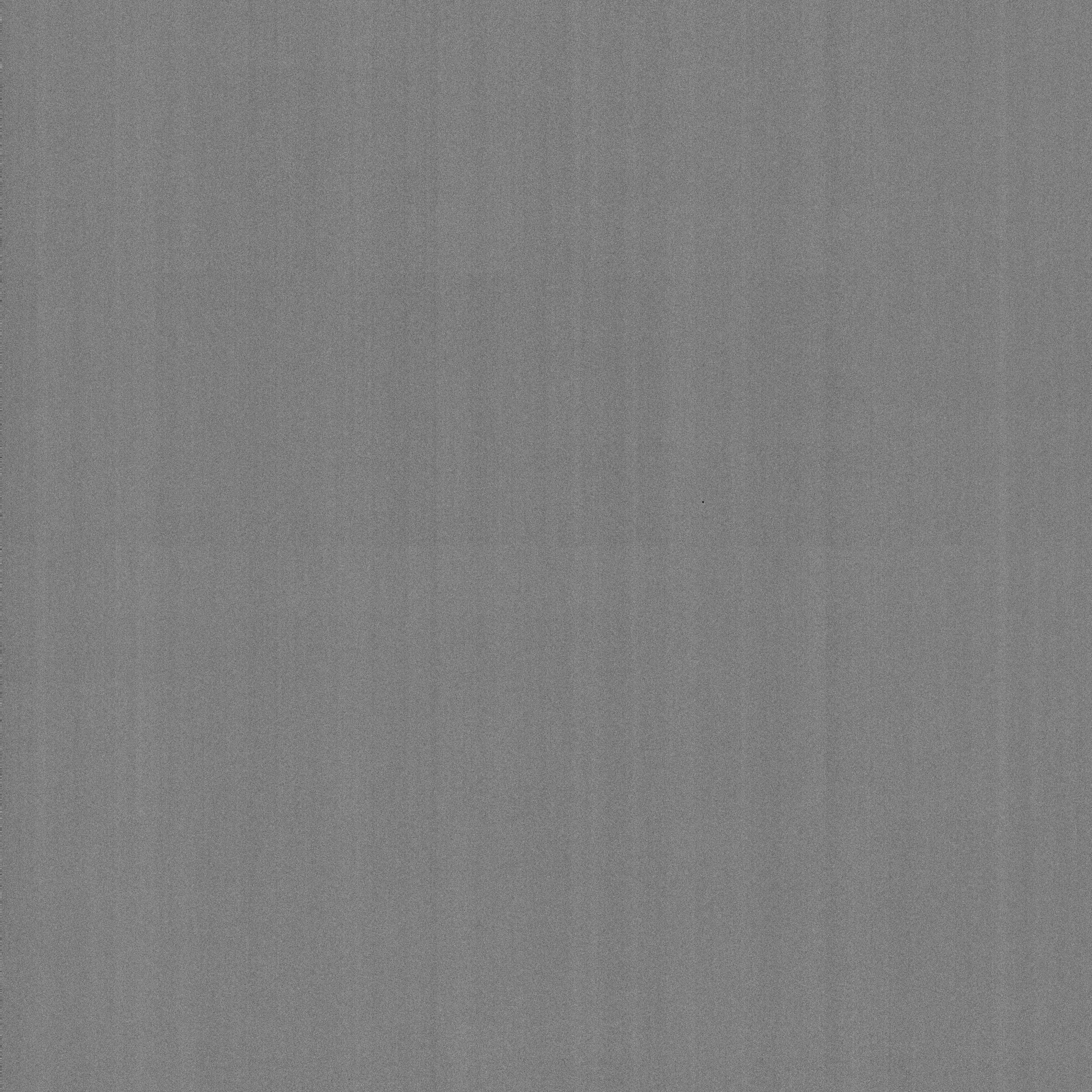

### S3 EL2 overnight_0002_1.pdf

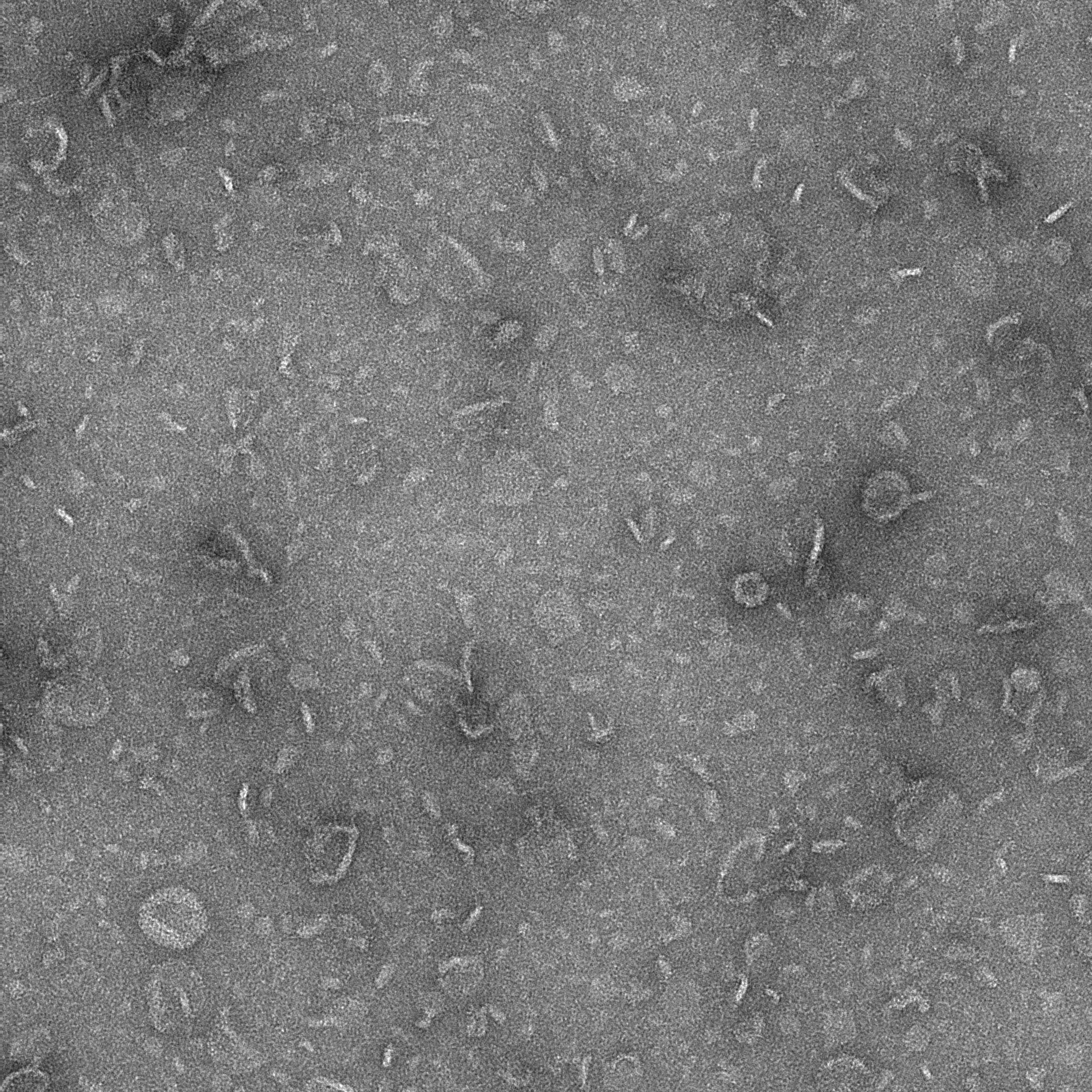

### S3 EL2 overnight_0003_1.pdf

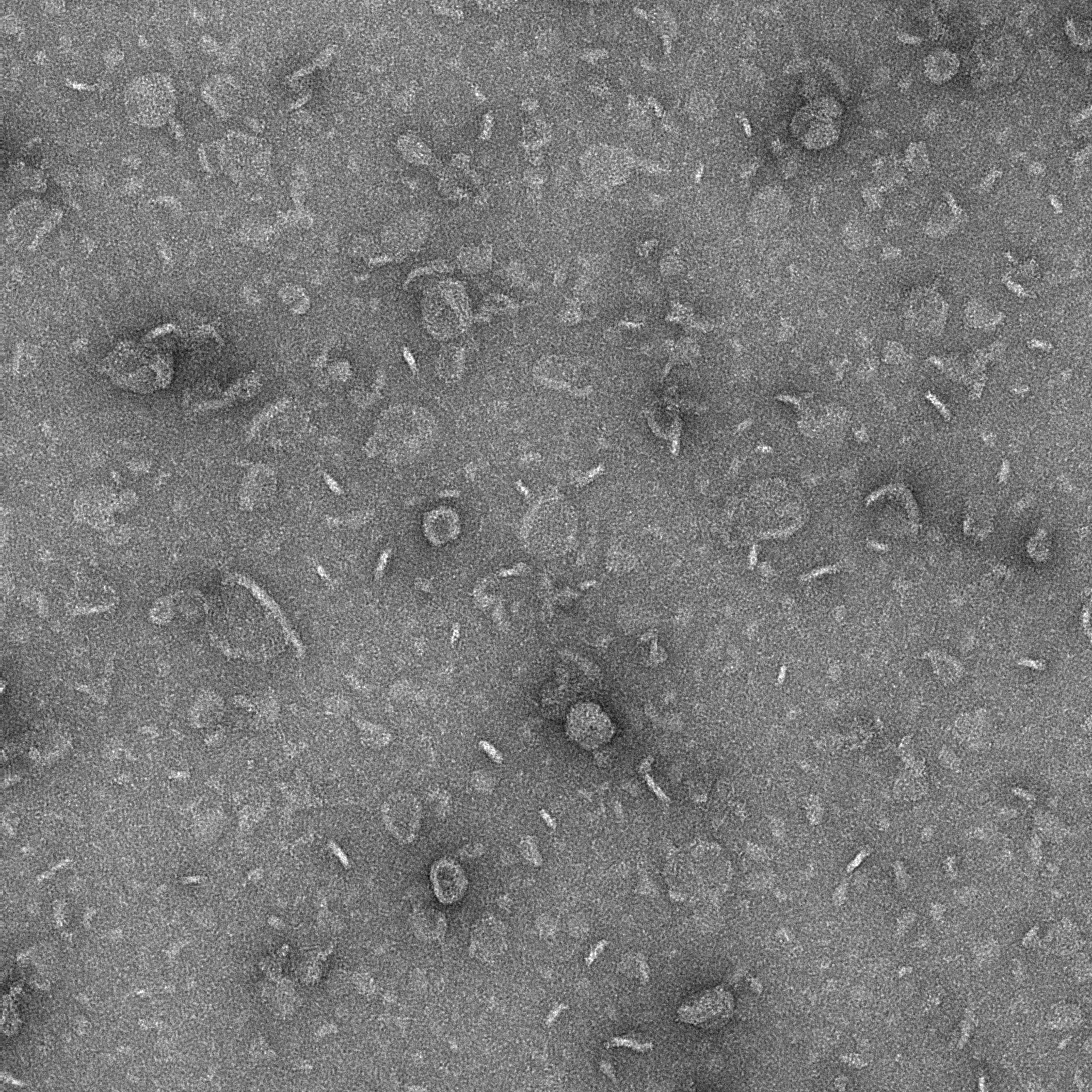

### S3 EL2 overnight_0004_1.pdf

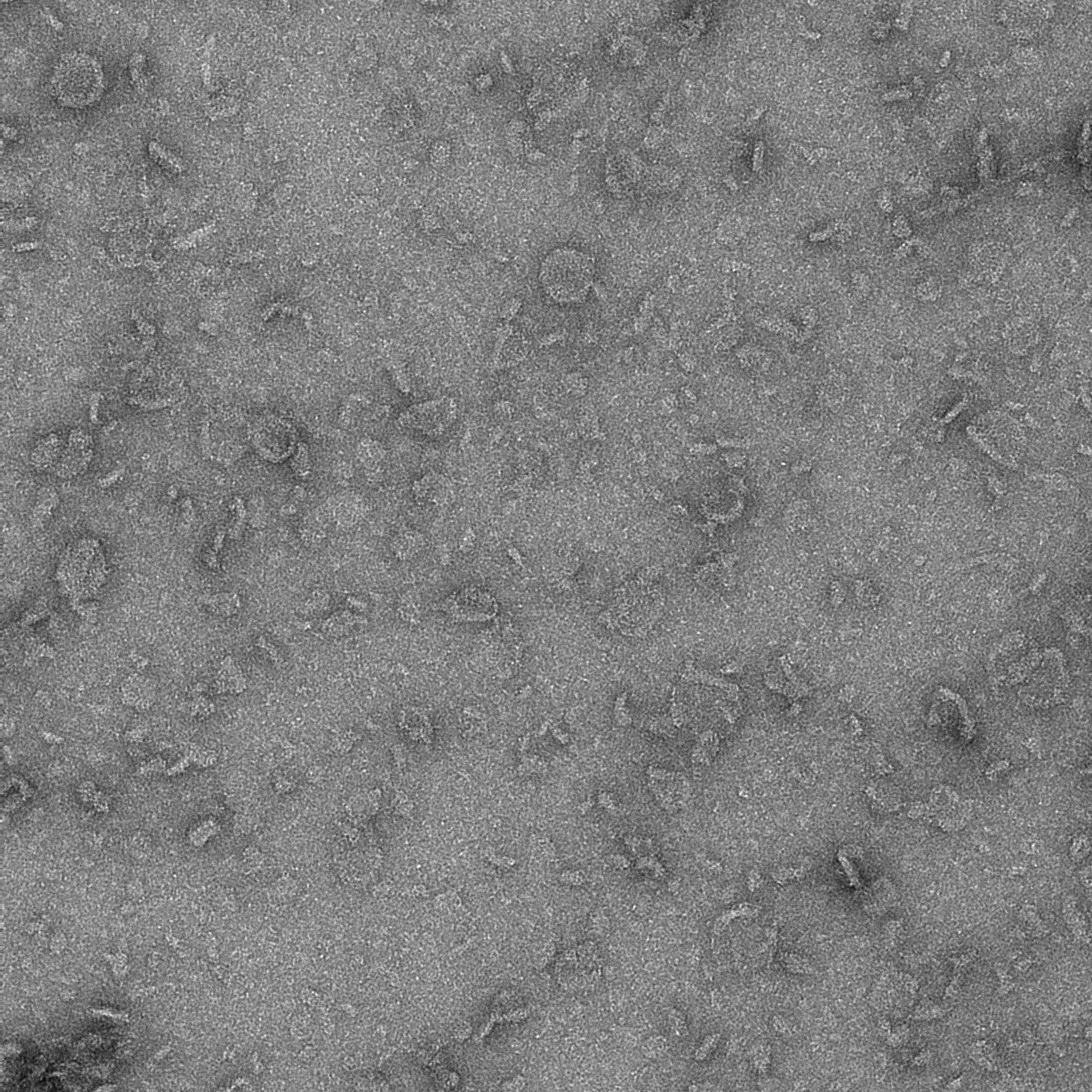

### S3 EL2 overnight_0005_1.pdf

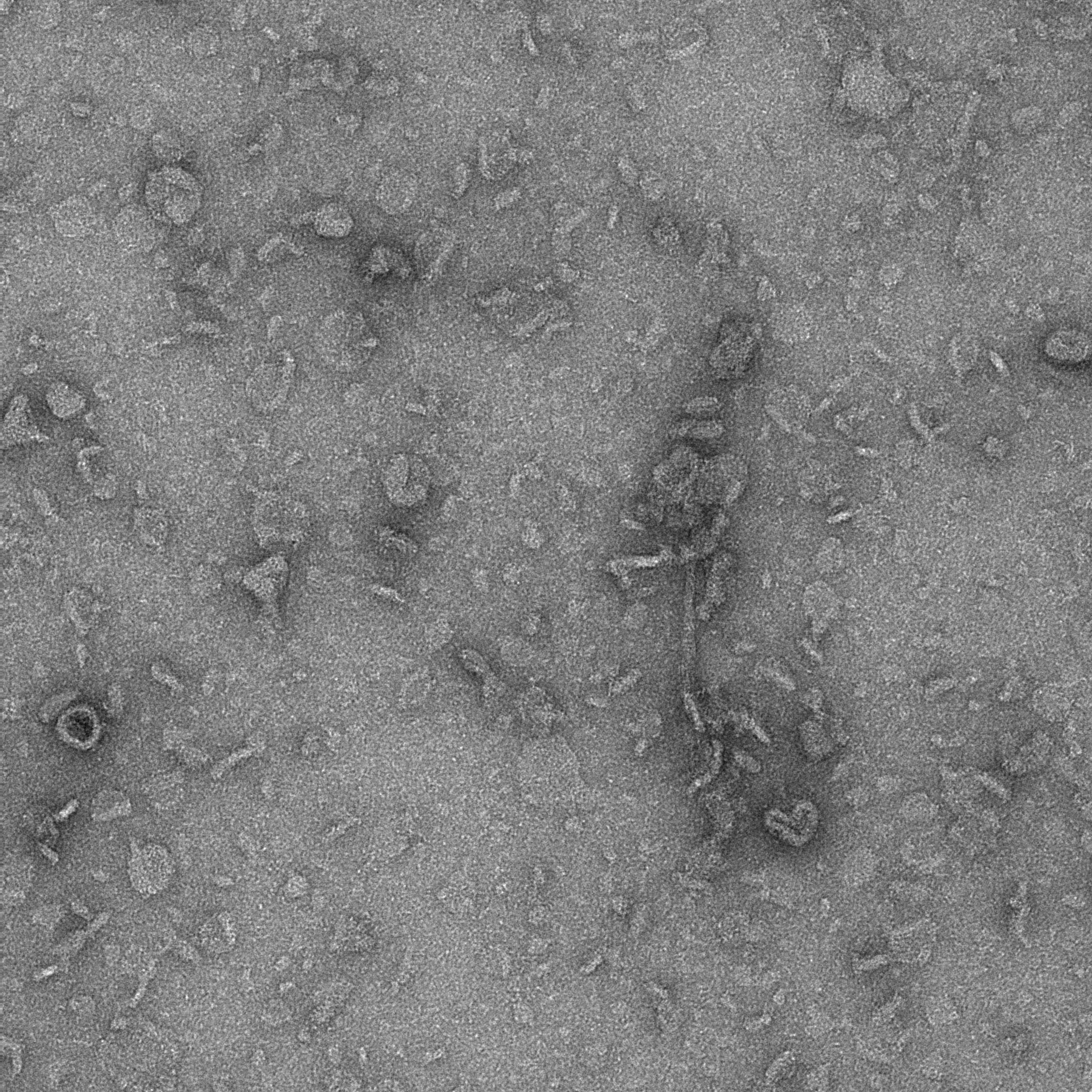

### S3 EL2 overnight_0006_1.pdf

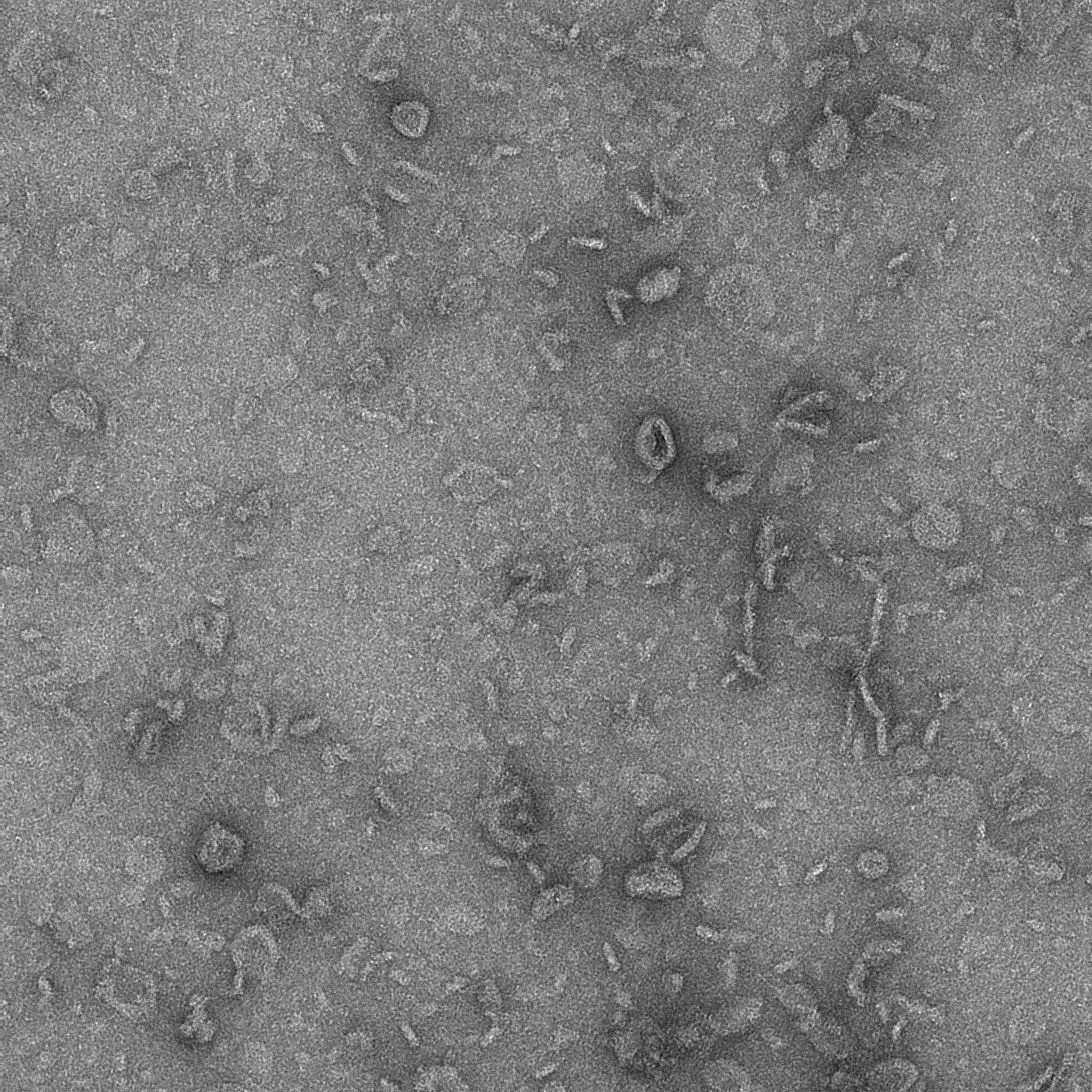

### S3 EL2 overnight_0007_1.pdf

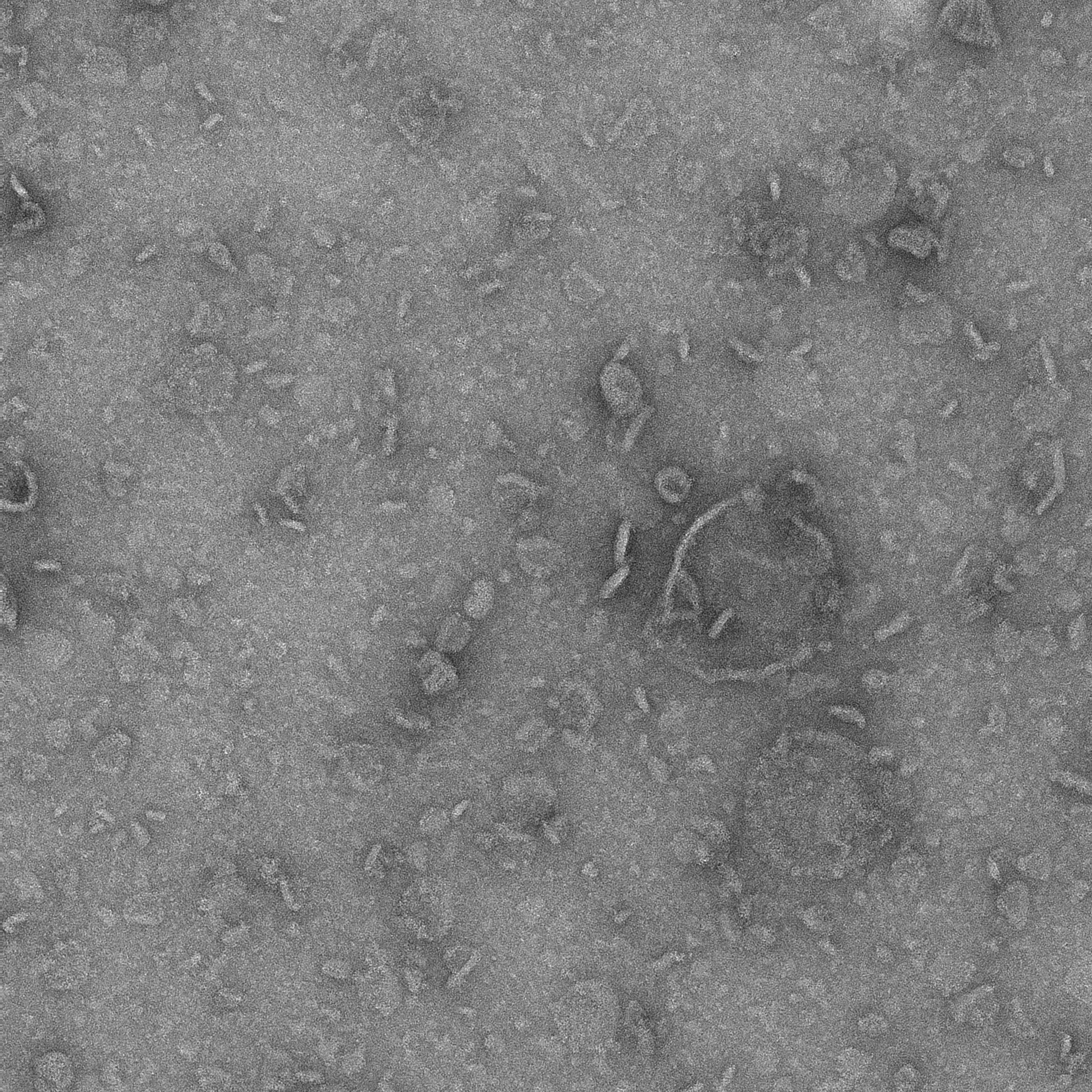

### S3 EL2 overnight_0008_1.pdf

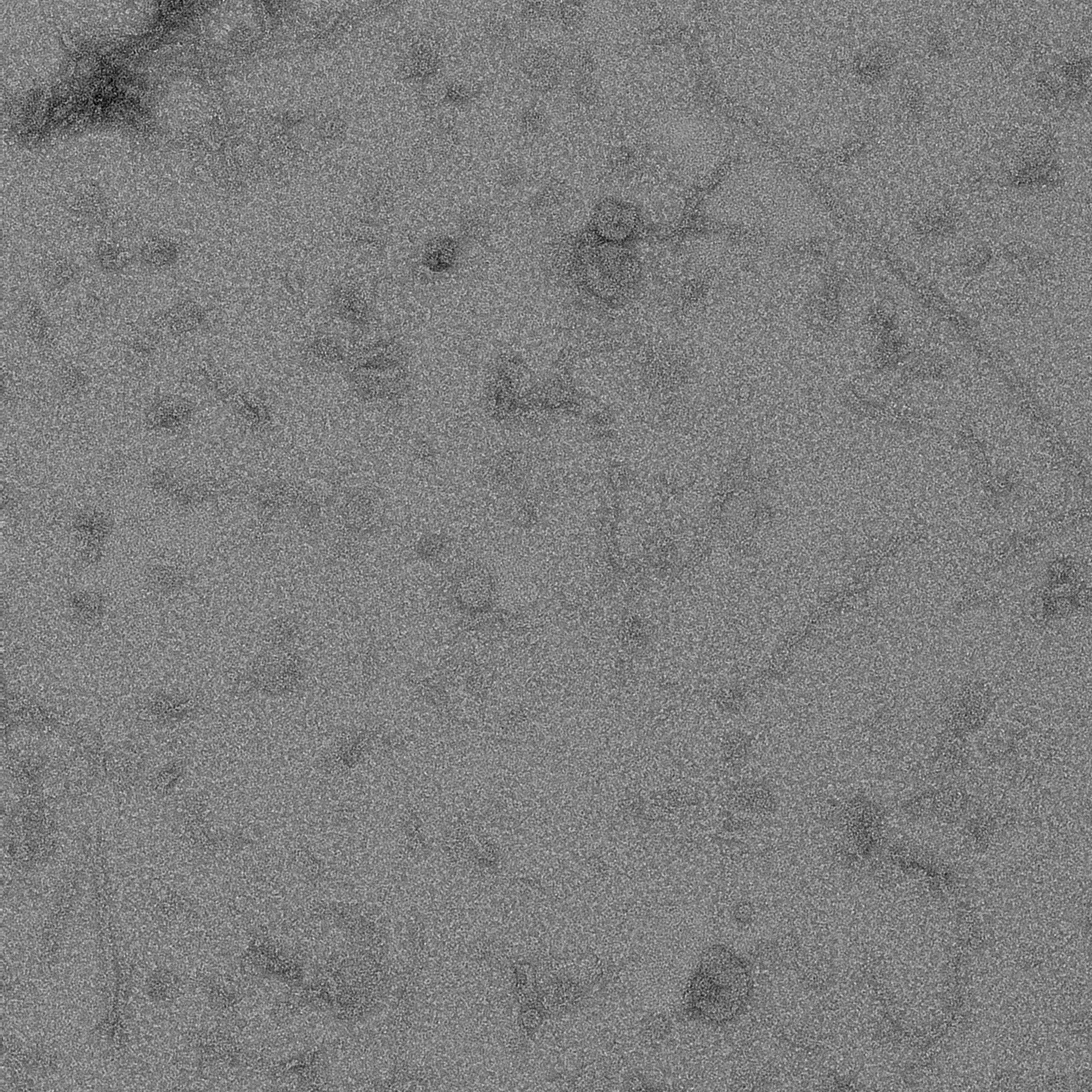

### S3 EL2 overnight_0009_1.pdf

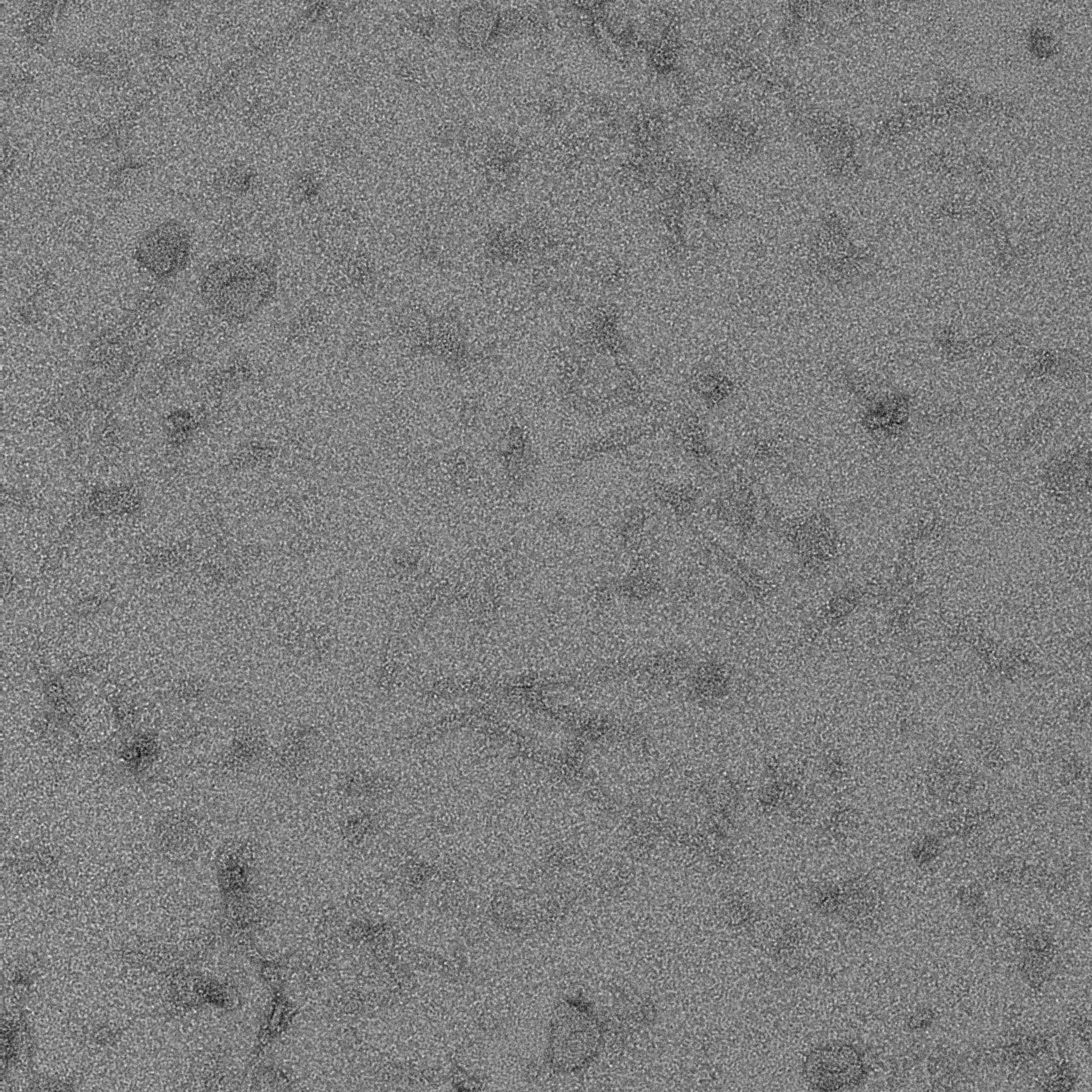

### S3 EL2 overnight_0010_1.pdf

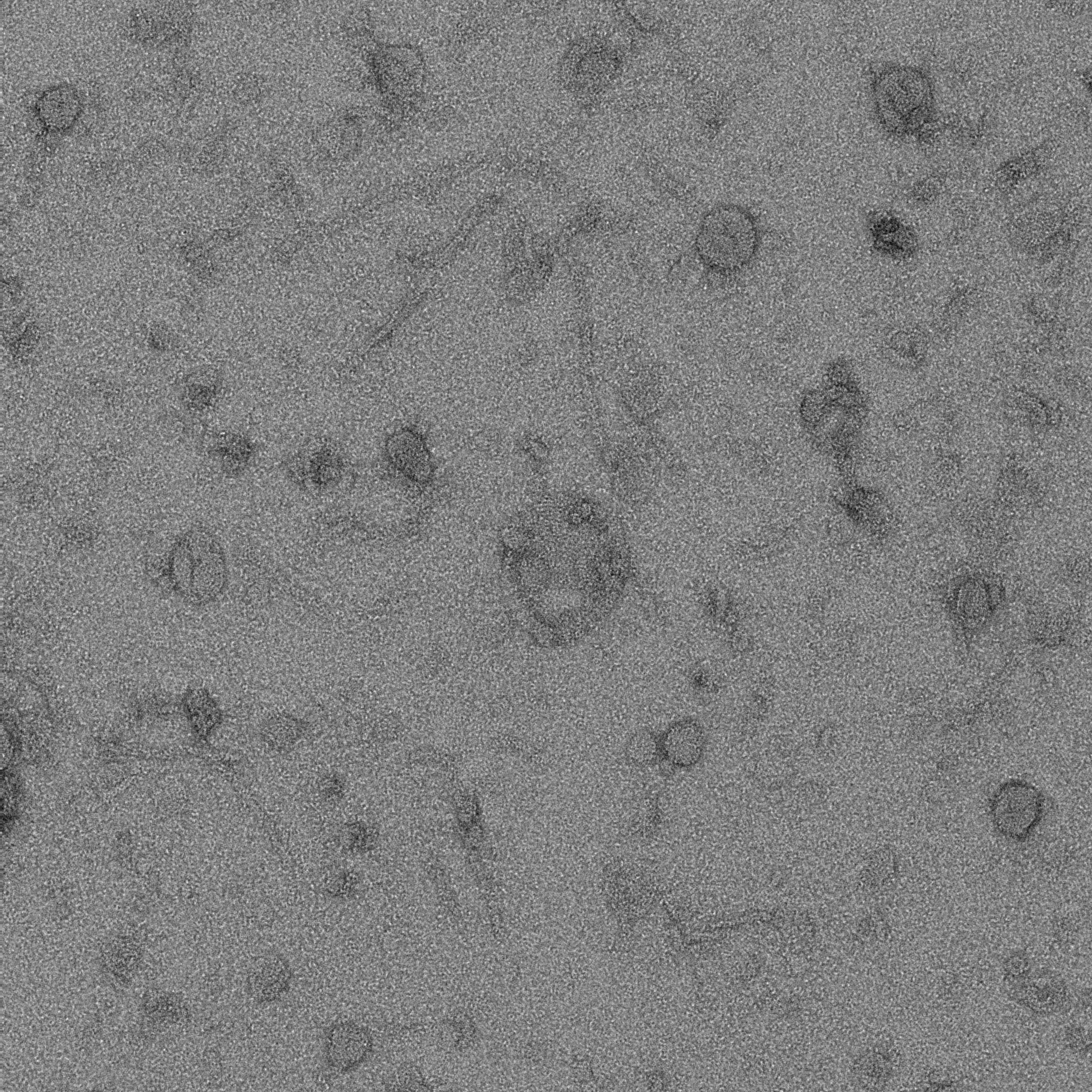

### S3 EL2 overnight_0011_1.pdf

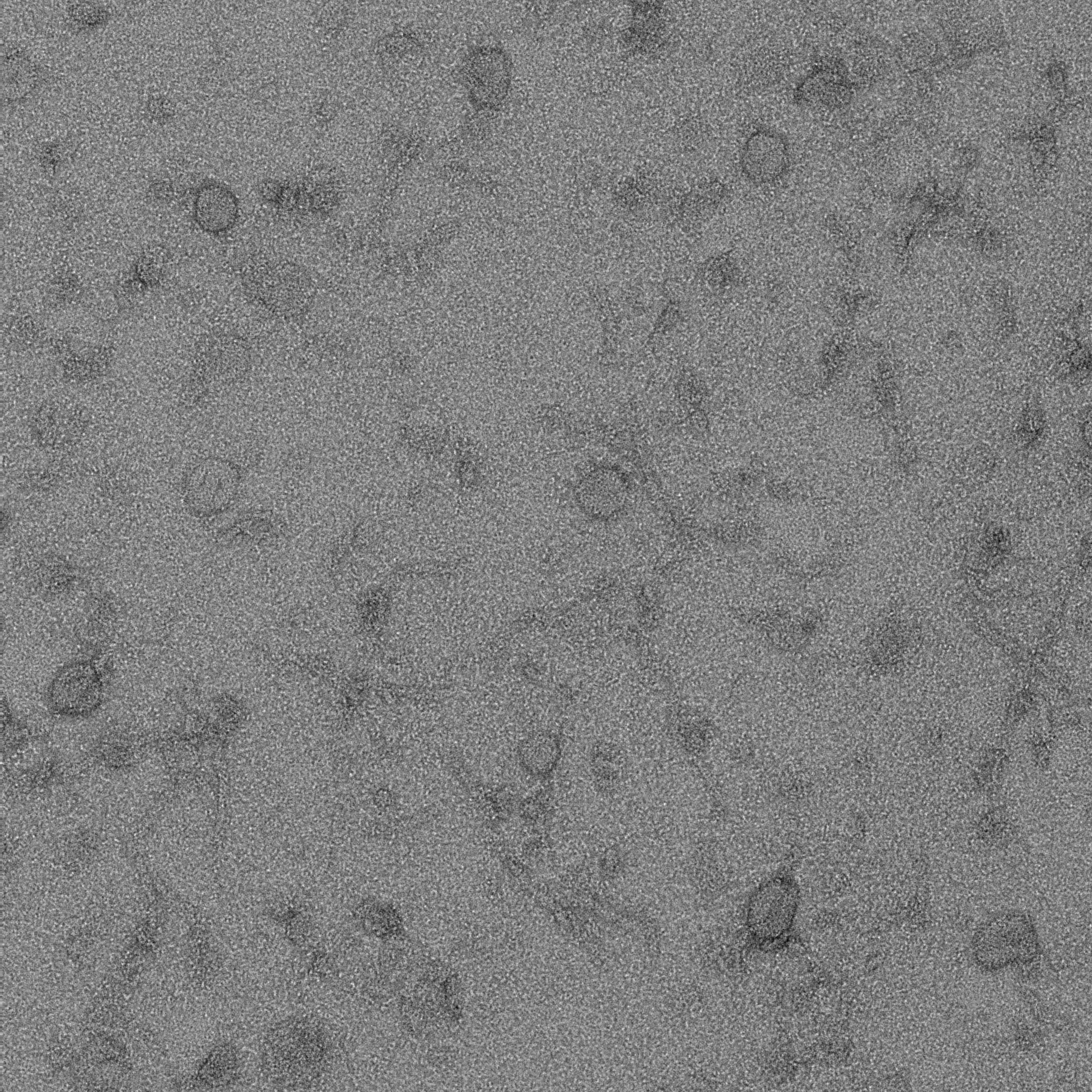

### S3 EL2 overnight_0012_1.pdf

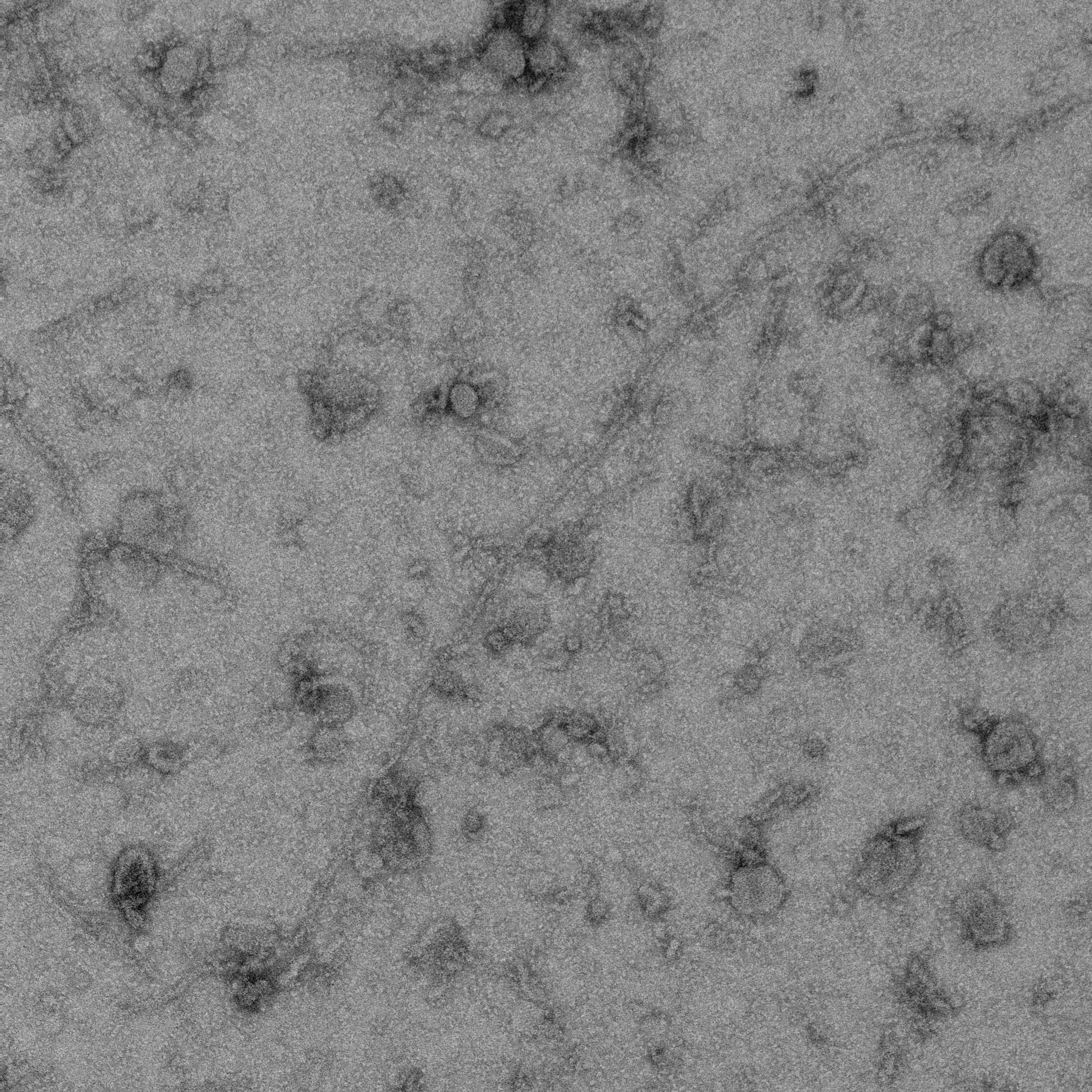

### S3 EL2 overnight_0013_1.pdf

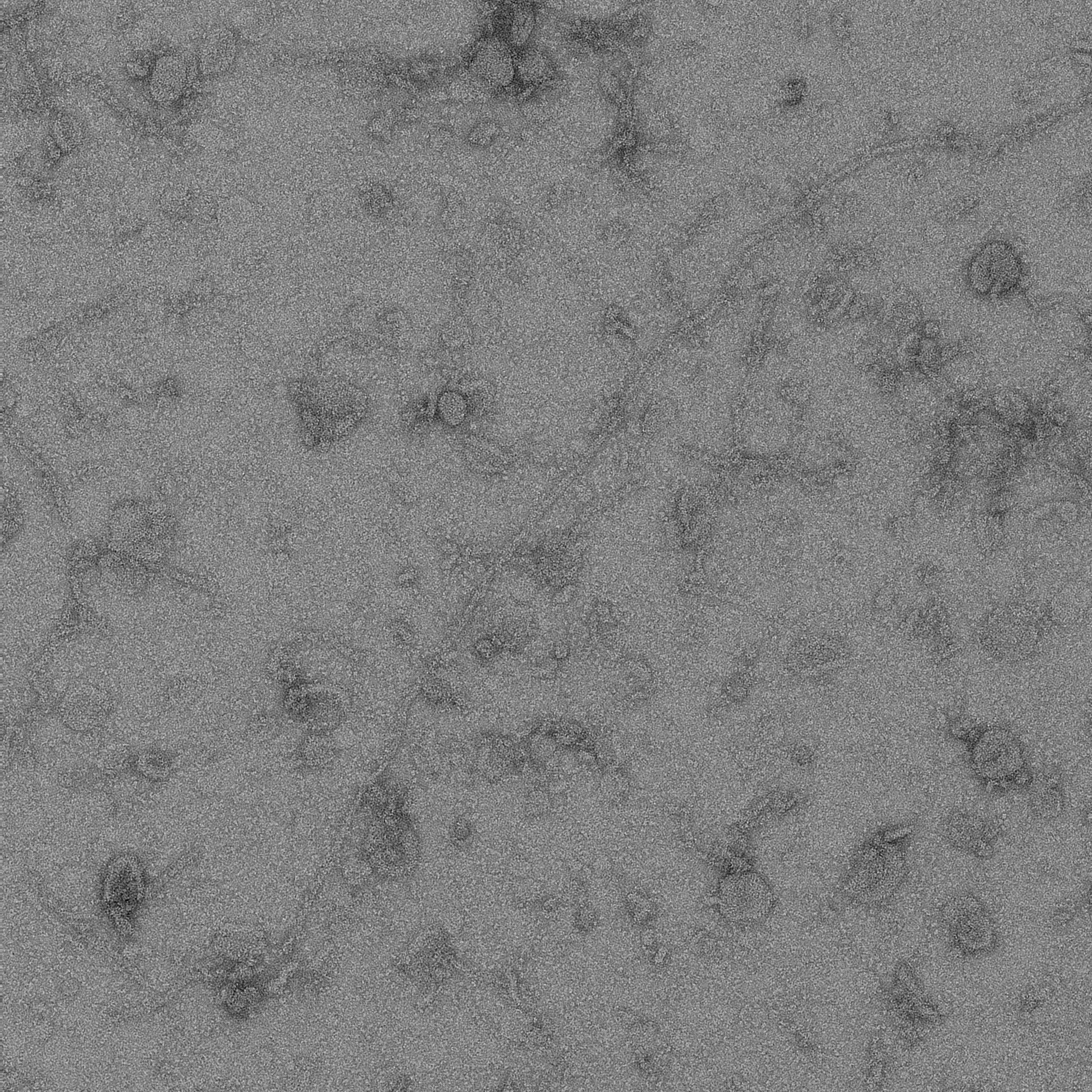

### S3 EL2 overnight_0014_1.pdf

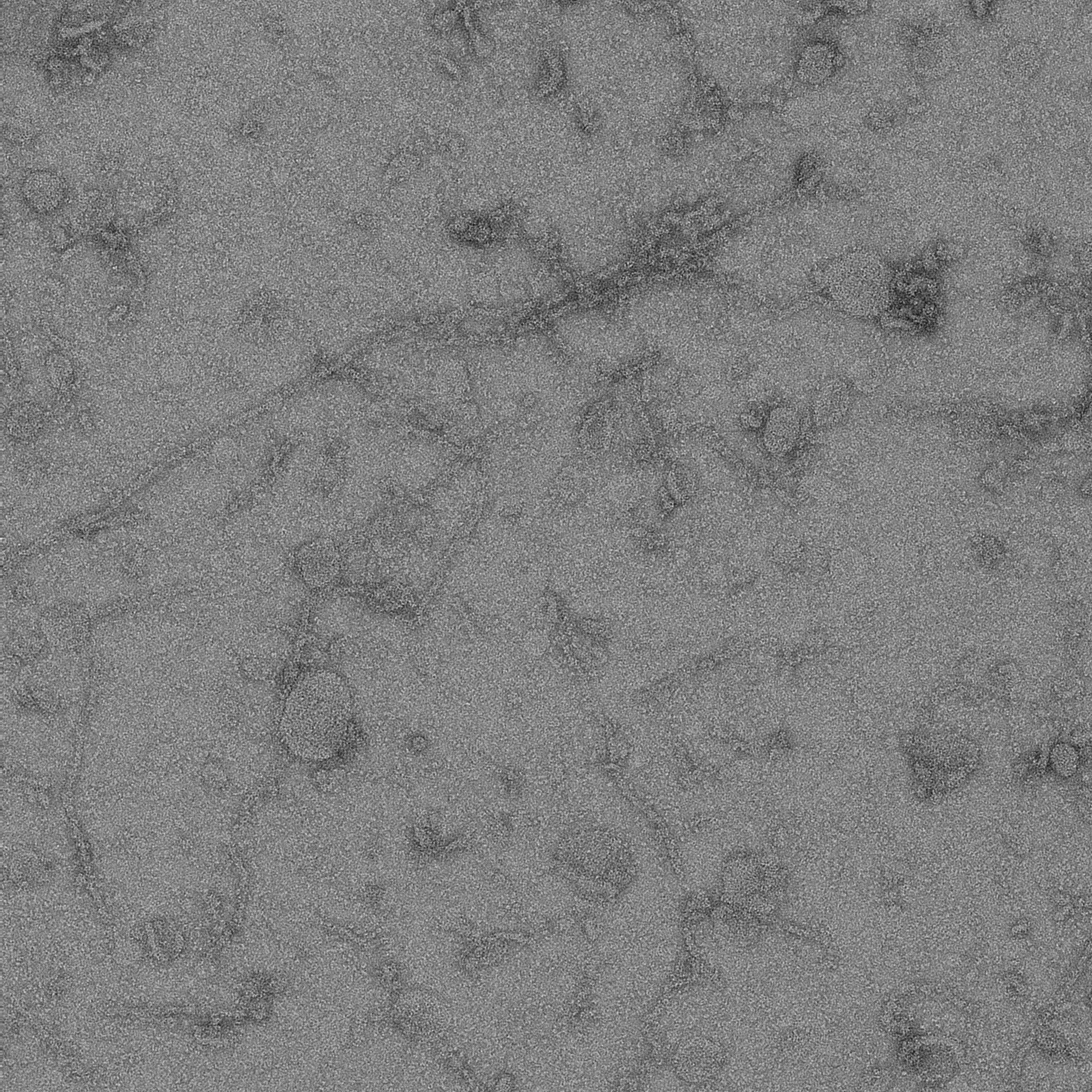

### S3 EL2 overnight_0015_1.pdf

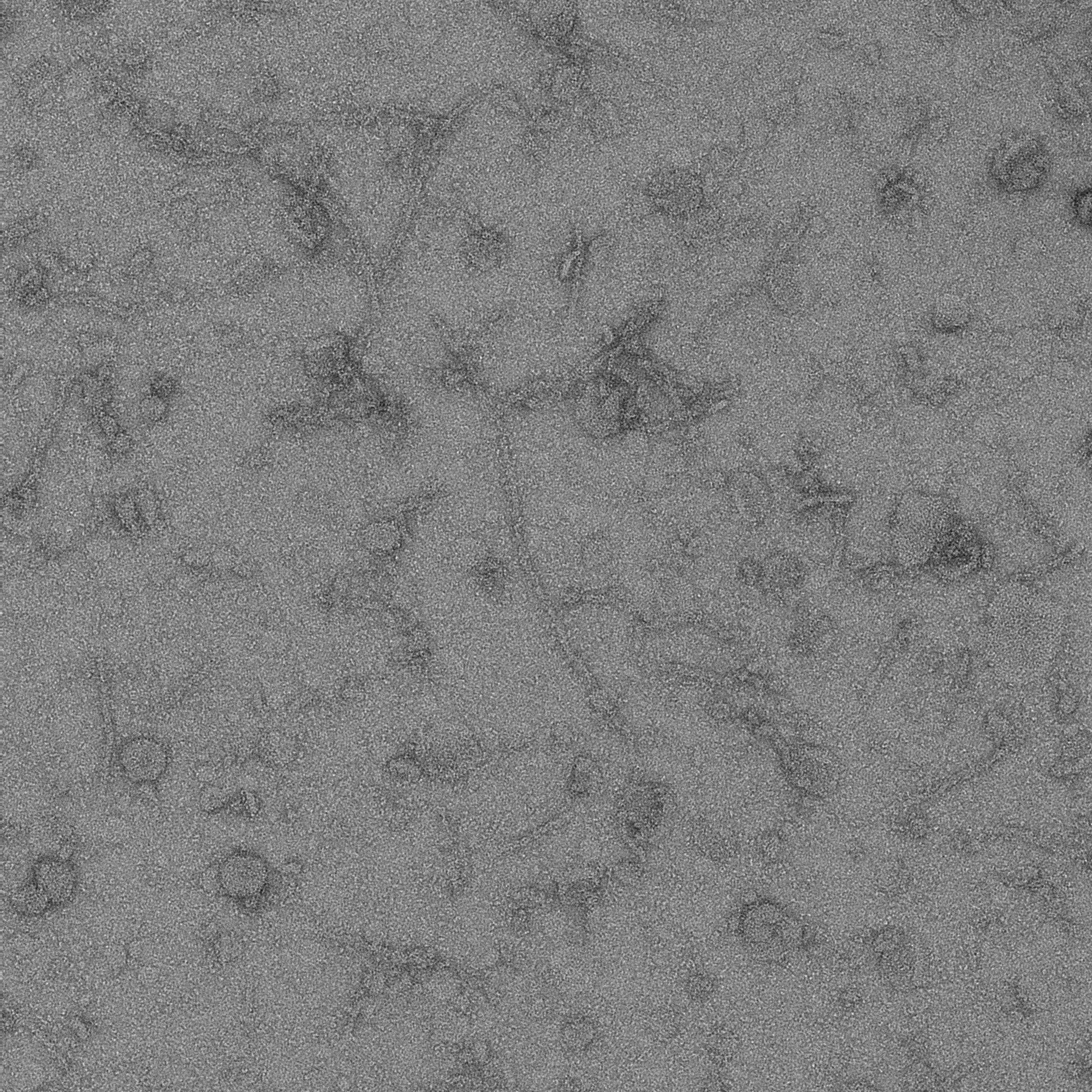

### S3 EL2 overnight_0016_1.pdf

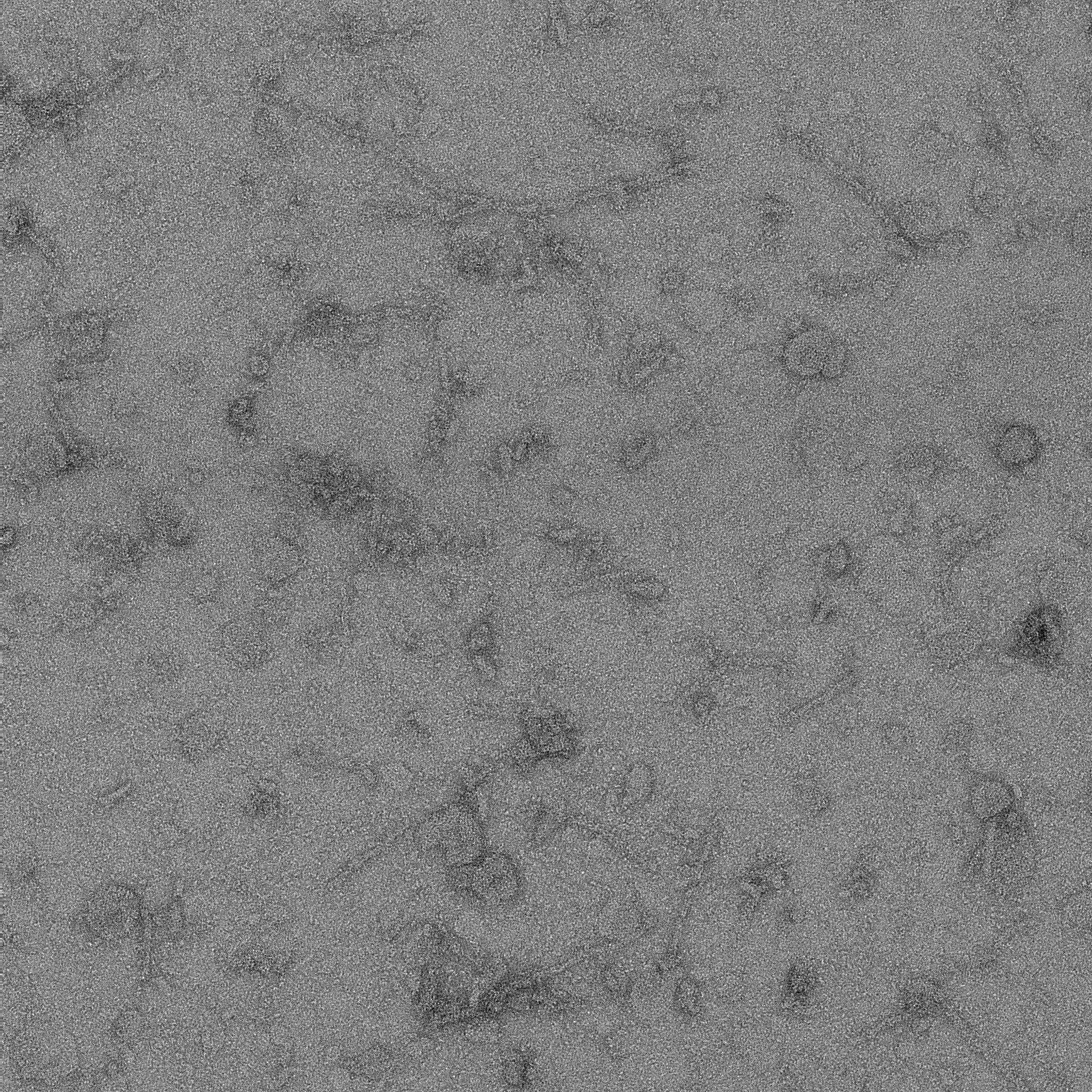

### S3 EL2 overnight_0017_1.pdf

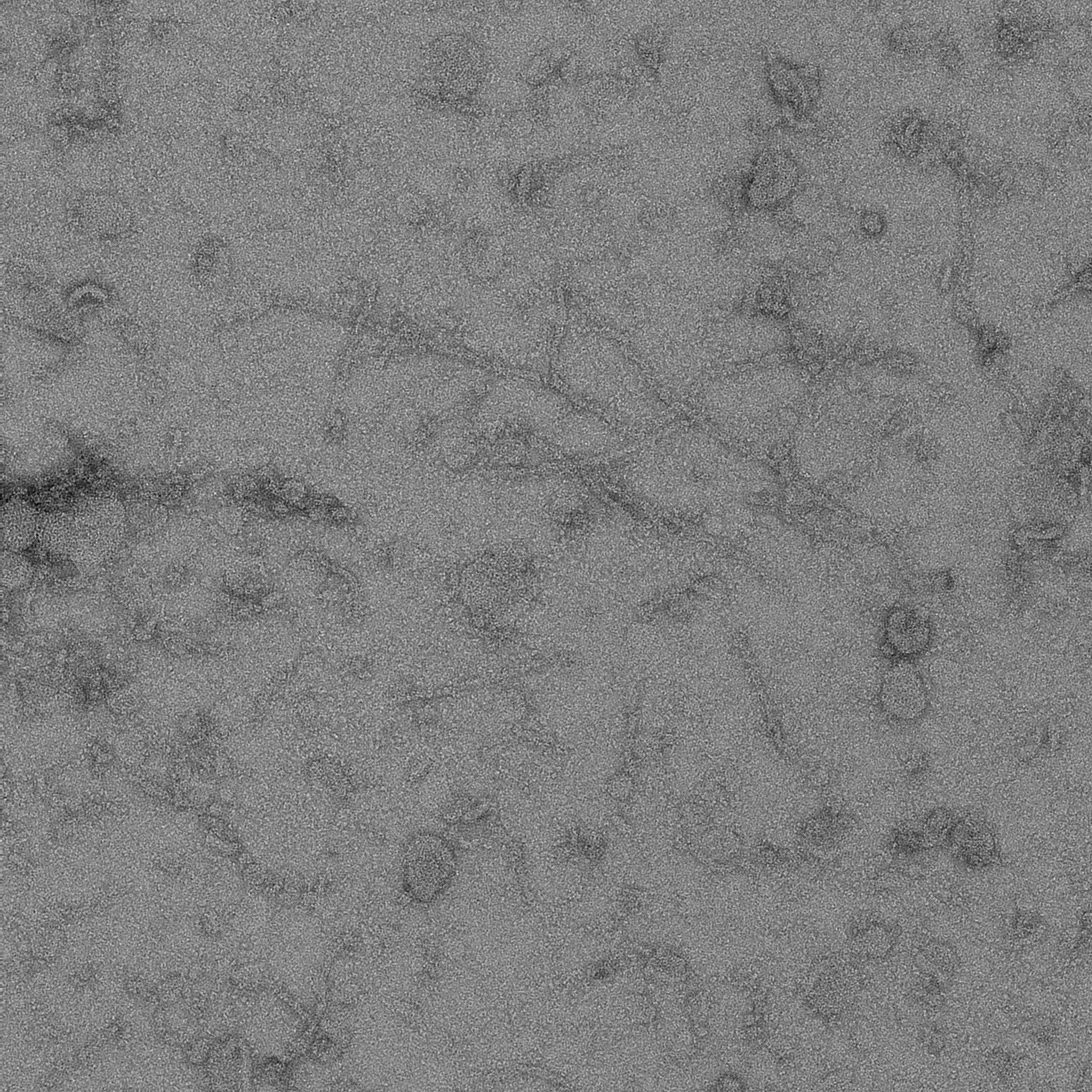
